# Atypical chemokine receptor 3 regulates synaptic removal in disease astrocytes

**DOI:** 10.1101/2024.07.03.601867

**Authors:** V Giusti, J Park, E Giusto, L Masatti, P Ramos-Gonzalez, L Iovino, M Sandre, A Emmi, G Kaur, E Coletto, Z. Van Acker, W. Annaert, E Calura, F Petrelli, A Porzionato, R. De Caro, F Cavaliere, WS Chung, L Civiero

## Abstract

Astrocytes participate in the clearance of obsolete or unwanted neuronal synapses. However, the molecular machinery involved in recognizing these synapses remains unclear, particularly under pathological conditions. Here, we investigated the phagocytic process of astrocytes through individual gene silencing with a druggable gene library. Our study demonstrates that astrocyte-mediated synapse engulfment is regulated by the Atypical chemokine receptor 3 (Ackr3). Mechanistically, we showed that Ackr3 recognizes phosphatidylethanolamine (PE)-bound C-X-C motif chemokine 12 (CXCL12) at synaptic terminals both in vitro and in vivo, thus serving as a novel marker of synaptic dysfunction. Notably, both the receptor and its ligand are upregulated in post-mortem human Alzheimer’s disease (AD) brains, and AD mouse models. Downregulation of the Ackr3 in AD mice significantly diminishes astrocyte-mediated synaptic elimination, and rescues pathological phenotypes, including synapse loss and cognitive impairment. Overall, this work unveils a novel, possibly targetable mechanism of astrocyte-mediated synaptic engulfment implicated in neurodegenerative disease.

## Introduction

Synapses require constant and multimodal surveillance by glial cells ^1^. Appropriate clearance of extraordinary and non-functional synapses is necessary for the homeostasis of the central nervous system (CNS) ^2,3^.

Similar to microglia, astrocytes engulf parts of living and dying cells, including synapses and axons both in physiological and pathological context context ^4–12^.

Multiple EGF-like domains 10 (Megf10) receptor and MER proto-oncogene tyrosine kinase (Mertk) are involved in synaptic removal by astrocytes, possibly forming a molecular complex together with other proteins ^1,13^. Interestingly, the process is driven by neuronal activity in the developing and adult brain ^4^. Functionally, the lack of Megf10 in astrocytes causes an accumulation of dysfunctional excitatory synapses in the hippocampus thus impacting long term synaptic plasticity and memories^11^. Megf10-mediated synaptic engulfment is an extremely conserved mechanism, with several examples being documented in Drosophila (Draper) and *C. elegans* (CED-1) ^14,15^. Recently, the ATP-binding cassette transporter A1 (Abca1) and the engulfment adapter phosphotyrosine-binding domain containing 1 (Gulp1) together with Megf10, have been identified as the responsible molecules for the enhanced astrocytic phagocytosis of synapses in a mouse model for brain ischemia^7^. Phosphatidylserines (PS), exposed by dying cells, are proposed to be recognized by Megf10 on the cell surface possibly through the involvement of bridging molecules whose identity is still unclear (e.g., C1q) ^4,16^. Although *via* still unknown pathways, hippocampal astrocytes contact and extensively eliminate synapses by recognizing C1q molecules in AD mouse models ^12^. In humans, AD astrocytes show enhanced phagocytosis of synapses through the recognition of Milk fat globule-EGF factor 8 protein (Mgf-E8), a PS-binding molecule ^17^. Astrocytes also release ATP in an inositol 1,4,5-trisphosphate receptor type 2 (Ip3r2)-dependent manner to regulate synapse elimination ^5^. Astrocytes also cooperate with microglia to clear up unwanted material within the brain parenchyma ^8,10^. Specifically, a cross talk between astrocytes and microglia through inflammatory molecules have been reported in clearing up synapses ^6,18,19^.

Despite the recent advances in understanding the role of astrocytes in synapse elimination, key molecular components remains unidentified, including additional astrocyte-specific receptors and modulators, as well as the synaptic tags used by astrocytes to recognize synapses for clearance.

Here, we identified Atypical chemokine receptor 3 (Ackr3) as a novel receptor through which astrocytes engulf synapses under pathological conditions. Ackr3 is a C-X-C chemokine receptor for Cxcl11 and Cxcl12 ^20–22^. Although Ackr3 has been detected in the brain, specifically in neurons and astrocytes but not microglia ^21,22^, the molecular mechanism behind Ackr3-mediated synaptic pruning remains completely unknown. Through our work, we found that Ackr3 is highly upregulated in patients with various pathologies, including AD ^20,23–25^. Notably, our data indicate that Cxcl12 binds to PE at synaptic terminals, which in turn specifically recognized by astrocytic Ackr3 for phagocytic elimination. Modulation of Ackr3 expression in AD mouse models not only reduces synaptic removal by AD astrocytes, but also recues cognitive impairments, highlighting the Ackr3-Cxcl12- PE axis as a potential therapeutic target for AD-related synapse loss.

## Results

### Identification of novel regulators of astrocyte-mediated synapse engulfment

To identify novel modulators involved in synapse engulfment mediated by astrocytes, we applied a siRNA-based screening approach (Fig.1A) and measured the internalisation of purified mouse brain neuronal terminals, namely synaptosomes, as final readout. To this aim we first optimised a protocol for the derivation and labelling of synaptosomes and subsequently set the parameters for our phagocytic assay. Synaptosomes have been isolated from whole mouse brains (Fig.S1A) ^26^ and the purity of the preparation has been assessed by western blot. As shown in Fig.S1B-C, the synaptosome fraction (S) was enriched in synaptic markers such as Vesicle associated membrane protein2 (Vamp2), Synaptophysin (Syp) and Postsynaptic Density Protein 95 (PSD95). The presence of markers associated with myelin and astrocytic contamination, namely Myelin Binding Protein (MBP) and Glial Fibrillary Acidic Protein (GFAP) was instead minimised (Fig.S1 B-C). By means of transmission electron microscopy (TEM) we could identify both pre (PSc) and post (PO) synaptic terminals, thus confirming the ultrastructural integrity of synaptosomes in our preparation (Fig.S1D). Furthermore, we characterised synaptosome protein composition by mass spectrometry analysis and assessed the synaptic gene enrichment using the SynGO geneset tool (https://www.syngoportal.org). As expected, both PSs and PO synaptic structures were represented, once more confirming the reliability of our preparation (Fig.S1E). Finally, to visualise the internalisation of synaptosomes and follow their progression along the phagocytic pathway, synaptosomes have been conjugated with pHrodo-Red TM succinimidyl ester, a pH sensitive dye (pH-RODO) which becomes fluorescent (red) when exposed to an acidic environment (e.g., endo-lysosomes), as described in ^27,28^. For the phagocytic assay, primary mouse astrocytes were co-transfected with a siRNA scramble (siControl) and an eGFP-encoding plasmid used to visualize the cell area, as previously described in ^29^. We assessed the identity of the eGFP-transfected cells by immunofluorescence staining using GFAP for astrocytes and Ionized calcium-binding adaptor molecule 1 (Iba1) for microglia (Fig.S2A-B). We did not detect any eGFP^+^/GFAP^-^ or eGFP^+^/Iba1^+^ cells, thus confirming that all the eGFP expressing cells were primary astrocytes (Fig.S2A-B). The day after co-transfection, astrocytes were treated with pH-RODO labelled synaptosomes. After washing the excess of synaptosomes, we acquired the images using an automatized live-imaging system and investigated the ability of astrocytes to internalise synaptosomes by calculating the Phagocytic Index (PI) as the ratio between the red area (pH-RODO) and the green area (eGFP) after 12h and 48h (Fig.1A). As shown in Fig.1B (black line), PI increases over time as already reported in ^27^. To further confirm that the sparse microglia contamination was not interfering with our readout, we treated our cells with liposomal clodronate, a well-known compound that eliminates microglial cells without affecting astrocytes and analysed the PI of eGFP^+^ cells over time ^30^. As shown in Fig.S2C, we did not detect any statistical differences in the PI in treated and untreated cells. To validate our experimental set up, we genetically manipulated the phagocytic pathway by downregulating Megf10, a receptor that recognises and promotes the engulfment of neuronal terminals ^4,31^ and Cathepsin B (CtsB), an enzyme that promotes lysosomal degradative capacity (Fig.S2D-F) ^32^. The efficacy of the downregulation in primary astrocytes was tested by western blot and compared to a siRNA scramble (siControl) (Fig.S2D). Overall, we confirmed a significant halving of Megf10 and CtsB expression compared to control (Fig.S2D-G). We also analysed the kinetics of internalisation, and we observed that the downregulation of Megf10 and CtsB caused significant changes in synaptosome uptake in primary astrocytes (Fig.1B-C). As expected, Megf10 downregulation reduces the phagocytic abilities of astrocytes compared to the siControl at both 12h and 48h (Fig.1B-C). Instead, the downregulation of CtsB causes an accumulation of engulfed synaptosomes, specifically at 48h compared to siControl, likely due to deficits in synaptosome degradation (Fig.1B-C).

**FIGURE 1.**
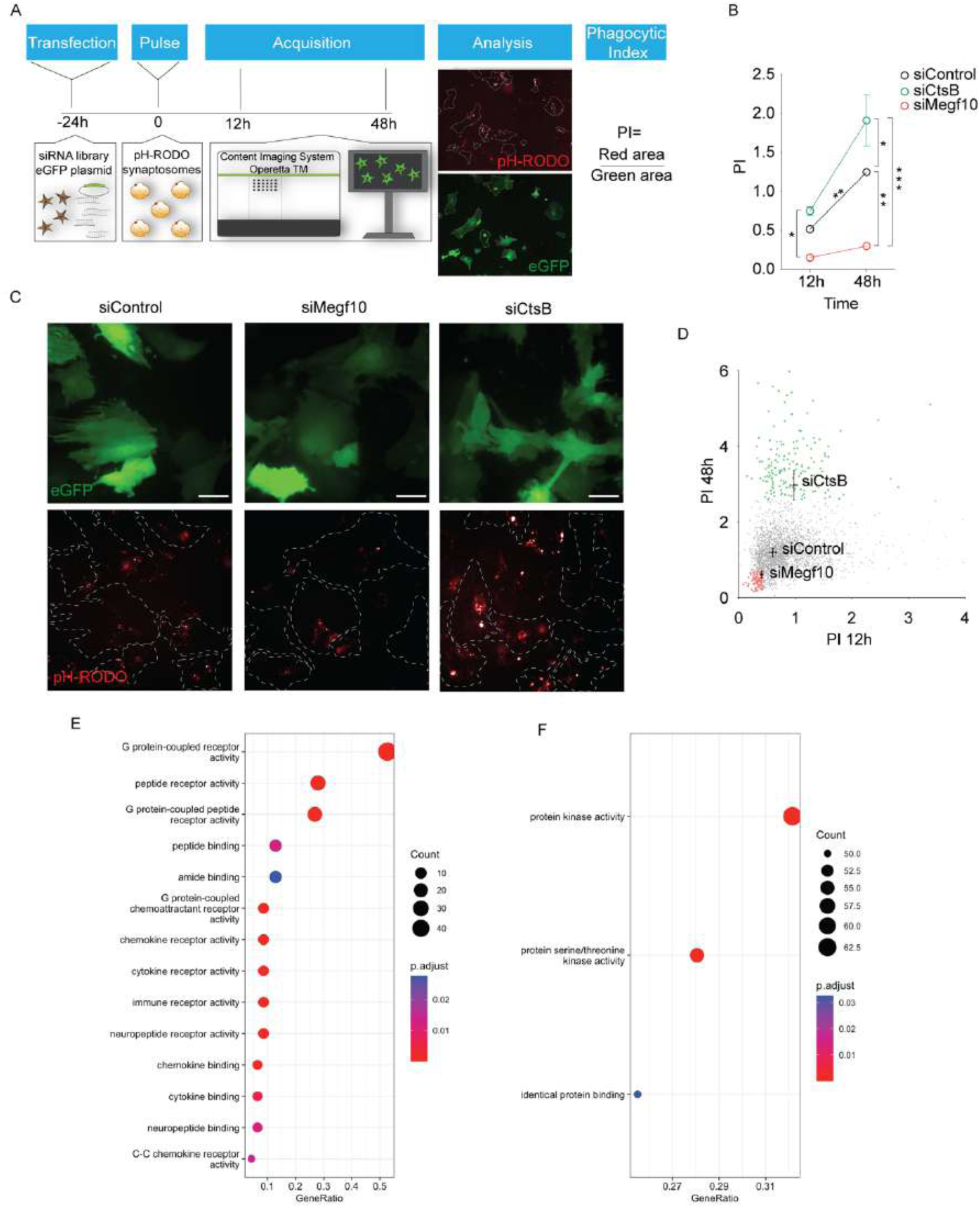
Genetic screening reveals novel modulators of synaptosome engulfment. **A.** Experimental design. Primary astrocytes were transfected with eGFP encoding plasmids in combination with siRNA oligonucleotides. The next day, cells were exposed to pHRODO-labelled synaptosomes and the phagocytic kinetics were measured through long-term fluorescence live imaging at 12 and 48 h. To quantify phagocytosis, the area (μm^2^) of red fluorescence (pHRODO) signal was normalized to the eGFP positive area (μm^2^) (Phagocytic Index, PI). **B.** PI of primary astrocytes transfected with siRNA Control (black line), siRNA against Megf10 (red line) and siRNA against CtsB (green line). (siControl at 12h vs 48h, p=0.0014; 12h: siCtsB vs. siMegf10 p= 0.0453; 48h: siControl vs. siMegf10 p=0.0037; siControl vs. siCtsB p= 0.0266; siCtsB vs. siMegf10 p <0.0001). N=40 images per condition were analysed; N=3 biological replicates. Statistical analysis was performed using Two-Way ANOVA with Tukey’s multiple comparison test. **C.** Representative images of primary astrocytes transfected with Control, Megf10 and CtsB siRNA and the eGFP- encoding plasmid at 48h. pH-RODO fluorescence has been analyzed within the eGFP positive area (dotted line) Scale bar 20 μm. **D.** Graph representing the PI at 12h vs 48h of primary astrocytes transfected with the siGENOME Mouse Druggable siRNA Library (one dot=1 hit). Downregulation of genes that abolishes the ability of astrocytes to internalize synaptosomes were highlighted as red hits; while downregulation of genes that enhances PI values compared to control were highlighted as green hits. Five images per condition were analysed. **E.** Dotplot of red hits. Genes are plotted with a qvalue <0.5. Bubble plots highlight p.adjusted values of the GO categories (color scale) and the number of hits falling in the GO categories (size scale). Gene ratio defines the percentage of the enriched hits in the given GO term. **F.** Dotplot of green hits. Genes are plotted with a qvalue <0.5. Bubble plots highlight p.adjusted values of the GO categories (color scale) and the number of hits falling in the GO categories (size scale). Gene ratio defines the percentage of the enriched hits in the given GO term.

We therefore performed a siRNA-based screening relying on the siGENOME Mouse Druggable siRNA Library composed of a total of 2905 genes. Upon the downregulation of each gene in primary astrocytes and the subsequent treatment with synaptosomes, PI values have been plotted at 12h (X axis) and 48h (Y axis) and compared with the control conditions (Fig.1D). Specifically, we have annotated the PI values at 12h and 48h of the siControl, siMegf10 and siCtsB (Fig.1D). As expected, most of the hits clustered around the PI values of astrocytes transfected with the siControl thus suggesting that these downregulated genes were not impacting on astrocyte-mediated synaptic clearance. However, the downregulation of several genes modulated synaptosome internalisation and accumulation. One group of hits (∼100), marked in red, represented those genes that when down- regulated reduced synapse internalisation (Fig.1D). Their PI values were in fact below the PI values of siMegf10 transfected cells (Fig.1D). A second group of hits (∼200), marked in green, instead represents those genes that potentially interfere with degradation (or enhance internalisation) (Fig.1D). Specifically, the PI values of those hits at 48h are above the PI of siCtsB transfected cells (Fig.1D). To investigate the molecular function (MF) of the hits involved in the synaptosome clearance, we performed a gene ontology (GO) analysis. We observed that those genes identified as red hits showed enrichment in G protein-coupled receptor activity, peptide receptor activity and chemokine receptor activity (Fig.1E, Fig.S3A). Specifically, G protein-coupled receptors and chemokine receptors are the top represented categories. Genes that are mainly represented in these categories are 8 chemokine receptors: Cxcr5, Ackr3, Cxcr2, Cxcr3, Ccr1, Ccr2, Ccr4 and Ackr2 (Fig.S3A, yellow dots). Instead, MF analysis for the green hits identified as top categories proteins with kinase activity, serine/threonine kinase activity and identical protein binding activity (Fig.1F, Fig.S3B).

Overall, our siRNA-based screening revealed both a cluster of genes encoding proteins with receptor activity that, when down-regulated, affected synaptosomes internalisation and a second group of genes encoding proteins involved in intracellular signalling pathways that, when downregulated promote intracellular synaptosome accumulation.

### Ackr3 is required for synaptosome internalisation

Our screening, coupled with the GO analysis, revealed a family of chemokine receptors whose downregulation impairs synaptosomes internalisation mediated by astrocytes. Using the transcriptomic dataset published in ^33^, we looked at the differential expression of these chemokine receptors in astrocytes, microglia and neurons (Fig.2A). Among these, we selected for validation Ackr3, based on its high expression in astrocytes as compared to the other two cell types. First, we confirmed that Ackr3 is endogenously expressed in primary mouse astrocytes, and it is efficiently downregulated using a siRNA pool against Ackr3 (Fig.S4A). As visualised by western blot, the global downregulation in mouse astrocytes reached 60% as compared to astrocytes transfected with siRNA Control (Fig.S4A-B). We then performed longitudinal live imaging of synaptosome engulfment following the same set up used for the siRNA-based screening. Internalisation was followed over time at different time points: 6h, 12h, 24h, 48h, 78h, 96h, 120h (Fig.2B-C). As expected, the internalisation of synaptosomes was abundantly suppressed in murine astrocytes downregulated for Ackr3 as compared to siControl transfected astrocytes resulting in a statistically significant decrease of the PI at 24h, 48h and 78h (Fig.2C). To further corroborate the involvement of Ackr3 in the modulation of synaptosome internalisation, we overexpressed the receptor in primary astrocytes using a Flag-tagged Ackr3 encoding plasmid (Fig.S4C-D). Here, the PI calculation over time confirmed that the ectopic expression of Flag-Ackr3 enhances astrocytes mediated internalisation of synaptosomes (Fig.S4C-D, blue closed dots) and this phenotype is partially reverted by Ackr3 downregulation (Fig.S4C-D, light blue closed dots). Likewise murine cells, human iPS-derived astrocytes (iHA) abundantly express ACKR3 and its expression can be downregulated by means of siRNA against the receptor (Fig.S4E-F). Interestingly, we observed that both human and mouse Ackr3 is distributed in patchy areas in control astrocytes (treated with PBS), possibly clustered at the plasma membrane and/or in vesicular structures, and that it only partially co-localizes with lysosomal associated membrane protein 1 (Lamp1) (Fig.2D-G; Fig.S5A-D). Instead, in astrocytes treated with TAMRA-labelled synaptosomes, Ackr3 decorates the internalized structures and massively re-localizes into Lamp1-positive vesicular structures (Fig.2E-G; Fig.S5B-D).

**FIGURE 2.**
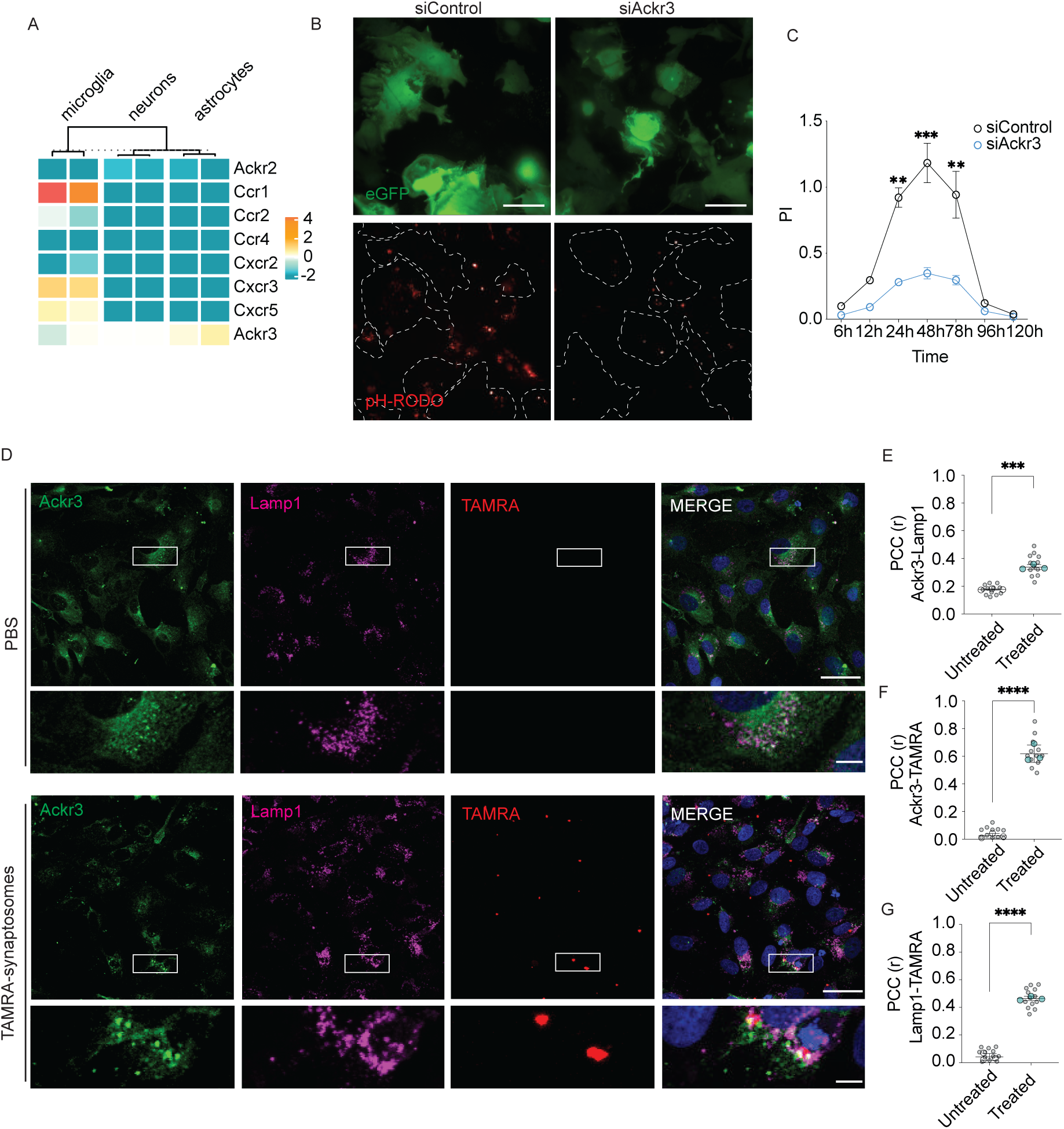
Ackr3 regulates synaptosome internalization in astrocytes. **A.** Transcriptomic data showing the differential expression of the chemokine receptors in astrocytes, microglia and neurons isolated from mice. The heatmap shows the RNA-seq analysis obtained from^33^. Columns show the duplicates for each type of cell, while rows represent the genes **B.** Representative images of primary astrocytes transfected with siControl and siAckr3 together with an eGFP-encoding plasmid followed by pH-RODO synaptosome treatment (48h). Scale bar 20 μm. **C.** PI of primary astrocytes transfected with siControl and siAckr3 at 6h, 12h, 24h, 48h, 78h, 96h and 120h upon synaptosomes treatment. Ackr3 (blue line) downregulation displayed a significant higher PI at 24h, 48h and 78h compared to control (24h: Ackr3 vs. Control p=0.0002; 48h: Ackr3 vs. Control p <0.0001; 78h: Ackr3 vs. Control p=0.0002). N=30 images per conditions were analysed; N=3 biological replicates. Statistical analysis was performed using Two-Way ANOVA with Tukey’s multiple comparison test. **D.** Representative z-stack confocal images of iHA treated with PBS or TAMRA-labelled synaptosomes (red). Cells were stained with anti-Ackr3 (green) and anti-Lamp1 (purple) antibodies. Scale bar 20 μm; insets 5 μm. **E.** Pearson’s correlation coefficient (PCC) of human Ackr3/TAMRA co-localization (untreated vs. treated p=0.0004). **F.** PCC of TAMRA/Lamp1- positive compartment co-localization (untreated vs. treated p<0.0001). **G.** PCC of Ackr3/Lamp1- positive compartment co-localization (untreated vs. treated p<0.0001). **E-G.** N=6 images for conditions were analysed; N=3 subjects. Statistical analysis in E-G was performed using unpaired t- tests.

Overall, our results suggested that Acrk3 is required for astrocyte-mediated synaptosome internalisation, establishing a molecular complex directed to the lysosome upon engulfment.

### Ackr3 recognizes chemokines on the synaptosome surface

We next examined a possible physical interaction between Ackr3 and the synaptosomes. Being Ackr3 a chemokine receptor ^20–22^, we investigated the role of its ligands, Cxcl11 (K_d_:2.5 nM) and Cxcl12 (K_d_:0.3 nM) in the internalisation of synaptosomes ^22,34–39^. First, we examined the chemokines’ content in the whole mouse brain (WB), in our synaptosome preparation (S) as well as in murine brain cells (primary astrocytes and primary neurons, A and N, respectively) (Fig.S6A). We observed that Cxcl11 is detected in the whole brain lysate and in astrocytes, but it is not present neither in the synaptosome preparation nor in neurons (Fig.S6A). Cxcl12 is instead present in the whole brain lysate as well as in neurons, astrocytes and in the synaptosome preparation (Fig.S6A). We could therefore speculate that the binding between Ackr3 and the synaptosomes is mediated by Cxcl12. To test this hypothesis, we performed a batch of experiments in which we modulated the availability of chemokine binding site in Ackr3 or the chemokine at the synaptosome surface. First, we set our basal condition by incubating purified Flag-tagged Ackr3 with synaptosomes. As expected, Vamp2, the synaptosome marker, was detected in the input (synaptosome preparation plus Flag-only or Flag- Ackr3-bound resin) both in the presence or absence of Ackr3. However, upon immunoprecipitation (IP), Vamp2 was retained only by the Flag-Ackr3-bound resin but not by the control one (Flag-only) confirming the specificity of Ackr3-synaptosomes interaction (Fig.3A-B). To test the role of Cxcl12 in mediating the binding of Ackr3 with synaptosomes, we saturated purified Flag-Ackr3-bound resin with an excess of recombinant murine SDF-1α-Cxcl12 or with Plerixafor, a well-known allosteric agonist for Ackr3 ^40^. We observed that the preincubation of the receptor with either Cxcl12 or Plerixafor interfered with Ackr3-synaptosome binding, as shown by a reduced co-immunoprecipitation of Vamp2 (Fig.3A-B). In a complementary experiment, we saturated purified synaptosomes with an anti-Cxcl12 antibody to reduce the amount of available Cxcl12. As expected, also in this case we observed a decrease in the binding between Ackr3 and the synaptosomes (Fig. S6C). Once attested the presence of Cxcl12 in our synaptosome preparation (Fig.S6A-C) and its functional involvement in the Ackr3-synaptosome interaction, we investigated whether the enrichment of Cxcl12 at the synaptosome surface could have an impact on the synaptosomes internalisation by astrocytes. Of note, intact synaptosomes are competent to bind recombinant Cxcl12 and the amount of retained recombinant chemokine increases with the amount of synaptosomes (Fig.S6D-F). We therefore incubated our synaptosomes with recombinant Cxcl12 to obtain Cxcl12- covered synaptosomes (cS). As shown in Fig.S6C, cS were effectively enriched with Cxcl12. We then investigated whether this overload of Cxcl12 could influence the rate of synaptosome internalization by looking at the PI of primary astrocytes treated with S or with cS. As expected, we observed that cS were internalised more efficiently as compared to S (Fig.3C-D), thus confirming the functional role of Cxcl12 in synapse internalization. In agreement with our previous results, the internalisation of S and cS synaptosomes was reduced in Ackr3-downregulated astrocytes, further confirming that both Ackr3 and CxCl12 are necessary to promote astrocyte-mediated internalization of synaptosomes.

**FIGURE 3.**
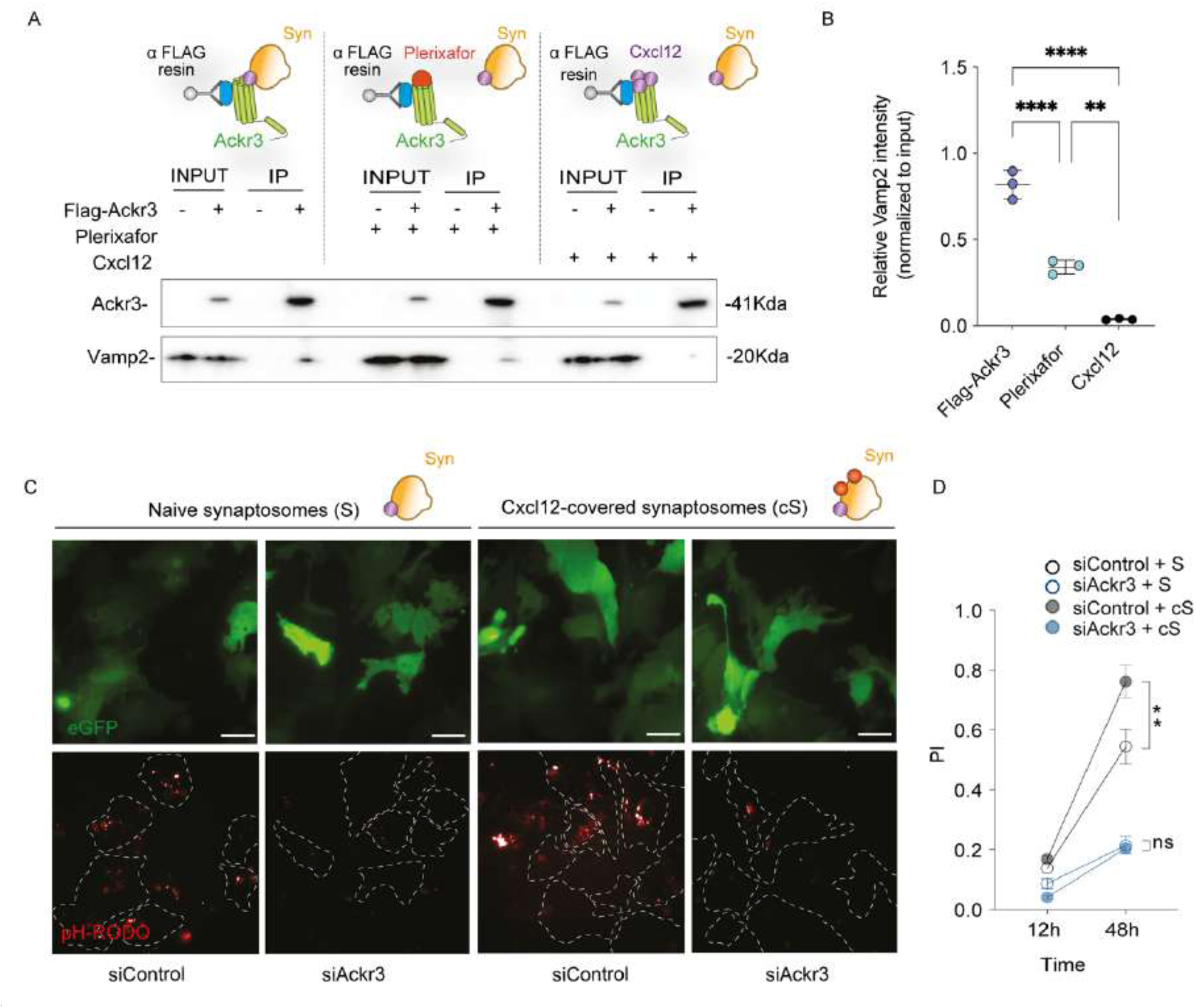
Cxcl12 modulates synaptosome recognition. **A.** Western blot analysis of pull-down assay using purified Flag-Ackr3 as bait and intact synaptosomes as prey. Untreated Flag-Ackr3 or preincubated Flag-Ackr3 with Plerixafor and Cxcl12 has been tested. Bound synaptosomes were detected using anti-Vamp2. **B.** Quantification of immunoprecipitated Vamp2 (IP) normalized to total Vamp2 (input). N=3 biological replicates. Statistical analysis was performed using One-Way ANOVA with Tukey’s multiple comparison test (Flag-Ackr3 vs. Plerixafor p<0.0001 Flag-Ackr3 vs. Cxcl12 p<0.0001; Plerixafor vs. Cxcl12 p=0.0011). **C.** Representative fluorescence images of primary astrocytes transfected with siRNA Control or siRNA against Ackr3 together with eGFP-encoding plasmid upon 48h of S or cS treatment. Scale bar 20 μm. **D.** PI of primary astrocytes transfected with siRNA Control (grey) or siRNA against Ackr3 (light blue) at 12 and 48 h upon treatment with S (empty dots) or cS (filled dots). N=12 images per condition were analysed; N=3 biological replicates. Statistical analysis was performed using Two-Way ANOVA with Tukey’s multiple comparison test (12h: siRNA Control treated with S vs. siRNA Control treated with cS p=0,0009; 12h: siRNA against Ackr3 treated with S vs. siRNA against Ackr3 treated with cS p>0.9999; 48h: siRNA Control treated with S vs. siRNA Control treated with cS p=0.0093; 48h: siRNA against Ackr3 treated with S vs. siRNA against Ackr3 treated with cS p>0.9999).

Taken together, these data reveal that Ackr3 recognizes Cxcl12 at the synaptosome surface and that this chemokine regulates Ackr3-mediated synaptosome uptake by astrocytes.

### Cxcl12 binds externalised PE at neuronal surface, a stimulus required for Ackr3-mediated synapse recognition

A subset of chemokines displays high affinity binding for PS and for other anionic phospholipids ^41^. We therefore tested the affinity of Cxcl12 for phospholipids. First, we probed PIP Strips™ membranes with recombinant Cxcl12 (Fig.4A). Among the 15 different phospholipids spotted on the array, Cxcl12 showed high affinity for PE and for phosphatidic acid but not for PS (Fig.4B). Although less investigated, PE are also lipids that are externalised in the early phase of neuronal death ^42^. We therefore set up a protocol to increase the exposure of PE in primary neurons and investigate whether this was sufficient to promote their interaction with Ackr3. Internalization and externalization of PS and PE are dynamic processes and are regulated by specific phospholipid scramblases and flippases ^43,44^. At the synapse, the stimulation of the externalisation of aminophospholipids can be induced by downregulating cell division control protein 50 a (Cdc50a), an accessory subunit of the lipid flippase P4-ATPase ^45,46^. First, we confirmed PS exposure in unpermeablized cortical mature neurons using pSIVA, an annexin-based fluorescent biosensor that specifically binds to PS (Fig. S7A-B) ^47,48^. Live neurons transfected with siControl or siCdc50a have been incubated with pSIVA and then fixed for subsequent staining of the neuronal marker Vamp2 ^49^. We reported a significant increase of Vamp2/pSIVA and PSD95/pSIVA upon Cdc50a downregulation, thus confirming PS externalisation in synaptic structures (Fig.S7A-C). Likewise, we used green fluorescent tagged Duramycin (Duramycin-Fc) to examine the exposure of PE upon Cdc50a downregulation in primary neurons. Indeed, Duramycin is a lantibiotic peptide derived from *Streptoverticillium cinnamoneus*, which binds PE at a 1:1 molar ratio with high affinity and specificity (Kd 4-6 nM) ^50,51^. Interestingly, we found that the downregulation of Cdc50a causes a significant exposure of PE at the synapse as visualised by the quantification of Vamp2/Duramycin-Fc and PSD95/Duramycin-Fc co-localization (Fig.S7F). Overall, our data confirmed that the downregulation of Cdc50a increased the exposure of both PS and PE at the level of the synapses. We therefore used this model to investigate whether a higher exposure of PE increases Cxcl12 amount at the neuronal surface, thus stimulating glial Ackr3- mediated synapse elimination.

**FIGURE 4.**
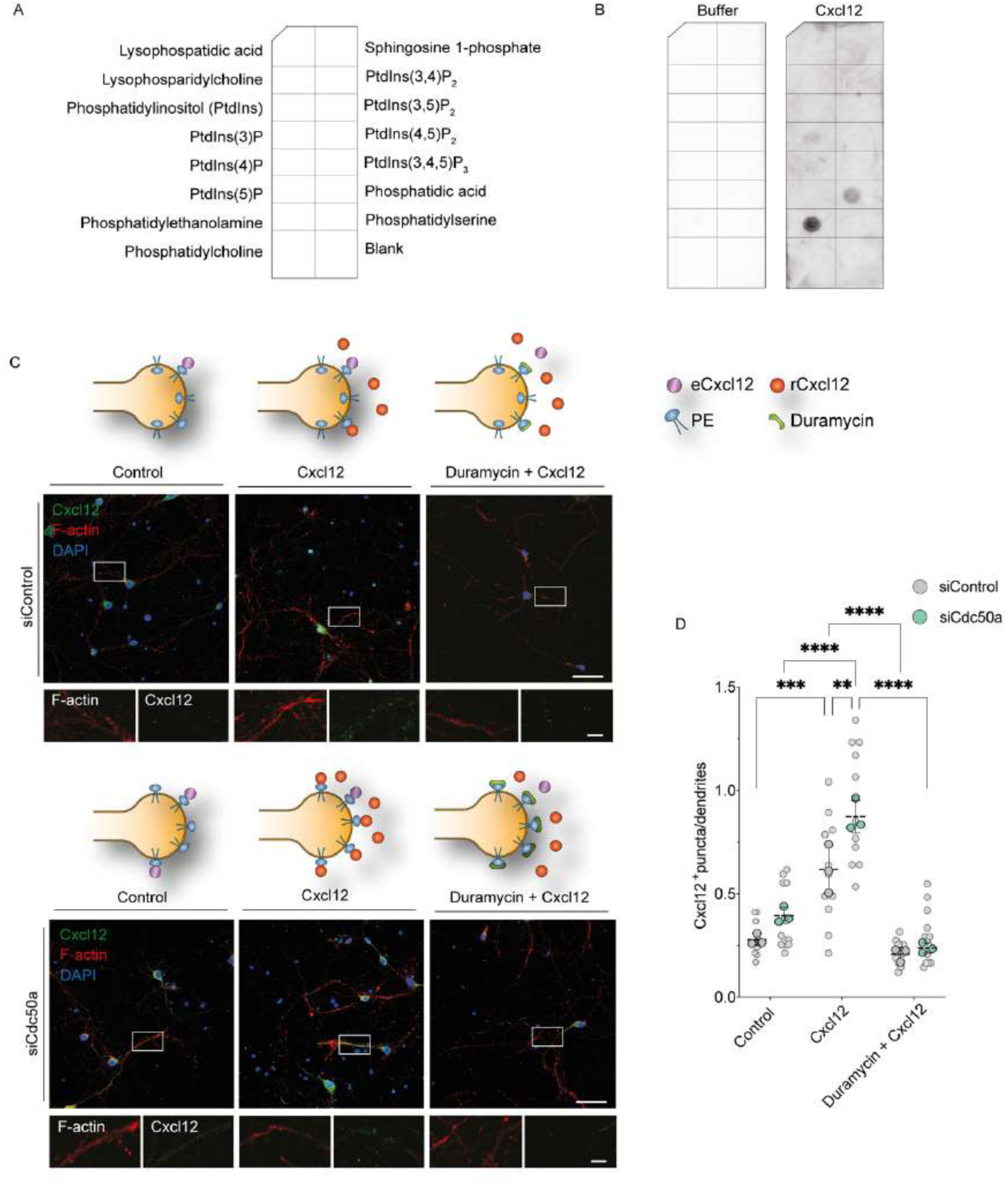
Cxcl12 binds externalized PE. **A.** Schematic picture of arrays spotted with 15 different phospholipids. **B.** Array incubated with RIPA buffer (left) or 0.1μg/ml of recombinant Cxcl12 (right). Bound chemokine was detected with the anti-Cxcl12 antibody. **C.** Representative fluorescence images of primary cortical neurons transfected with siRNA Control or siRNA against Cdc50a. Fixed, unpermeabilized cells were treated or not (control) with recombinant Cxcl12 in the presence or in the absence of duramycin preincubation. Insets show Cxcl12 (green) puncta along the F-actin positive neuronal dendrites (red). Scale bar 20 μm; inset scale bar 5 μm. **D.** Quantification of Cxcl12-positive puncta along the neuronal dendrites (Control/siControl vs. Cxcl12/siControl p=0,0003; Control/siControl vs. Cxcl12/siCdc50 p<0.0001; Control/siCdc50a vs. Cxcl12/siControl p=0.0077; Control /siCdc50a vs. Cxcl12/siCdc50a p<0.0001; Cxcl12/siControl vs. Cxcl12/siCdc50a p=0,0038; Cxcl12/siControl vs. Duramycin+Cxcl12/siControl p<0.0001; Cxcl12/siControl vs. Duramycin + Cxcl12/siCdc50a p=0.0001; Cxcl12/siCdc50a vs. Duramycin + Cxcl12/siControl p<0.0001; Cxcl12/siCdc50a vs. Duramycin+Cxcl12/siCdc50a p<0.0001). N=12 images per condition were analysed; N=3 biological replicates. Statistical analysis was performed using Two-Way ANOVA with Tukey’s multiple comparison test.

We first quantified endogenous Cxcl12 present at the neuronal surface upon the downregulation of Cdc50a and compared to control cells. Unpermeabilized cells were stained for Cxcl12 and F-actin and quantified as the number of Cxcl12^+^ puncta per μm of dendrite section. As shown in Fig.4C-D, there was a slight but not significant difference in the amount of Cxcl12 along dendrites (F-actin) between neurons transfected with siControl and those transfected with siCdc50a. However, the incubation of neurons with exogenous, recombinant Cxcl12 promoted an increase of the chemokine signal along dendrites both in siControl and, more significantly, in siCdc50a transfected neurons (Fig.4C-D). The amount of bound Cxcl12 to the dendrites reflects the level of externalised PE in siControl versus siRNA Cdc50a (Fig.S7D-E-F). This result agrees with the observation that a downregulation of Cdc50a induces the exposure of PE, which likely binds a higher amount of Cxcl12. Notably, treatment with Duramycin before incubation with Cxcl12 prevented the binding between Cxcl12 and the neurons, likely due to competitive binding, thus pointing to PE as the Cxcl12 binding partner at the synapse (Fig.4C-D).

We next sought to determine whether PE-positive synapses are more readily internalized by astrocytes and whether this process is regulated by Ackr3. To investigate this, primary neurons were genetically depleted of Cdc50a (N-siCdc50a) or treated with control siRNA (N-siControl), and co-cultured with CellTrace-labeled astrocytes in which Ackr3 was either downregulated (A-siAckr3) or left unaltered (A-siControl). We then quantified the number of Vamp2⁺/PSD95⁺ puncta (representing synapses) located within astrocytes. Consistent with our hypothesis, we found that Cdc50a downregulation in neurons, which increases PE exposure, enhances the internalization of Vamp2⁺/PSD95⁺ synaptic puncta by astrocytes (Fig. 5A–B; Fig. S8A–D). Notably, downregulation of Ackr3 in astrocytes significantly impaired their ability to engulf synapses, despite the elevated PE exposure, confirming that Ackr3 is essential for the internalization of live synapses (Fig. 5A–B; Fig. S8A–D). To further elucidate the source of Cxcl12 in neuron-astrocyte co-cultures, we selectively downregulated Cxcl12 expression in neurons (N-siCxcl12), astrocytes (A-siCxcl12), or both. The efficiency of Cxcl12 knockdown in each cell type is shown in Fig. S9A-C. We then quantified the number of Vamp2⁺/PSD95⁺ synaptic puncta internalized by astrocytes (CellTrace-labeled) under these conditions (Fig. 5C-D). In all experiments, PE externalization in neurons was induced via Cdc50a downregulation. Our data show that Cxcl12 depletion in neurons modestly reduces synaptic engulfment by astrocytes. However, Cxcl12 knockdown in astrocytes, or in both neurons and astrocytes, markedly diminished synapse internalization, indicating that astrocytes are the primary source of Cxcl12 required for effective synaptic engulfment (Fig. 5D).

**FIGURE 5.**
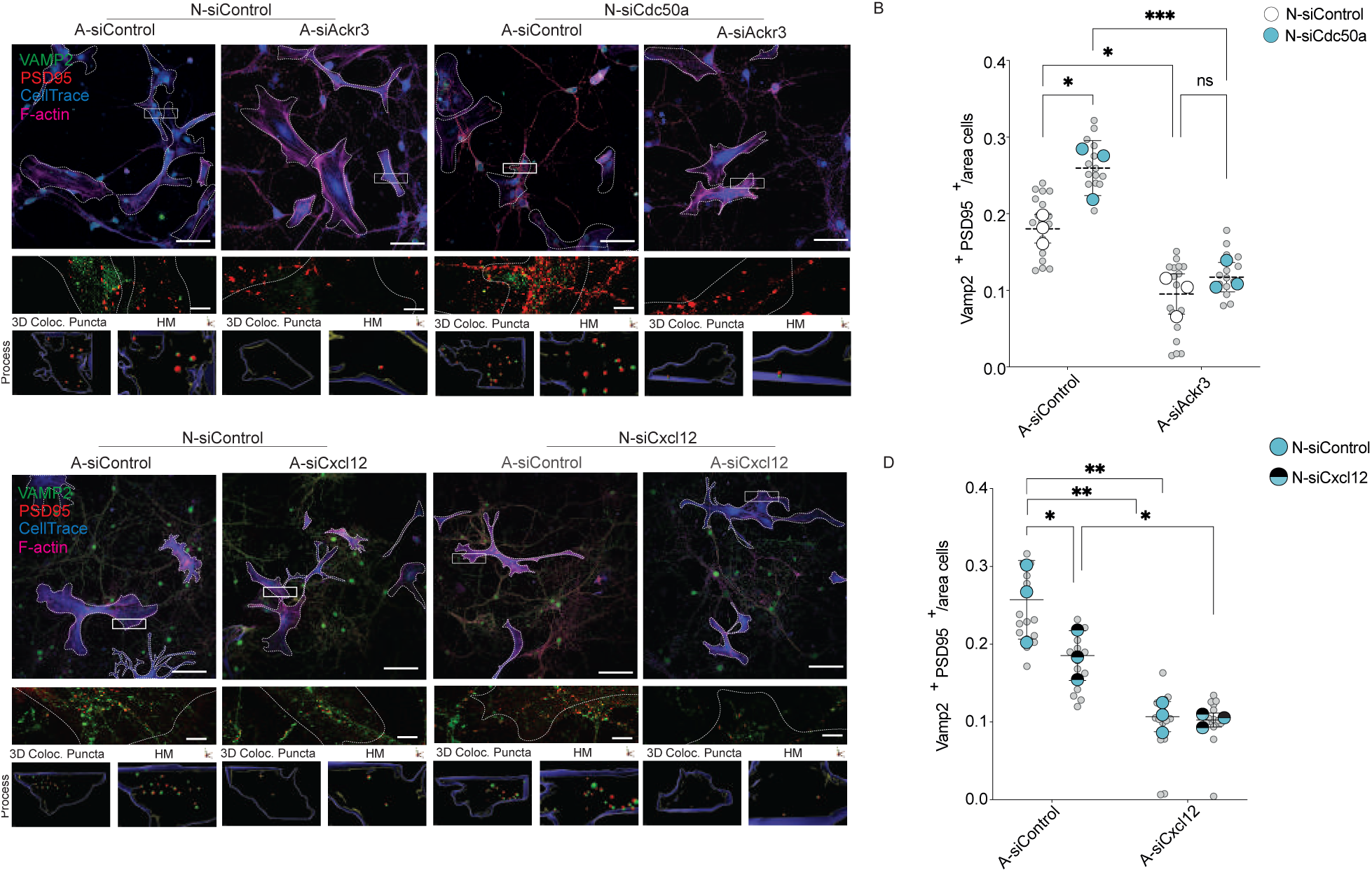
PE externalization stimulates Ackr3-mediated synaptic removal. **A.** Representative fluorescence images of primary neurons co-cultured with primary astrocytes. Neurons were transfected with siRNA Control (N-siControl) or siRNA against Cdc50a (N- siCdc50a). Synaptic terminals were stained with anti-PSD95 (red) and anti-Vamp2 (green). Dendritic trees are marked using F-actin (magenta). Astrocytes were transfected with siRNA Control (A- siControl) or siRNA against Ackr3 (A-siAckr3) and pre-stained with Cell Trace (blue). 3D rendering showing Vamp2+/PSD95+ co-localizing spots inside the astrocytes. 3D reconstructions were done using IMARIS software (Bitplane 9.9.1). Scale bar 20 μm, inset scale bar 5 μm **B.** Number of synaptic terminals (Vamp2+/PSD95+) within the astrocyte area (A-siControl /N-siControl vs. A- siControl/N-siCdc50a p=0,0237; A-siControl /N-siControl vs. A-siAckr3 /N-siControl p=0,0163; A-siControl /N-siCdc50a vs. A-siAckr3/N-siCdc50a p=0,0007; A-siAckr3 /N-siControl vs. A- siAckr3/N-siCdc50a p=0,7321) N=3 biological replicates. Statistical analysis was performed using Two-Way ANOVA with Tukey’s multiple comparison test. **C.** Representative fluorescence images of primary neurons co-cultured with primary astrocytes. Neurons were transfected with siRNA against Cdc50a together with siRNA Control or siRNA against Cxcl12. Synaptic terminals were stained with anti-PSD95 (red) and anti-Vamp2 (green). Dendritic trees are marked using F-actin (magenta). Astrocytes were transfected with siRNA Control or siRNA against Cxcl12 and pre- stained with Cell Trace (blue). 3D rendering showing Vamp2+/PSD95+ co-localizing spots inside the astrocytes. 3D reconstructions were done using IMARIS software (Bitplane 9.9.1). Scale bar 20 μm, inset scale bar 5 μm **D.** Number of synaptic terminals (Vamp2+/PSD95+) within the astrocyte area (A-siControl/N-siControl vs. A-siControl/N-siCxcl12 p=0,0161; A-siControl/N-siControl vs. A-siCxcl12/N-siControl p=0,0180; A-siControl/N-siCxcl12 vs. A-siCxl12/N-siCxcl12 p=0,0239; A- siCcontrol/N-siControl vs. A-siCxcl12/N-siCxcl12 p=0,1504) N=3 biological replicates. Statistical analysis was performed using Two-Way ANOVA with Tukey’s multiple comparison test.

Collectively, these findings reveal that Cxcl12 recognizes PE *in vitro* and this binding is stimulated by PE externalisation at the neuronal membrane. Moreover, Cxl12 released by astrocytes is key for Ackr3-mediatedsynapse recognition upon PE externalisation.

### Ackr3 is upregulated in Alzheimer’s disease astrocytes

Upregulated of ACKR3 in astrocytes is a marker of diseased brain, including AD ^23^, a neurodegenerative disorder characterized by excessive synapse elimination ^52,53^] We therefore examined the expression of the receptor in the hippocampus of AD patients and age-matched controls by immunofluorescence. Consistent with previous reports ^23^, we observed that ACKR3 is significantly upregulated in the hippocampus of AD patients compared to Healthy controls (Fig.6A- B) and the receptor exhibits a patchy distribution decorating the GFAP-positive filaments (Fig.6A), as observed in iHA and murine astrocytes (Fig.1D; Fig.S5A). Moreover, our analysis revealed an increased co-localization of ACKR3 with GFAP-positive astrocytes in the brains of AD patients compared to controls (Fig. 6C). Consistently, we also found that the CXCL12 chemokine signal is increased and enhances its co-localization with ACKR3 in AD patients compared to age-matched controls (Fig.6D-F). The increased co-localization of CXCL12 with ACKR3 indicates that this receptor-ligand interaction is heightened during pathology, further supporting the role of this pathway in synapse elimination in the context of AD. We also investigated the expression and the distribution of Ackr3 in 5xFAD mice, a well-characterized model of AD pathology available in our laboratory ^54,55^. Consistent with our observations in the brains of AD patients, we found an accumulation of Ackr3 in the hippocampus of 5xFAD mice (Fig.7A-B). Specifically, Ackr3 accumulation is predominantly observed in astrocytes, as evidenced by its co-localization with the astrocytic marker S100b in the CA1 region of the hippocampus in 5xFAD mice, compared to wild-type (WT) controls (Fig. 7C). To further evaluate our results, we extended our analysis in APPNLGF knock-in mice ^56^ and APPNLGF crossed with PS2FAD mice ^57^, two murine models that exhibits a slower progression of AD pathology compared to the 5xFAD model. Coherently with our previous observation, we found an increased expression of Ackr3 in GFAP positive hippocampal astrocytes of AD mice model compared to the WT (Fig.S10A-D).

**FIGURE 6.**
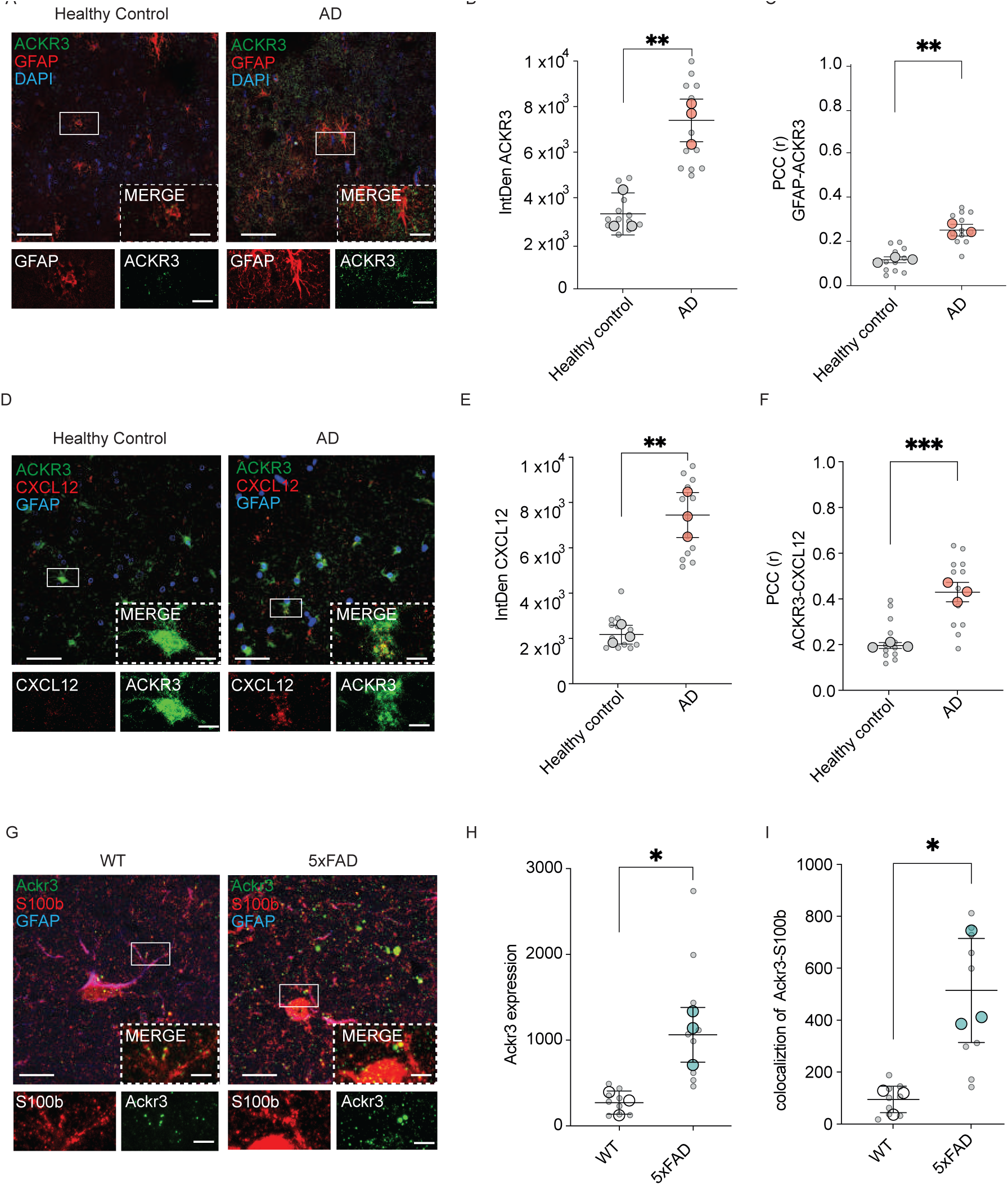
Astrocytic Ackr3 expression is AD patients. **A.** Representative double-labeling images for ACKR3 (green) and GFAP (red) in AD human hippocampus and age-matched controls; scale bar 50 μm, inset 20 μm. **B.** Quantitative analysis of ACKR3 IntDen; N=3 age-matched control samples and AD hippocampus samples. Statistical analysis was performed using unpaired t test (Healthy control vs. AD p<0.0001) **C.** PCC of ACKR3 co-localizing with GFAP-positive astrocytes; N=3 age-matched control samples and AD hippocampus samples. Statistical analysis was performed using unpaired t test (Healthy control vs. AD p<0.0001). **D.** Representative double-labelling images for ACKR3 (green) and Cxcl12 (red) in AD human hippocampus and age-matched control; scale bar 50μm, inset 20μm. **E.** Quantitative analysis of CXCL12 IntDen; N=3 age-matched control samples and AD hippocampus samples. Statistical analysis was performed using unpaired t test (Healthy control vs. AD p=0.0010) **F.** PCC of ACKR3 co-localizing with CXCL12-positive puncta; N=3 age-matched control samples and AD hippocampus samples. Statistical analysis was performed using unpaired t test (Healthy control vs. AD p= 0.0008) **G.** Representative confocal images for Ackr3 (green), GFAP (blue) and S100β (red) in CA1 hippocampus of 4-month-old 5xFAD and WT mice. Scale bar 50 μm, inset 20 μm. **H.** Quantitative analysis of Ackr3 expression in the hippocampus of 5XFAD and WT mice; N=3 samples for both WT and 5XFAD 4-month-old mice. Statistical analysis was performed using unpaired t test (WT vs. 5XFAD p=0,0170). **I.** Quantitative analysis of Ackr3 co-localizing with S100β−positive astrocytes in the hippocampus of 5XFAD and WT mice; N=3 samples for both WT and 5XFAD 4-month-old mice. (WT vs. 5XFAD p= 0,0147). Statistical analysis was performed using unpaired t test.

**FIGURE 7.**
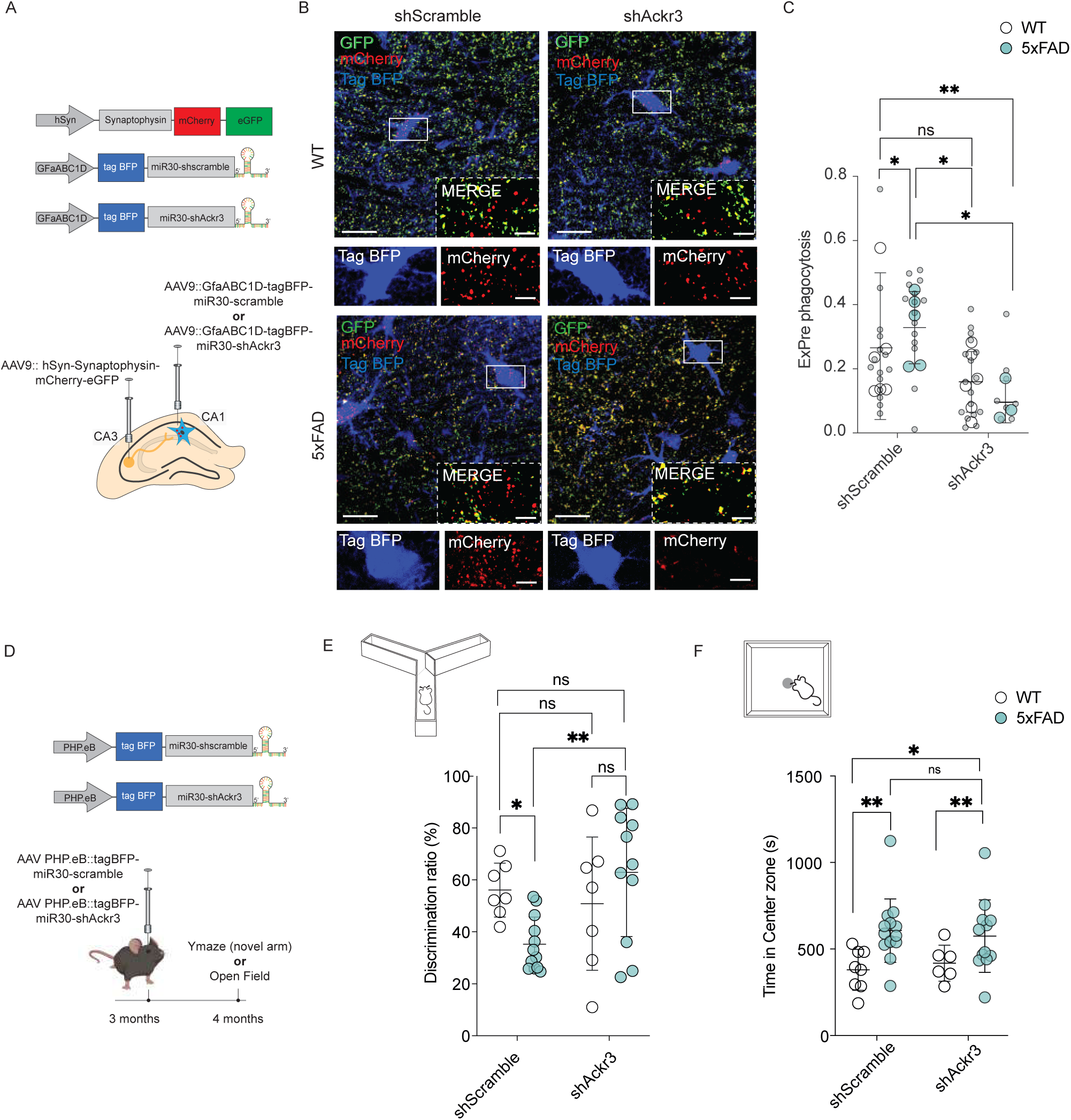
Astrocytic Ackr3 expression is stimulated in 5xFAD mice. **A.** Schematic illustration of: mCherry-eGFP synapse reporter, astrocytic shRNA-scramble and astrocytic shRNA-Ackr3 reporters. Experimental scheme of AAV reporter injection in the CA1 and CA3 hippocampal region of 1-year-old 5xFAD mice. **B**. Representative confocal z-stack images of GFP- (green) and mCherry- (red)-positive excitatory synapses in the hippocampus of 1-year-old 5XFAD and WT mice injected with AVV9-GfaABC1D-tagBFP-shScramble and AVV9- GfaABC1D-tagBFP-ShAckr3. Scale bar 50 μm, inset 20 μm. **C.** Quantitative analysis showing the volume of ExPre engulfed by astrocytes in the hippocampus of 1-year-old 5XFAD and WT mice injected with AVV9-GfaABC1D-tagBFP-shScramble and AVV9-GfaABC1D-tagBFP-ShAckr3. Scale bar 20 μm; N=5 samples for WT and N=3 samples for 5XFAD mice. Statistical analysis was performed using Two-Way ANOVA. **D**. Schematic illustration of shRNA-scramble and astrocytic shRNA-Ackr3 reporters. Experimental scheme of retro-orbital AAV reporter injection in the 3- month-old 5xFAD and WT mice. **E.** Y-maze behavioural analysis showing the 4-month-old 5XFAD and WT mice injected with AVV9-PHP.eB-tagBFP-shScramble and PHP.eB-tagBFP-ShAckr3. N=5 samples for WT and N=7 samples for 5XFAD mice. Statistical analysis was performed using Two- Way ANOVA. **F.** Open field behavioural analysis showing the 4-month-old 5XFAD and WT mice injected with PHP.eB-tagBFP-shScramble and PHP.eB-tagBFP-ShAckr3; N=5 samples for WT and N=7 samples for 5XFAD mice. Statistical analysis was performed using Two-Way ANOVA.

Overall, our data indicate that this receptor is consistently upregulated under pathological conditions in the hippocampus of AD patients and across various AD mouse models, supporting its involvement as part of a bona fide disease-associated pathway.

### Ackr3 depletion rescues pathological phenotypes in AD mice

Given the correlation between synaptic terminal loss and receptor overexpression in glial cells during Alzheimer’s disease (AD), we sought to functionally validate the role of Ackr3 in AD mouse models. To this end, we employed an *in vivo* assay to quantify astrocyte-mediated synaptic engulfment using the mCherry-eGFP fluorescent phagocytosis reporter system that we previously developed ^11,58–60^ (Fig. 7A). Briefly, we generated adeno-associated virus 9 (AAV9) vectors encoding a human synapsin promoter (hSYN) to drive the expression of Synaptophysin-mCherry-eGFP (Fig.7A). This vector was injected into the CA3 hippocampus of 5xFAD mice, and the corresponding neuronal terminals were visualized in the CA1 hippocampal region (Fig. 7A). Functional neuronal terminals are detected as yellow puncta (eGFP and mCherry), whereas phagocytosed synapses are recognized as red puncta due to the rapid denaturation of eGFP in acidic glial lysosomes (Fig. 7A-B). To genetically modulate Ackr3 expression at the CA1 region *in vivo*, we generated AAV9 vector encoding shRNA against Ackr3 together with a Blue Fluorescent Protein (BFP) tag under the GfaABC1D promoter to assure expression in astrocytes (Fig.7A). As control, we produced AVV9 vector encoding scramble shRNA with tagBFP under the GfaABC1D promoter (Fig.7A). The efficacy of the Ackr3 downregulation in astrocytes was validated *in vivo* by quantifying Ackr3 expression in tagBFP positive cells (Fig. S11A-B). Of note, we observed that the absence of Ackr3 does not affect GFAP expression in astrocytes (Fig. S11C). We then monitored the ability of CA1 hippocampal astrocytes to engulf Synaptophysin-mCherry-eGFP synapses *in vivo* in 5xFAD and WT mice following the injection of AAV9-shAckr3 versus AAV9-shScramble (Fig. 7B-C). Consistent with previous findings, ^12,17,61^ we observed excessive glia-mediated synaptic removal in 5xFAD mice compared to controls (Fig. 7B-C). In accordance with our in vitro data, astrocytic downregulation of Ackr3 significantly reduced the number of engulfed synapses in 5xFAD mice, indicating a key role for this receptor in aberrant synaptic elimination (Fig. 7B-C). Notably, this reduction was not observed when comparing AAV9-shAckr3 to AAV9-shScramble in wild-type mice, suggesting that Ackr3-dependent synaptic removal requires pathological, disease-specific synaptic tags (Fig. 7B-C). The 5xFAD mice exhibit behavioral impairments, including deficits in spatial working memory and increased anxiety-like behaviors, as early as four months of age ^62–64^. To determine whether early Ackr3 depletion could impact disease-associated behaviour symptoms, we retro-orbitally injected mice at 3 months of age with either AAV PHP.eB::GfaABC1D-tagBFP-miR30-scramble or AAV PHP.eB::GfaABC1D-tagBFP-miR30-shAckr3 and conducted behavioural assessments one month post-injection (Fig. 7D). As expected, 5xFAD mice displayed impaired spatial working memory relative to WT controls. Remarkably, astrocytic Ackr3 depletion significantly improved performance, restoring memory function to wild-type levels (Fig. 7E). To assess anxiety-like behaviour, we measured the time spent in the center of the open field (Fig. 7F). 5xFAD mice spent significantly more time in the center compared to WT controls, consistent with altered anxiety-like behaviour (Fig. 7F). However, Ackr3 depletion did not rescue this phenotype, suggesting that anxiety-related traits may be independent of astrocytic Ackr3 activity at this stage.

In the same mice, we investigated whether the recovery of the spatial working memory ability was due to a restoration of synapse loss by analysing the total volume of VGLUT1 and PSD95 in the hippocampus of wild-type mice and comparing them with 5xFAD mutants genetically modified for Ackr3 (Fig.S12A-B-C). Already at 4 months, a trend toward decreased synaptic volume was observed in the AD mice compared to controls, along with a significant recovery in the case of Ackr3 ablation (Fig.S12A-B-C). To indicate the specificity of the receptor’s role in synapse removal in the context of AD pathology, we reported that the increased plaque load in the hippocampus of 5xFAD mice is not reduced following astrocytic downregulation of the receptor (Fig.S12D-E). To further investigate the role of this receptor in synapse removal in AD, we overexpressed Ackr3 specifically in astrocytes of both wild-type and 5xFAD mice. Interestingly, Ackr3 overexpression had no detectable effect in wild-type animals and did not exacerbate the pathological phenotype in 5xFAD mice, possibly due to the absence of additional tagged synapses available for engulfment (Fig. S13A-B-C-D).

Overall, these findings indicate that Ackr3 acts as a specific receptor mediating astrocyte-driven synapse elimination in AD. Limiting aberrant Ackr3-dependent synaptic engulfment in 5xFAD mice partially mitigates cognitive deficits and rescues memory-related behavioural phenotypes.

## Discussion

Astrocyte-mediated synaptic removal involves a complex and not yet fully understood network of molecular signals, receptors, and tagging mechanisms. To better characterize this phenomenon, we adopted a genetic screening approach aimed at identifying novel, cell-specific regulators of astrocyte-driven synapse engulfment. This strategy revealed key molecular interactions that modulate the process. Gene Ontology (GO) analysis identified G protein-coupled receptors (GPCRs) and C-X-C chemokine receptors as top categories associated with synaptosome internalization. We also uncovered enrichment in intracellular signaling components, such as kinases, that may influence synapse accumulation by either enhancing the rate of internalization or inhibiting degradation.

Among the genes that negatively regulate synaptosome uptake, several chemokine receptors stood out. Specifically, our screen identified eight receptors: Cxcr5, Ackr3, Cxcr2, Cxcr3, Ccr1, Ccr2, Ccr4, and Ackr2 as potential modulators. We then examined their expression in acutely isolated mouse astrocytes using published transcriptomic datasets ^33^. Ackr3 (also known as Cxcr7) emerged as the most highly expressed receptor in astrocytes, but notably not in microglia. In line with this, previous reports have demonstrated Ackr3 expression and signaling in cultured astrocytes, where it has been linked to astrocyte reactivity and disease states in both mouse and human brains ^23,36,65–67^. Ackr3 is proposed to function either as a scavenger of extracellular Cxcl12 (also known as SDF-1) or as a modulator of Cxcr4, the canonical Cxcl12 receptor ^49,68,69^. Various studies in glial, endothelial, and tumor cells have described diverse roles for Ackr3 ^70–75^. For instance, Ma and colleagues showed that Ackr3 activation in macrophages enhanced phagocytic uptake of FITC-labeled *E. coli*, an effect abolished by Ackr3 silencing ^76^.

Building on this, we demonstrated that Ackr3 knockdown in primary astrocytes significantly impaired synaptosome internalization, providing the first direct evidence of Ackr3’s role in this process. Further analysis revealed that Ackr3 interacts with Cxcl12, which we found to be associated with externalized phosphatidylethanolamine (PE) on synaptosomes and live synapses. These findings identify PE-bound Cxcl12 as a novel marker of synaptic dysfunction. Supporting this concept, a recent study showed that certain chemokines can bind exposed phosphatidylserine (PS) to promote macrophage-mediated phagocytosis ^41^. While both PS and PE exposure are linked to neuronal degeneration ^42^, only PS has previously been recognized as an "eat-me" signal for astrocytes at the synapse ^4,16^.

Future investigations should explore how PE and PS tagging is spatially and temporally regulated during neurodegeneration. Our results indicate that the amount of PE-bound Cxcl12 critically influences Ackr3-mediated synapse recognition and engulfment by astrocytes. Notably, a significant increase in astrocyte-mediated synapse removal was observed only when exogenous Cxcl12 was added, suggesting that Cxcl12 availability is a limiting factor. We further found that astrocytes not only respond to Cxcl12 but also produce it, at least in vitro, potentially establishing autocrine or paracrine feedback loops that modulate their phagocytic activity.

Together, these data place the astrocytic PE-Cxcl12-Ackr3 axis at the center of a novel mechanism underlying synapse recognition following PE externalization. However, further studies are needed to determine what triggers astrocytic Cxcl12 release and how this pathway integrates with other synaptic clearance mechanisms.

Importantly, the expression of both Cxcl12 and Ackr3 is altered in neurodegenerative diseases such as Alzheimer’s disease (AD) ^23,77^, where astrocyte-mediated synapse loss is gaining attention as a key pathological feature. Both astrocytes and microglia contribute to synaptic loss in AD, in part through MFG-E8-dependent opsonophagocytosis ^17^. Additional mechanisms, including complement dysregulation ^12,53,60^ and chronic inflammation ^19^, further promote glial synaptic pruning. Despite this, astrocyte-specific mechanisms remain poorly understood.

We observed upregulation of Ackr3 and Cxcl12 in the hippocampus of AD patients and multiple AD mouse models, correlating with increased synaptic loss. In 5xFAD mice, astrocyte-specific Ackr3 deletion reduced synaptic engulfment in the hippocampus, an effect not observed in wild-type controls. These findings suggest that Ackr3 selectively promotes synaptic elimination under pathological conditions.

Moreover, Ackr3 knockdown in young 5xFAD mice slowed disease progression. Working memory deficits, commonly linked to synaptic loss in AD, were fully rescued, with treated mice performing comparably to wild-type controls ^62,63,78^. This cognitive recovery correlated with increased synaptic volume but not with changes in amyloid plaque burden, suggesting that Ackr3 modulation specifically affects synaptic integrity rather than plaque pathology.

These findings provide strong evidence that aberrant Ackr3-mediated synaptic pruning contributes to cognitive decline in AD. Disrupting this pathway preserves neural circuitry, offering a promising therapeutic strategy. However, questions remain regarding the upstream signals that drive Ackr3 overexpression and its precise targets during disease progression.

AD is marked by multiple pathological features, amyloid-beta plaques, tau tangles, neuroinflammation, and synaptic loss, with synapse loss being one of the earliest and most predictive indicators of cognitive decline ^52,53^. Despite its central role, no approved therapies currently target synapse degeneration directly. This underscores the need for novel interventions to protect synaptic function in AD.

Glial cells are essential for synaptic homeostasis ^1^, contributing to synaptic pruning during development and plasticity ^3^. In AD, however, they become dysfunctional and excessively remove synapses ^12^, ^#160^,^17,61^. While inhibition of pan-glial pruning through the complement cascade has shown promise ^79,80^, it remains unclear whether all forms of glial pruning are detrimental. Astrocytes, though capable of synaptic engulfment, are considered non-professional phagocytes with limited degradative capacity ^81^. Excessive astrocytic phagocytosis may overload their endo-lysosomal system, leading to dysfunction and inflammation ^82,83^. Recent work has described AD-associated astrocytes with impaired lysosomal function ^54^.

Based on these insights, we hypothesize that selective inhibition of excessive astrocyte-mediated synapse elimination could preserve synaptic structure and cognitive function, while reducing astrocytic stress and reactivity.

In conclusion, our study identifies a novel astrocytic mechanism of synapse removal driven by the PE-Cxcl12-Ackr3 axis. Ackr3, a druggable receptor, emerges as a key modulator of synaptic pruning with direct implications for cognitive decline in AD. Targeting Ackr3 and its downstream signaling may offer a promising route to preserve synaptic integrity and slow disease progression. Continued investigation is needed to fully elucidate how Ackr3 is regulated in pathological contexts and how its modulation impacts the broader neurodegenerative cascade.

## MATERIALS AND METHODS

### Animals

C57Bl/6J wild-type mice were provided by the Jackson Laboratory and used for the *in vitro* studies. Housing and handling of C57Bl/6J mice were done in compliance with national guidelines. All animal procedures were approved by the Ethical Committee of the University of Padova and the Italian Ministry of Health (license 46/2012 and 105/2019). For the *in vivo* studies, experiments were approved by the Institutional Animal Care and Use Committee (IACUC) of Korea Advanced Institute of Science and Technology (KAIST). Wild-type C57BL/6J mice were purchased from DBL, and B6SJL-Tg(APPSwFlLon,PSEN1*M146L*L286V)6799Vas/Mmjax (5xFAD) mice were gifted by Prof. Chan Hyuk Kim (Seoul National University). Mice were maintained on a 12-hour light cycle and had *ad libitum* access to chow and water. Both sexes were included in all experiments. For immunofluorescence studies, all procedures on wild-type, APPNLGF knock-in ^56^ and APPNLGF crossed with PS2FAD ^57^ mice were approved by the animal ethics committee of the University of Leuven (reference number 173/2022). Mice were housed under standard conditions, on a diurnal 14- hour light/dark cycle with ad libitum access to food and water.

### Plasmids and AAV virus preparation

The pEGFP-N1 plasmid obtained from Clontech (Mountain View, CA, USA) and CXCR7-Tango plasmid (Flag-tagged ACKR3; Plasmid #66265, Addgene) were used for mammalian expression of the eGFP and Flag-ACKR3 protein, respectively. Plasmids were amplified as described in ^29^.

For *in vivo* experiments, pAAV-GfaABC1D-tagBFP-miR30 plasmid was generated by combining the tagBFP sequence with the miR30 sequence (Horizon, pGIPZ plasmid) using overlapping PCR and subcloned into pAAV-GfaABC1D-eGFP plasmid (Addgene #176861). The pAAV-GfaABC1D- tagBFP-miR30-scramble and pAAV-GfaABC1D-tagBFP-miR30-shAckr3 constructs were subcloned into the pAAV-GfaABC1D-tagBFP-miR30 vector with the following target sequences.

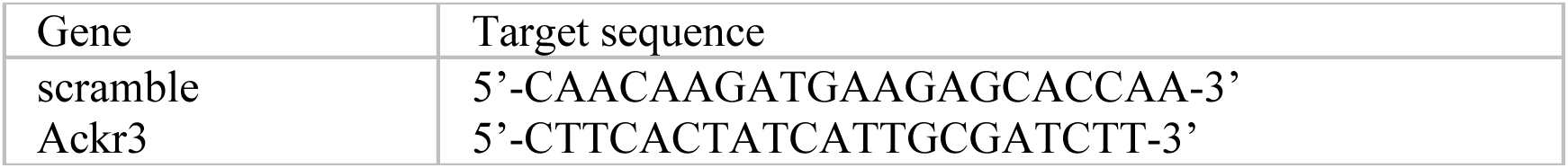

The AAV virus was harvested as previously described ^84,85^. Briefly, HEK293 cell lines were transfected with ITR-containing transfer plasmid (pAAV-GfaABC1D-tagBFP-miR30-scramble, pAAV-GfaABC1D-tagBFP-miR30-shAckr3, and pAAV-hSyn-Synaptophysin-mCherry-eGFP), a helper plasmid (Addgene #112867), and a pAAV9 capsid plasmid (Addgene #198016) with polyethylenimine (PEI, Sigma). For AAV virus purification from the cell media, supernatant was collected after 72h and 120h. Cells were harvested and lysed by freezing/thawing after 120h. Both the cell lysate and supernatant were purified with PEG8000 (Sigma) and ammonium sulfate (Sigma). The AAV virus was concentrated using Amicon Ultra (Merck), and the viral titer was quantified with quantitative PCR (qPCR) as previously described ^85^.

### Mouse primary astrocyte cultures

Mouse primary striatal astrocytes were obtained from postnatal pups between day 1 and 3 (P1-3) as described in ^29,86^. Briefly, brains were dissected from the skull and placed in cold Dulbecco’s Phosphate Buffered Saline (DPBS, Biowest). Olfactory bulbs and cortices were removed and striata were selected. After the dissection, supplemented medium composed of Basal Medium Eagle (BME, Biowest), 10% fetal bovine serum (FBS, Corning), 100 U/ml penicillin and 100 μg/ml streptomycin was added to the tissues and the striata were then sifted through a 70-μm cell strainer (Sarstedt) using a syringe plunger. The cell suspension was centrifuged (300×g, 15 min) and the pellet was washed twice with supplemented medium. Cells were seeded in cell culture flasks at the density of 5×10^6^ cells/10 ml medium and maintained in supplemented medium at 37°C and 5% CO_2_ atmosphere. The culture medium was changed after 7 days and subsequently after every 3–4 days. When cell confluency reached about 80%, contaminant microglia were detached by shaking the flask (800) for 2h at room temperature (RT). After shaking, the medium containing microglia was removed and replaced with fresh supplemented medium.

### Generation of human induced astrocytes (iHA)

Human iPS-derived astrocytes were obtained as described in ^87^. Briefly, human iPSCs were differentiated to neural precursor cells (NPCs) in laminin 521 (LN521) and laminin 211 (LN211) (both from Biolamina). Differentiation of NPCs to progenitor astrocytes was triggered in 50% laminin 111 (LN111) and 50% LN211 (Biolamina) using the astrocyte differentiation medium (STEMdiff astrocyte differentiation #100-0013, StemCell). To maintain the appropriate cell density (70% of confluence) cells were splitted once a week for 21 days. Finally, astrocytes progenitor cells were maturated in Astrocyte Maturation Medium (STEMdiff astrocyte maturation #100-0016, StemCell) for 60 to 75 days.

### Primary cortical neurons

Primary cortical neurons were obtained from postnatal pups between day 0 and 1 (P0-P1) as in ^88^. Briefly, cortices were removed, snipped into smaller pieces, transferred in papain solution (Worthington Biochemical Corporation) and further triturated. The suspension was incubated for 40 min at 37°C and spun down in a swinging-bucket rotor for 5 min at 200xg. The supernatant was discarded and 3 ml of solution containing DNase and Trypsin inhibitors were added to the pellet: after pipetting 3 times, the tubes were incubated for 10 min at RT to allow the bigger pieces of tissue to precipitate. Then, the supernatant was gently pipetted drop-by-drop upon 5 mL of bovine serum albumin (BSA) and Trypsin inhibitor solution and spun down for 10 min at 100xg. After discarding the supernatant, the pellet was resuspended in 5 ml of Neurobasal (Life Technologies) supplemented with 5% FBS, 2% B27 supplement (Invitrogen), 0.5 mM Glutamine (Life Technologies), penicillin (100Units/ml) and streptomycin (100μg/ml) (Life Technologies). 2x10^5^ cells were plated onto 12mm glass coverslips in 24-well plates and grown in Neurobasal medium supplemented as above and cultured in a 5% CO_2_ atmosphere at 37°C. After 7 days, 50% of the Neurobasal medium was removed and replaced with fresh one. Experiments were performed at DIV10.

### Neuron-glia co-cultures

To obtain neuron-glia co-cultures, primary neurons and astrocytes were prepared as described above and plated in a 5:1 ratio. Astrocytes were labelled using a pre-treatment with CellTrace Violet Cell Proliferation Kit (Thermo Fisher Scientific) at a final concentration of 1μg/ml for 20 min at 37°C and then added to the neuronal cultures. Co-cultures were incubated at 37°C and 5% CO_2_ and the synapse- engulfment experiments were carried out after 48h.

### Cell transfection

For the screening, primary astrocytes were seeded in μClear 384-well (PerkinElmer) plates at a seeding density of 4x10^3^ cells/well using Janus Liquid Handler (PerkinElmer). The day after, astrocytes were transfected using the G-014675-E2 Mouse Druggable Subset Dharmacon siGENOME® SMARTpool® siRNA Library Lot.1400 (Dharmacon). siRNA SMARTpool was mixed with pEGFP-N1 plasmid obtained from Clontech (Mountain View, CA, USA) at a final concentration of 50nM; this mixture was then combined with Lipofectamine 2000 (Thermo Scientific) following a 1:3 RNA-DNA/Lipofectamine ratio. The mixture was diluted in OPTIMEM (Gibco) and incubated for 20 min at RT to allow the formation of liposomal complexes. After 90 min, the transfection medium was replaced with complete BME.

For the live-imaging experiments, immunocytochemistry and western blot procedures, astrocytes and neurons were transfected following the protocol described above. Instead, iHA were seeded in μClear 96-well (Greiner) plates coated with LN111/LN211 (Biolamina). The day after, cells were transfected with siRNA Ackr3 (Mouse) SMARTPool (Dharmacon) at a final concentration of 100nM in combination with 0.5 μg of pmaxGFP^TM^ (LONZA) and Lipofectamine Stem Transfection Reagent (Thermo Fisher Scientific) following 1:1 siRNA-DNA/Lipofectamine ratio.

### Synaptosome purification and dye-conjugation

Synaptosomes were prepared as described in ^89^. Briefly, four Sucrose Tris-Percoll solutions (23-15- 10-3% of Percoll, respectively) were stratified into a centrifuge tube. Whole brain tissues were homogenised in a solution containing 10mM Tris, 0.32M sucrose (pH 7.4) and protease inhibitors using a glass-Teflon tissue grinder (0.25mm clearance). After centrifugation (1,000xg for 5 min at 4°C), the supernatant was added to the discontinuous density gradient solution. Upon centrifugation (33,500xg for 6 min at 4°C), the synaptosome fraction was located between the 23% and the 15% density gradient solutions. Synaptosomes were then collected and re-suspended in physiological solution (140mM NaCl; 3mM KCl; 1.2mM MgSO_4_; 1.2mM NaH_2_PO_4_; 10mM HEPES; 10mM glucose; dissolved in distilled water, pH 7.4). A final centrifugation step at 20,000xg for 10 min at 4°C was performed and the pellet containing the synaptosome fraction was quantified and either conjugated or stored at -80°C in PBS containing 5% DMSO.

Conjugation was performed as described in ^4,27^. Synaptosomes were incubated with pH-RODO™ Red, succinimidyl ester (pH-RODO) (Thermo Scientific) or 5-TAMRA (5- Carboxytetramethylrhodamine) (Thermo Scientific) in 0.1M Na_2_CO_3_ (pH 9.0) at RT with gentle agitation. After 2h, unbound pH-RODO was washed-out with DPBS by performing six rounds of centrifugations at 21,092xg for 2 min. pH-RODO-conjugated synaptosomes were re-suspended with DPBS containing 5% DMSO for subsequent freezing.

For chemokine binding assay, 10 μg of synaptosomes were incubated with recombinant Cxcl12 (DBA italia) at a final concentration of 250nM for 1h at 37°C with gentle agitation. After 2h of incubation, unbound Cxcl12 was washed-out with DPBS by performing four rounds of centrifugations at 21,092xg for 2 min.

### Mass spectrometry (LC–MS/MS)

Synaptosomes (20 μg) were sent for proteomic analysis at EMBL Proteomics Core Facility in Heidelberg, Germany, for identification and quantification of proteins in complex mixtures. Samples were subjected to in-solution tryptic digestion using a modified version of the Single-Pot Solid-Phase-enhanced Sample Preparation (SP3) ^90^. Data were analysed with MaxQuant and the iBAQ algorithm which represents the abundances of a protein within a sample. Methods are fully established and detailed in ^91–93^.

### Transmission electron microscopy analysis

Synaptosomes were centrifuged at 21,092xg for 2 min at RT and resuspended in Fixative buffer (glutaraldehyde 2.5% in 0.1M sodium cacodylate buffer) for 1h at 4°C. The samples were postfixed using 1% osmium tetroxide and 1% potassium ferrocyanide in 0.1M sodium cacodylate buffer for 1h at 4°C. After three washes in 0.1M sodium cacodylate buffer samples were dehydrated in a graded ethanol series and embedded in epoxy resin (Sigma-Aldrich). Ultrathin sections (60-70 nm) were obtained using Ultrotome V (LKB) ultramicrotome and counterstained using a saturated solution of uranyl acetate in ethanol 50% and lead citrate 2,5% in water). Sections were imaged as described in^29,86^.

### AAV stereotaxic injection

Mice were anesthetized with isoflurane (Piramal) and maintained using a veterinary vaporizer (Surgivet). AAV9-GfaABC1D-tagBFP-miR30-scramble or AAV9-GfaABC1D-tagBFP-miR30- shAckr3 (1×10^13^ vector genomes (vg)/ml, 300nl) and AAV9-hSyn-Synaptophysin-mCherry-eGFP (1×1013 vg/ml, 300nl) were injected unilaterally into the hippocampus CA1 (CA1; ML: +1.2 mm, AP: -2.0 mm, DV: −1.25 mm from bregma) and the hippocampus CA3 (CA1; ML: +2.5 mm, AP: - 2.0 mm, DV: −2.35 mm from bregma), respectively. All surgery was performed with a stereotaxic frame (Kopf) and a motorized Hamilton syringe pump (Harvard apparatus) with glass pipettes (WPI). After the incision was sutured, the mice were recovered and returned to their home cage. All injection were performed 3 weeks before sacrifice. Both males and females were included in injections.

**Table 1.**
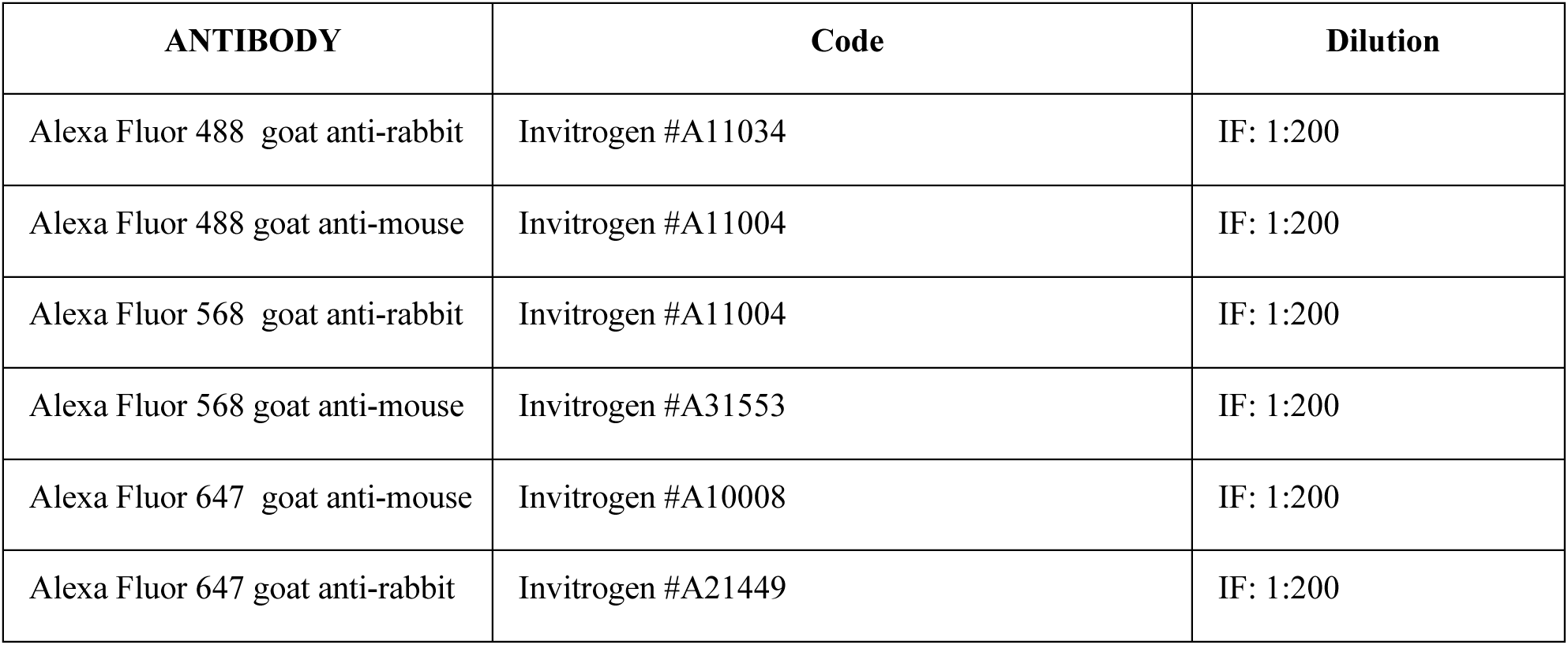
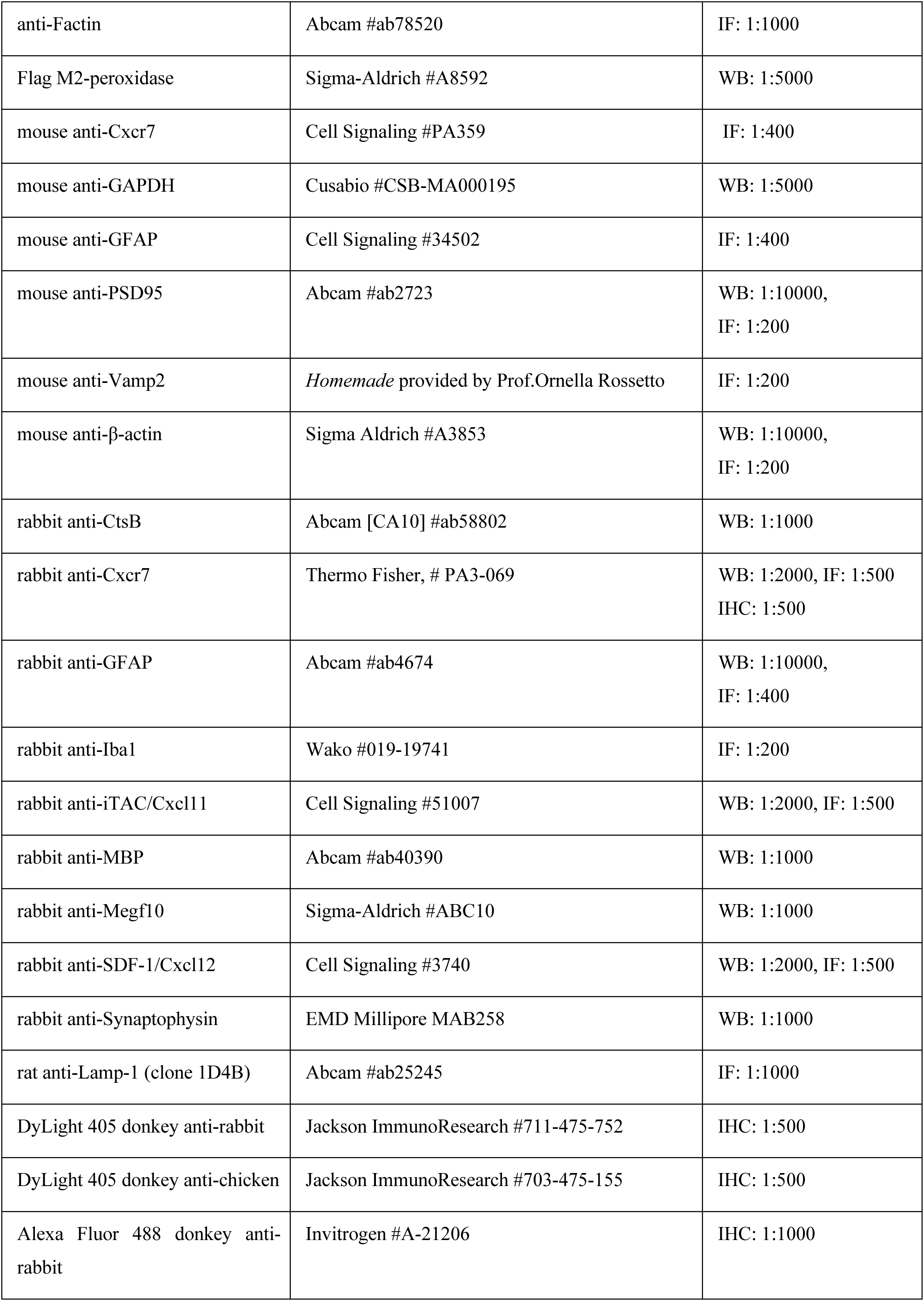
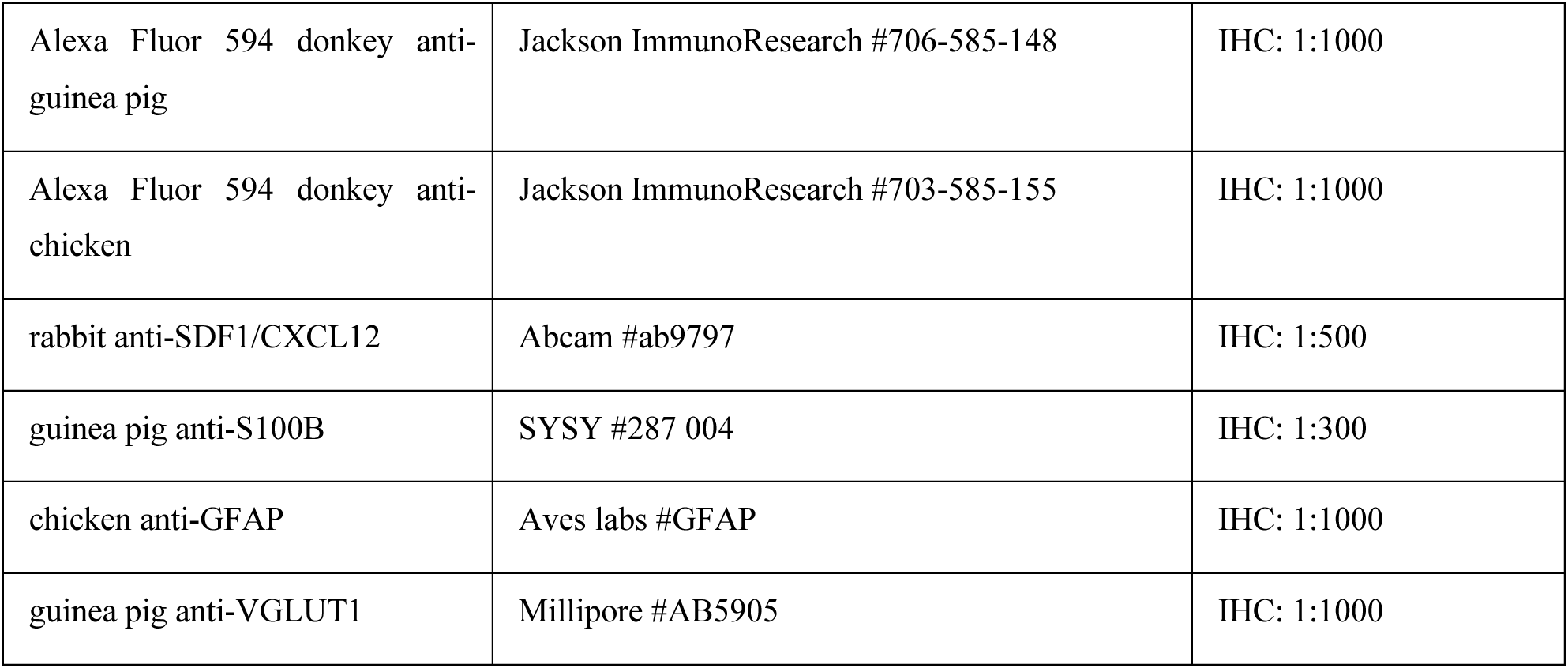
Antibodies. (IF: immunofluorescence; WB: Western-Blot; IF: immunofluorescence; IHC: immunohistochemistry)

| ANTIBODY | Code | Dilution |
| --- | --- | --- |
| Alexa Fluor 488 goat anti-rabbit | Invitrogen #A11034 | IF: 1:200 |
| Alexa Fluor 488 goat anti-mouse | Invitrogen #A11004 | IF: 1:200 |
| Alexa Fluor 568 goat anti-rabbit | Invitrogen #A11004 | IF: 1:200 |
| Alexa Fluor 568 goat anti-mouse | Invitrogen #A31553 | IF: 1:200 |
| Alexa Fluor 647 goat anti-mouse | Invitrogen #A10008 | IF: 1:200 |
| Alexa Fluor 647 goat anti-rabbit | Invitrogen #A21449 | IF: 1:200 |
| anti-Factin | Abcam #ab78520 | IF: 1:1000 |
| Flag M2-peroxidase | Sigma-Aldrich #A8592 | WB: 1:5000 |
| mouse anti-Cxcr7 | Cell Signaling #PA359 | IF: 1:400 |
| mouse anti-GAPDH | Cusabio #CSB-MA000195 | WB: 1:5000 |
| mouse anti-GFAP | Cell Signaling #34502 | IF: 1:400 |
| mouse anti-PSD95 | Abcam #ab2723 | WB: 1:10000,<br>IF: 1:200 |
| mouse anti-Vamp2 | *Homemade* provided by Prof.Ornella Rossetto | IF: 1:200 |
| mouse anti-β-actin | Sigma Aldrich #A3853 | WB: 1:10000,<br>IF: 1:200 |
| rabbit anti-CtsB | Abcam [CA10] #ab58802 | WB: 1:1000 |
| rabbit anti-Cxcr7 | Thermo Fisher, # PA3-069 | WB: 1:2000, IF: 1:500<br>IHC: 1:500 |
| rabbit anti-GFAP | Abcam #ab4674 | WB: 1:10000,<br>IF: 1:400 |
| rabbit anti-Iba1 | Wako #019-19741 | IF: 1:200 |
| rabbit anti-iTAC/Cxcl11 | Cell Signaling #51007 | WB: 1:2000, IF: 1:500 |
| rabbit anti-MBP | Abcam #ab40390 | WB: 1:1000 |
| rabbit anti-Megf10 | Sigma-Aldrich #ABC10 | WB: 1:1000 |
| rabbit anti-SDF-1/Cxcl12 | Cell Signaling #3740 | WB: 1:2000, IF: 1:500 |
| rabbit anti-Synaptophysin | EMD Millipore MAB258 | WB: 1:1000 |
| rat anti-Lamp-1 (clone 1D4B) | Abcam #ab25245 | IF: 1:1000 |
| DyLight 405 donkey anti-rabbit | Jackson ImmunoResearch #711-475-752 | IHC: 1:500 |
| DyLight 405 donkey anti-chicken | Jackson ImmunoResearch #703-475-155 | IHC: 1:500 |
| Alexa Fluor 488 donkey anti-rabbit | Invitrogen #A-21206 | IHC: 1:1000 |
| Alexa Fluor 594 donkey anti-guinea pig | Jackson ImmunoResearch #706-585-148 | IHC: 1:1000 |
| Alexa Fluor 594 donkey anti-chicken | Jackson ImmunoResearch #703-585-155 | IHC: 1:1000 |
| rabbit anti-SDF1/CXCL12 | Abcam #ab9797 | IHC: 1:500 |
| guinea pig anti-S100B | SYSY #287 004 | IHC: 1:300 |
| chicken anti-GFAP | Aves labs #GFAP | IHC: 1:1000 |
| guinea pig anti-VGLUT1 | Millipore #AB5905 | IHC: 1:1000 |

### Immunocytochemistry and immunofluorescence

1x10^5^ cells/well primary striatal astrocytes and iHA were plated onto 12mm glass coverslips (Thermo-Scientific) in 24-well plates coated with Poly-lysine (Sigma) or in 50% laminin 111 (LN111) and 50% LN211 (Biolamina). After treatments, fixation was performed with 4% w/v Paraformaldehyde (PFA) (Merck/Sigma), pH 7.4, for 20 min at RT. Cells were permeabilized with 0.1% v/v Triton® X-100 in DPBS for 20 min and successively incubated for 60 min with blocking solution (5% v/v FBS in DPBS) to saturate non-specific sites. Primary antibodies diluted in blocking solution were then added to the permeabilized cells overnight (O/N) at 4°C in a humidity chamber. The day after, after washing of the primary antibody solution, secondary antibodies diluted in blocking solution were incubated for 1h at RT. Cover glasses were then washed three times in PBS and were mounted on a microscope slide (Thermo Scientific) using Mowiol (Calbiochem) supplied with DAPI.

For the staining of the human hippocampi, paraffin-embedded sections (8 μm) were cut using a microtome (Thermo Scientific). To remove paraffin wax, 8 μm slices were washed in xylene and rehydrated in 100% ethanol. Slices were treated for 15 min with a quenching solution containing 50mM NH4Cl in phosphate buffer saline (PBS) for immunofluorescence analysis. Slices were then washed in TBS-T and incubated at 90 °C for 15 min with a citrate buffer (10mM sodium citrate and 0.1M citric acid dissolved in distilled water; pH 7.0) to perform epitope retrieval. Subsequently, slices were saturated for 1h at RT in a blocking solution containing 1% BSA, 15% goat serum, 0.25% gelatine, 0.20% glycine, and 0.5% Triton X-100. Slices were then incubated O/N at 4 °C with primary antibodies diluted in blocking solution The following day, slices were washed in TBS-T and incubated for 1h at RT with a mixture of the secondary antibodies Upon washes in TBS- T, slices were incubated for 5 min with Hoechst (1:10,000; Invitrogen) to visualize the nuclei and mounted with Mowiol (Calbiochem).

For *in vivo* experiments, mice were anesthetized using avertin (0.2g/ml, Sigma) and transcardially perfused with phosphate-buffered saline (PBS) followed by 4% paraformaldehyde (PFA, Bio- Solution). The brains were post-fixed for 12h in 4% PFA at 4°C, then cryoprotected in 30% sucrose (Sigma)/PBS for 24h at 4°C. After the brain samples were embedded and frozen in OCT compound (Leica), consecutive brain sections (30 μm) were collected in PBS. Each section was subsequently treated with a blocking solution (4% bovine serum albumin, 0.3% Triton X-100 in PBS) for 1 hour at room temperature (RT) and incubated with the primary antibodies in blocking solution for at least 24h at 4°C: The sections were rinsed five times with PBST (0.1% Tween 20 in PBS, Sigma) and incubated with secondary antibodies for 12h at 4°C. After multiple washes in PBST, the sections were coverslipped with an antifade mounting medium (Vectorshield).

### Gene Ontology (GO) enrichment analysis

Molecular function enrichment analysis was performed using GO ^94^ categories. The list of genes that abolish the ability of astrocytes to internalize synaptosome (red hits) and those that elicited the opposite (green hits) were analyzed testing the enrichment analysis of MF GO terms by using the ClusterProfiler R package ^95,96^. MF GO terms with an adjusted p-values lower than 0.05 were considered for data interpretation and the top 10 ranked terms with the lowest p-value were represented in figures. Figures were generated with dotplot and cnetplot functions. All the analyses have been performed with R version 4.1.2 and Bioconductor version 3.14 ^97^.

### Transcriptomic data from public datasets

Expression matrices of publicly available dataset have been downloaded from GEO database (details of sequencing and experimental procedures on GSE52564 series, Zhang Y, Chen K, Sloan SA, Bennett ML et al. An RNA-sequencing transcriptome and splicing database of glia, neurons, and vascular cells of the cerebral cortex ^33^. The data have been filtered by the 8 genes responsible for the expression of chemokine receptors, and a log expression matrix of these eight gene has been plotted thorough R package ComplexHeatmap (version 2.18.0), and column are clustered through the “euclidean” method. All the analyses have been performed with R version 4.3.2 and Bioconductor version 3.18.

### Immunoprecipitation and pull-down Assay

Flag-Human CXCR7(ACKR3)-Tango (Plasmid #66265, Addgene) immune-purification was performed as described in ^98^. Briefly, transfected Hek293FT cells were lysed in an appropriate volume of RIPA buffer. Lysates were centrifuged for 30 min at 14,000×g and subsequently incubated with anti-Flag M2 agarose resin beads O/N at 4 °C with gentle agitation. Immuno-purified Flag- ACKR3 was then incubated O/N at 4°C with 60 μg of intact synaptosomes. The following day, anti-Flag beads were centrifuged for 1 min at 9000xg and washed several times with lysis buffer. Samples were then resolved by SDS-PAGE and Western Blot.

### Pip Strips

PIP Strips (Thermo Scientific) were incubated with blocking buffer 1X Tris-buffered saline (TBS), 0.1% Tween-20, 3% BSA for 1h at RT. Then, recombinant murine SDF-1α-CXCL12, (Peprotech) was added at 0.1μg/ml in blocking buffer and incubated for 1h at RT. Strips were washed with TBS, 0.1% Tween-20 (TBS-T) and bound SDF-1α-CXCL12 chemokine was detected with anti-murine SDF-1α-CXCL12 antibody (diluted 1:2000, Peprotech) followed by an HRP-conjugated anti-rabbit antibody (diluted 1:16000, Abcam, Cambridge, Massachusetts, USA). Strips were developed with Immobilon Forte Western HRP substrate (Millipore) and the VWR Imager Chemi Premium.

### Western Blot Analysis

Samples were lysed in an appropriate volume of RIPA buffer (20mM Tris-HCl pH 7.5, 150mM NaCl, 1mM EDTA, 0.5mM sodium pyrophosphate (Na4P2O7), 1mM β-glycerophosphate (C_3_H_7_Na_2_O_6_P), 1mM sodium orthovanadate (Na_3_VO_4_) containing 1% protease inhibitor cocktail (Sigma-Aldrich). Protein concentration was determined through the Pierce BCA Protein Assay Kit following the manufacturer’s instructions (Thermo Scientific) and 25 μg of each sample was prepared for SDS-PAGE. Electrophoresis was performed using ExpressPlus PAGE precast gels 4- 20% (GeneScript). After electrophoresis, protein samples were transferred to PVDF membranes (Bio-Rad) through a Trans-Blot TurboTM Transfer System (Bio-Rad) in semi-dry conditions, with 1x transfer buffer (Bio-Rad) at 25V for 20 min. Membranes were incubated with 5% milk (blocking buffer) diluted in TBST at RT for 1h and then incubated for 1h at RT with appropriate horseradish peroxidase (HRP)-conjugated secondary antibodies (Invitrogen). The visualization of the signal was conducted using Immobilon Forte Western HRP substrate (Millipore) and the VWR Imager Chemi Premium. Images were acquired in .tiff format and processed with the ImageJ software to quantify the total intensity of each single band.

For dot-blot assay, 10-20-30 μg of synaptosomes were spotted onto nitrocellulose membranes (Bio-Rad) and allowed to dry. Membranes were incubated with blocking buffer at RT for 1h. Then, recombinant Cxcl12was added at 0.1 μg/ml diluted in blocking buffer and incubated for 1h at RT. Proteins were identified by the appropriate primary antibodies against Vamp2, PSD95 and Cxcl12 antibodies. Then the membrane was incubated for 1h at RT with appropriate horseradish peroxidase (HRP)-conjugated secondary antibodies (Invitrogen). The visualization of the signal was conducted using Immobilon Forte Western HRP substrate (Millipore) and the VWR Imager Chemi Premium. Images were acquired in .tiff format and processed with the ImageJ software to quantify the total intensity of each single band.

### Human Samples and clinical data

All procedures were carried out in accordance to the Declaration of Helsinki. Samples were anonymous to the investigators and used in accordance with the directives of the Committee of the Ministers of EU member states on the use of samples of human origin for research. Donor subjects gave their written informed consent prior to death according to regulations of the Body Donation Program of the Institute of Human Anatomy, Department of Neuroscience, University of Padova.

**Table 2.**
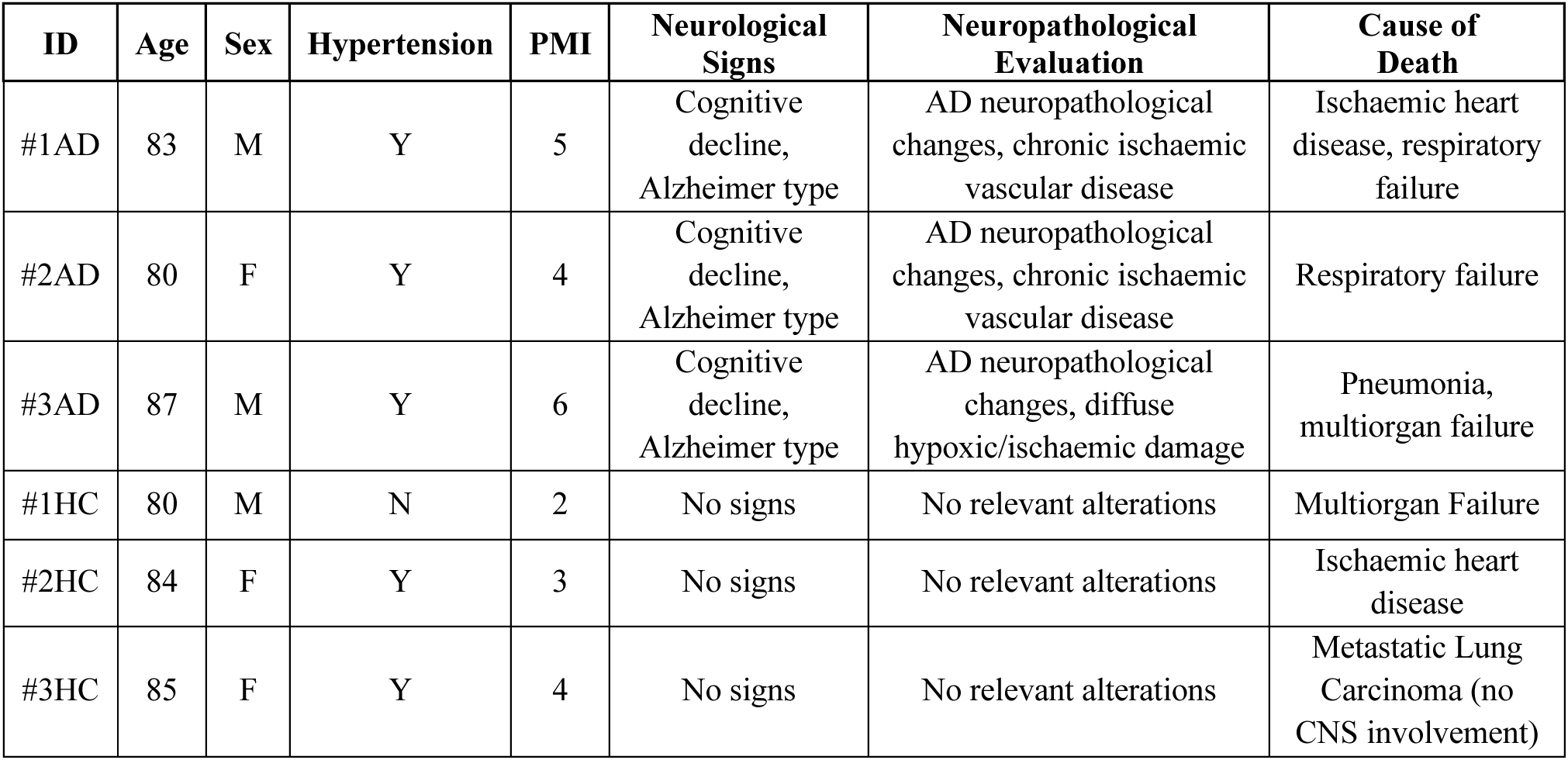
Sample demographic and clinical data of the human cases of AD and Healthy control (HC) used in this study. The table illustrates sex, age, hypertension, Post-Mortem Interval (PMI), Neurological signs, Neuropathological evaluation and Cause of Death.

| ID | Age | Sex | Hypertension | PMI | Neurological Signs | Neuropathological Evaluation | Cause of Death |
| --- | --- | --- | --- | --- | --- | --- | --- |
| #1AD | 83 | M | Y | 5 | Cognitive decline, Alzheimer type | AD neuropathological changes, chronic ischaemic vascular disease | Ischaemic heart disease, respiratory failure |
| #2AD | 80 | F | Y | 4 | Cognitive decline, Alzheimer type | AD neuropathological changes, chronic ischaemic vascular disease | Respiratory failure |
| #3AD | 87 | M | Y | 6 | Cognitive decline, Alzheimer type | AD neuropathological changes, diffuse hypoxic/ischaemic damage | Pneumonia, multiorgan failure |
| #1HC | 80 | M | N | 2 | No signs | No relevant alterations | Multiorgan Failure |
| #2HC | 84 | F | Y | 3 | No signs | No relevant alterations | Ischaemic heart disease |
| #3HC | 85 | F | Y | 4 | No signs | No relevant alterations | Metastatic Lung Carcinoma (no CNS involvement) |

### Image acquisition and analysis

For live-imaging studies in primary striatal astrocytes or iHA, images were acquired at 32-bit intensity resolution over 1024×1024 pixels through the Operetta High Content Imaging System (Perkin Elmer) or ApoTome2 (Carl Zeiss) using a 10x long working distance dry objective. eGFP- positive and pH-RODO positive areas were assessed using Harmony Software (Perkin Elmer). Phagocytic Index was calculated as total pH-RODO positive area normalized by total eGFP positive area multiplied by 100.

For post-fixation imaging of primary striatal astrocytes or iHA, images were acquired at 8-bit intensity resolution over 1024×1024 pixels, through Leica SP5 or Leica TCS STED CW SP8 confocal microscope and a HC PL FLUOTAR×40/0.70 oil objective or HC APO L 40x/0.80 W U-V-I immersion objective. After setting the threshold, the JACoP plugin (ImageJ, https://imagej.nih.gov/ij/plugins/track/jacop2.html) was applied for the quantification of co- localization between Ackr3 signal and the Lamp1 and TAMRA-positive structures, graphically displayed as the Pearson’s Correlation Coefficient.

For post-fixation imaging of primary neurons and neurons-astrocytes co-cultures, images were acquired at 8-bit intensity resolution over 1024×1024 pixels, through Leica SP5 and a HC PL FLUOTAR×40/0.70 oil objective. For PE-exposure and synapses internalization experiment, after setting the threshold, ComDet plugin (https://imagej.net/Spots_colocalization_ (ComDet)), using the following parameters: max co-localization distance (0.4 pixels for pSIVA and Duramycin and 0.2 pixels for Vamp/Psd95 experiment) and particles dimension (Vamp2/PSD95: 3 pixels and pSIVA/Duramycin: 2 pixels). 3D reconstructions were done using IMARIS software (Bitplane 9.9.1). For Cxcl12 quantification: after setting the threshold, we identified a 50 nm rectangle based on F-actin positive dendrites. The number of Cxcl12 dots were assessed by “puncta count” output using the ComDet plugin, with the following parameter: particle dimension (Cxcl12: 2 pixels) and normalized by the ROI’s dimension.

For human tissues, images were acquired at 8-bit intensity resolution over 1024 × 1024 pixel on a Leica SP5 confocal microscope using an HC PL FLUOTAR 40x/0.70 oil objective. Using ImageJ, the IntDen of ACKR3, GFAP and CXCL12 signals upon setting scale and threshold were measured. The co-localization of GFAP/ACKR3^+^ cells was defined using the JACoP plugin. For the staining of the mouse hippocampi, Confocal images of mouse brain tissue were acquired using a Nikon Ti2 AX inverted confocal microscope with a 63× (NA 1.42) objective. Imaging was performed using NSI Elements software, with consistent acquisition parameters across all samples, including 16-bit depth and 1024 × 1024 resolution. Using ImageJ, the IntDen of ACKR3, GFAP signals upon setting scale and threshold were measured. The co-localization of GFAP/ACKR3+ cells was defined using the JACoP plugin.

For *in vivo* analysis, all images were acquired using an Airyscan LSM880 confocal microscope (Zeiss) and processed with ImageJ (Fiji). For colocalization and expression analysis, each plane of the z-stack images was processed using the ImageJ Distance Analysis (DiAna) plugin ^88^. For excitatory pre-synapse phagocytosis (ExPre) analysis, mCherry-alone signals were obtained by subtracting mCherry signals from eGFP signals. The DiAna plugin was then used to assess colocalization between mCherry alone signals and tagBFP, normalized by eGFP area. All representative images were merged in the z-stack planes.

### AAV PHP.eB Injection and Behavioral tests

The production of AAV-PHP.eB from pAAV-GfaABC1D-tagBFP-miR30-scramble and pAAV- GfaABC1D-tagBFP-mir30-shAckr3 was performed and titered by the IBS (Institute for Basic Science) LSI (Life Science Institute) facility. 3-month-old 5xFAD mice and littermate wild-type mice were anesthetized and received retro-orbital injections of the virus (5.0 × 1010 GC). After 1 month, all behavioral tests were conducted and recorded.

Behavioral assessments included the open field test and Y-maze test. Briefly, to measure anxiety-like behavior, the open field test was conducted in a white box (30 cm × 30cm × 30cm) for 30 minutes. To evaluate short-term spatial memory, the novel arm recognition was conducted in a white Y-maze (30 cm × 5cm × 12cm). In the novel arm recognition test, mice were trained in the familiar arms for 15 minutes. After 30 minutes, the novel arm was opened, and the mice were allowed to explore for 5 minutes. Both male and female mice were utilized in all behavioral experiments. Behavioral data were analyzed using EthoVision software (Noldus).

### Statistical Analysis

Results are expressed as mean ± standard deviation (SD). Data were analysed with unpaired t tests or Two-way ANOVA (Gaussian distribution) respectively followed by Tukey’s multiple comparisons test. Levels of significance were defined as p ≤ 0.05, p ≤ 0.01, p ≤ 0.001. Statistical analysis was performed with Prism 9 (GraphPad).

## ABBREVIATIONS

Adeno-associated virus 9: AAV9
Alzheimer’s disease: AD
ATP-binding cassette transporter: A1 Abca1
Atypical chemokine receptor 3: Ackr3
Basal Medium Eagle: BME
Cathepsin B: CtsB
Cell division control protein 50 a: Cdc50a
Central nervous system: CNS
Cxcl12-covered synaptosomes: cS
Dulbecco’s Phosphate Buffered Saline: DPBS
Fetal bovine serum: FBS
Glial Fibrillary Acidic Protein: GFAP
Gene ontology: GO
Human iPS-derived astrocytes: iHA
Human synapsin promoter: hSYN
Immunoprecipitation: IP
Inositol 1,4,5-trisphosphate receptor type 2: Ip3r2
Luria-bertani: LB
MER proto-oncogene tyrosine kinase: Mertk
Myelin Binding Protein: MBP
Naive Synaptosomes: S
Pearson’s Correlation Coefficient: PCC
Phosphatidylethanolamine: PE
Phosphatidylserines: PS
Phosphotyrosine-binding domain containing 1: Gulp1
pHrodo-Red TM succinimidyl ester: pH-RODO
Post synapses compartment: PO
Postsynaptic density: PSD
Postsynaptic Density Protein 95: PSD95
Pre synapses compartment: PSc
siRNA Ackr3: siAckr3
siRNA Control: siControl
Synaptophysin: Syp
Transmission electron microscopy: TEM
Vesicle associated membrane protein2: Vamp2
Vesicle associated membrane protein2: Vamp2
Wild-type: WT

## AUTHOR CONTRIBUTIONS

VG conceived and designed the experiments, supervised the whole project and wrote the paper. PJ conducted the experiments in vivo, analysed the data and generated the figures. GE analysed the data and generated the figures. ML and ECA conducted the gene ontology analysis. PRG generated human ipsc-derived astrocytes. LI and KG prepared the primary culture. MS carried out the conjugation experiments. AE and AP provided the human brain samples. ECO has contributed to manuscript editing and optimization. FP generated the 3D reconstruction. FC supervised the analysis of the data obtained from human ipsc-derived astrocytes. WSC supervised the analysis of the data obtained from in vivo experiments and contributed to manuscript editing and optimization. LC conceived and designed the experiments, supervised the whole project and wrote the paper. All authors contributed to data analysis and preparation of the manuscript.

## FUNDING

This work was supported by UniPD (STARs 2019: Supporting TAlents in ReSearch) and the Italian Ministry of Health (GR-2016-02363461) to LC; the Company of Biologists and Böhringer Ingelheim Fonds to VG. This study was also supported by the Institute of Basic Science (IBS-R025-A1 to W.- S.C.) funded by the Ministry of Science and ICT, Republic of Korea

## ACKNOWLEDGEMENTS

We acknowledge and respect the University of Padova to support LC as Associate professor and the IRCCS San Camillo Hospital in Venice, Italy for scientific collaboration. We acknowledge Matthew Holt for critical discussion and scientific support.

## CONFLICT OF INTEREST

The authors declare no conflicts of interest.

## SUPPLEMENTARY FIGURES

**Supplementary 1.**
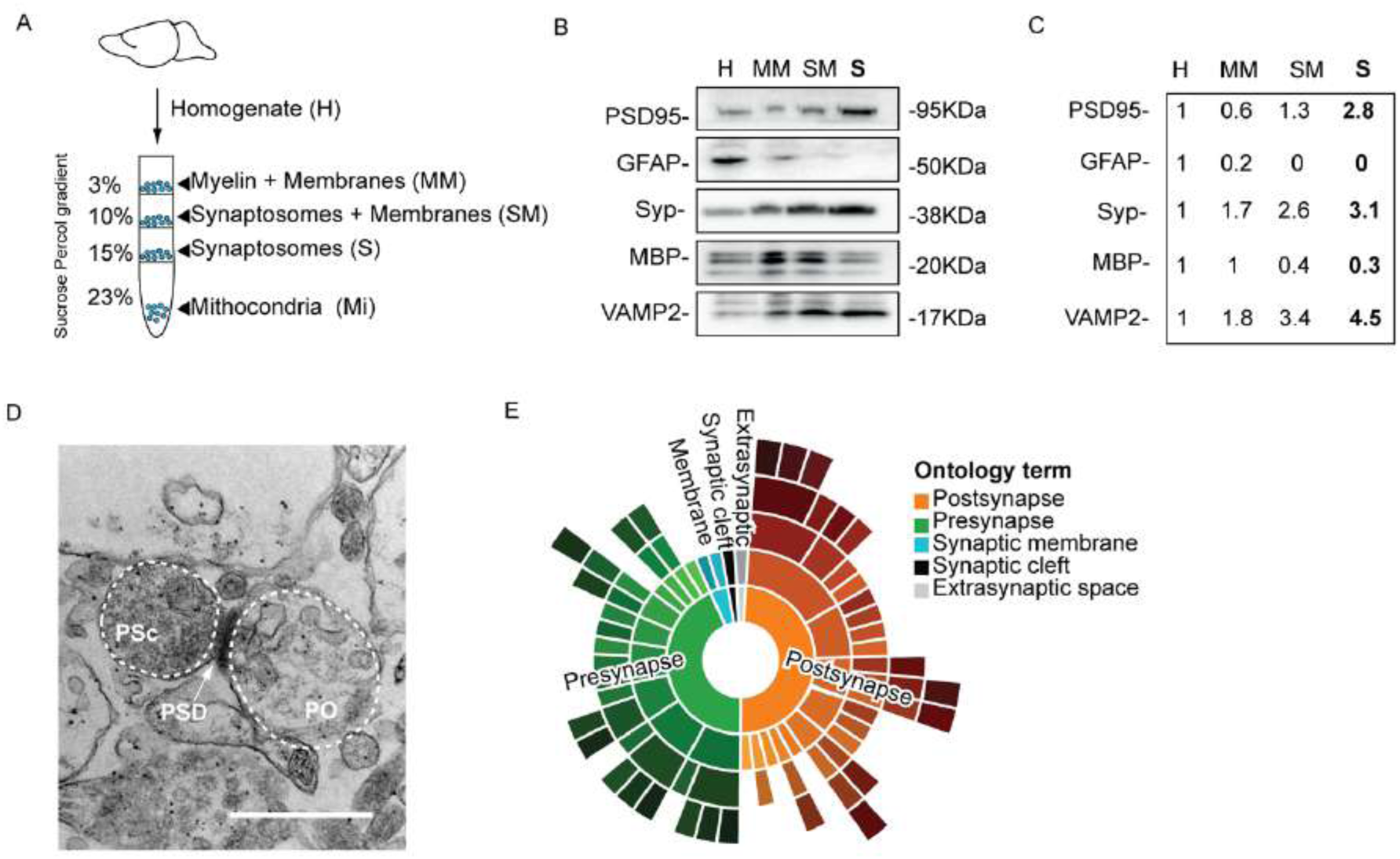
Synaptosomes purification and characterization. **A.** Graphical overview of synaptosome purification using Percoll gradient. Brain Homogenate (H), Myelin+Membrane (MM), Synaptosomes+Membranes (SM), pure synaptosomes (S) and Mitochondria (Mi). **B.** Western blot analysis of the fraction identity using specific antibodies: anti- PSD95 (post-synaptic), anti-GFAP (astrocytes), anti-Synaptophysin (pre-synaptic), anti-MBP (myelin) and anti-Vamp2 (pre-synaptic). **C.** Quantification of the band intensity of the marker relative to homogenate using ImageJ software. **D.** Representative TEM images of purified synaptosomes. The synapse comprises a presynaptic terminal (PSc), an electron-dense postsynaptic density (PSD) membrane showing the features of PSc and a postsynaptic terminal (PO). Scale bar 250 nm. **E.** Proteomic characterization and data analysis using SynGO of whole-brain purified synaptosomes. Ontology terms are visualized as ‘sunburst plots’ for cellular components and colour coded as indicated in the legend.

**Supplementary 2.**
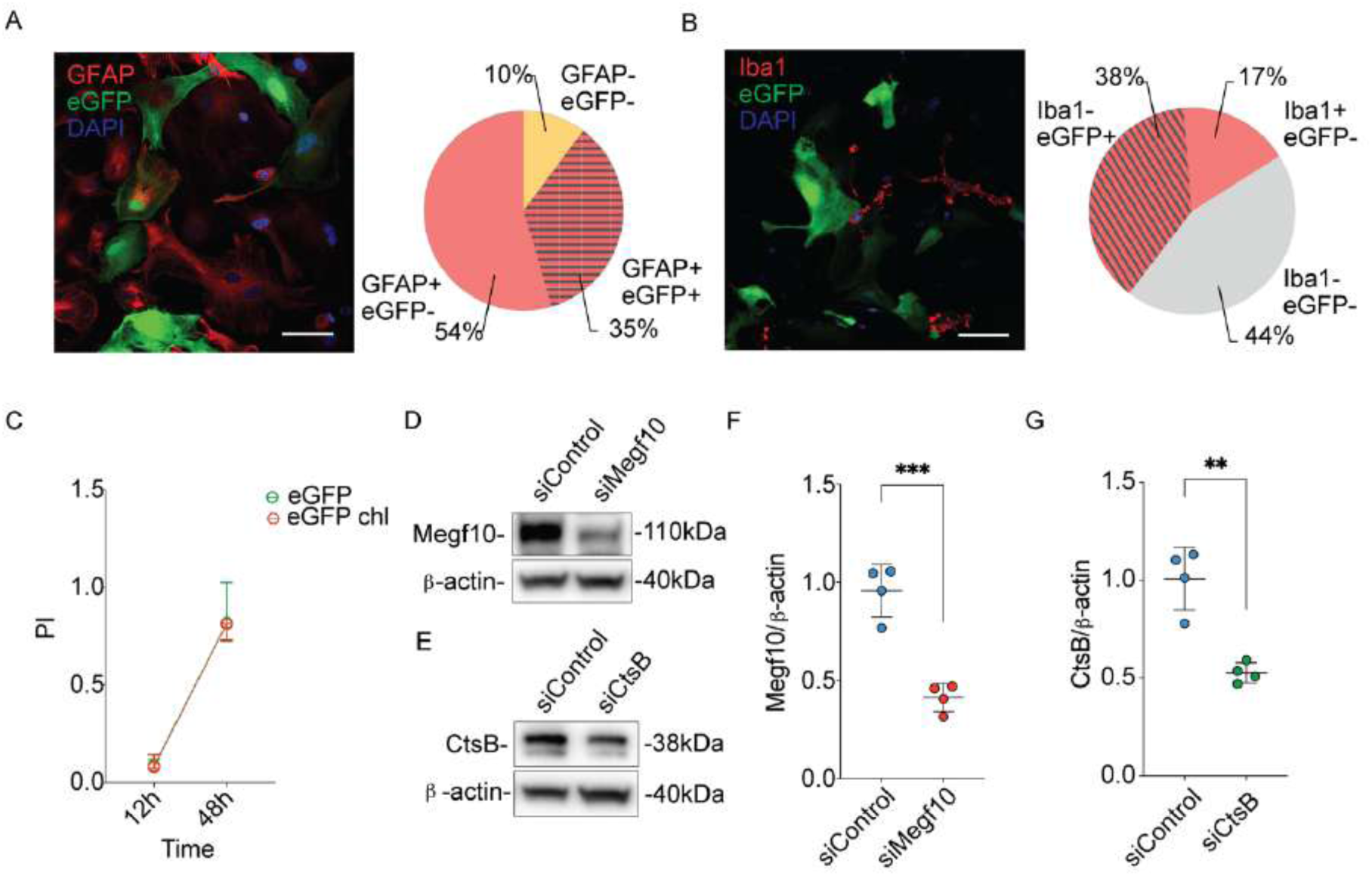
siRNA screening set up. **A.** Representative fluorescence microscopy image and relative quantification of GFAP^+^/eGFP^-^ (non-transfected astrocytes), GFAP^-^/eGFP^-^ (non-transfected, non-astrocytic cells, GFAP^+^/eGFP^+^ cells (transfected astrocytes), GFAP^-^/eGFP^+^ (transfected, non-astrocytic cells). Cells were transfected with eGFP-encoding plasmid (green) and stained for GFAP (red). Scale bar 50 µm. **B.** Representative fluorescence microscopy images and relative quantification of Iba1^-^/eGFP^+^ (transfected, non-microglial cells), Iba^-^/eGFP^-^ (non-transfected, non-microglia cells) Iba1^+^/eGFP^-^ (non-transfected microglia), Iba1^+^/eGFP^+^ (transfected microglia). Cells were transfected with GFP-encoding plasmid (green) and stained for Iba1 (red). Scale bar 50 µm. **C.** PI of primary astrocytes transfected with eGFP-encoding plasmid with (orange) or without (green) liposomal chlodronate treatment at 12h and 48h upon pH-RODO synaptosomes treatment. **D.** Western blot analysis of primary astrocytes transfected with siControl and siMegf10. **E.** Western blot analysis of primary astrocytes transfected with siControl and siCtsB. **F.** Quantification of Megf10 band of siControl and siMegf10 astrocytes. Band intensity quantification was performed using ImageJ and normalized to β-actin (Control vs. Megl10 p=0.0004); N=4 biological replicates. **G.** Quantification of CtsB band of siControl and siCtsB astrocytes. Band intensity analysis was performed using ImageJ and normalized to β-actin (Control vs. CtsB p=0.0012); N=4 biological replicates. Statistical analysis in F and G was performed using unpaired t tests.

**Supplementary 3.**
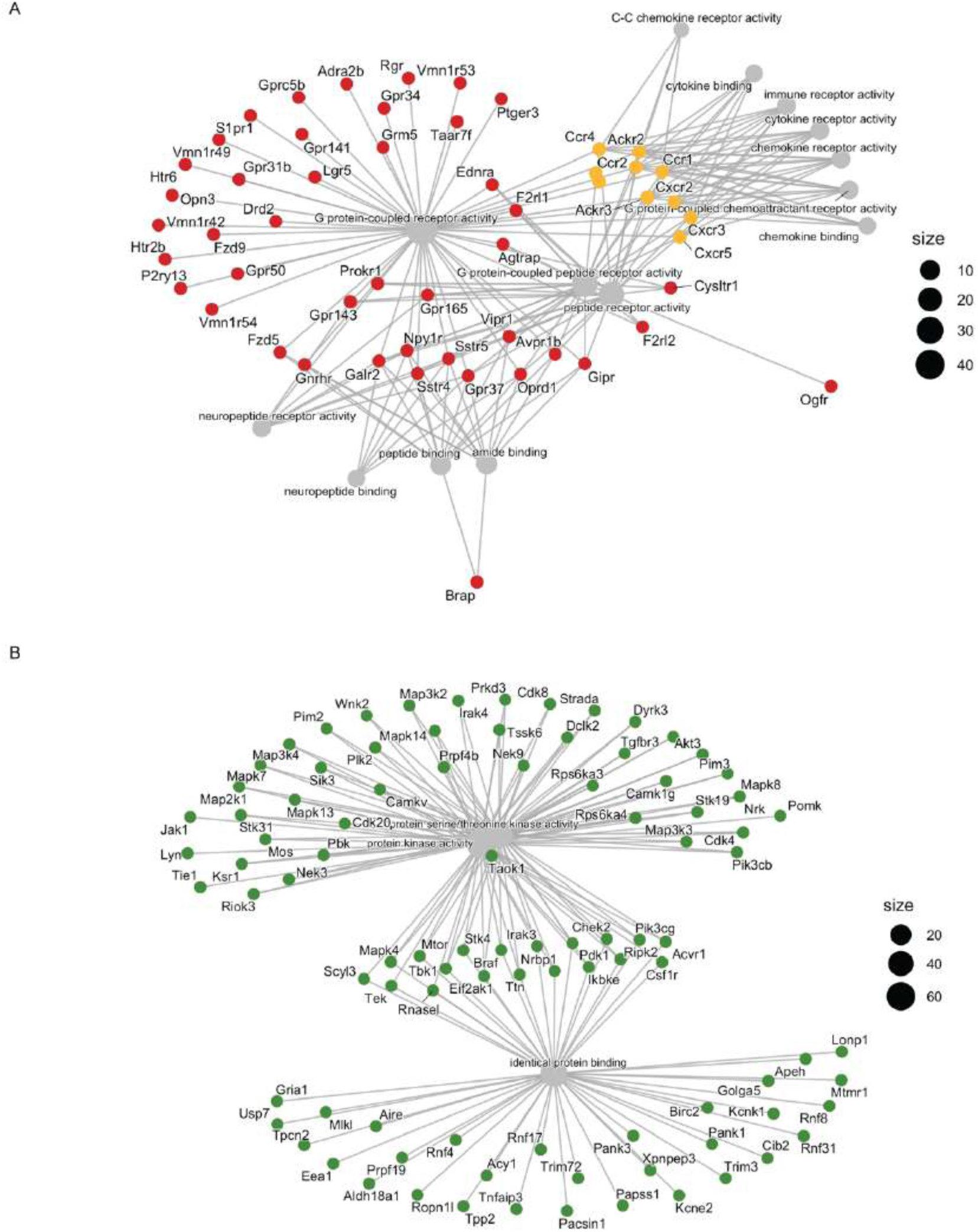
Molecular Functional enrichment analysis of target genes. **A.** Cnetplot diagram showing the pathway-gene network of red hits. The linkages of genes and biological categories (grey) are represented as a network. Eight chemokine receptors, which are top hits: Cxcr5, Ackr3, Cxcr2, Cxcr3, Ccr1, Ccr2, Ccr4 and Ackr2, are highlighted in orange. **B.** Cnetplot diagram showing the pathway-gene network of green hits. The linkages of genes and biological categories (grey) are represented as a network.

**Supplementary 4.**
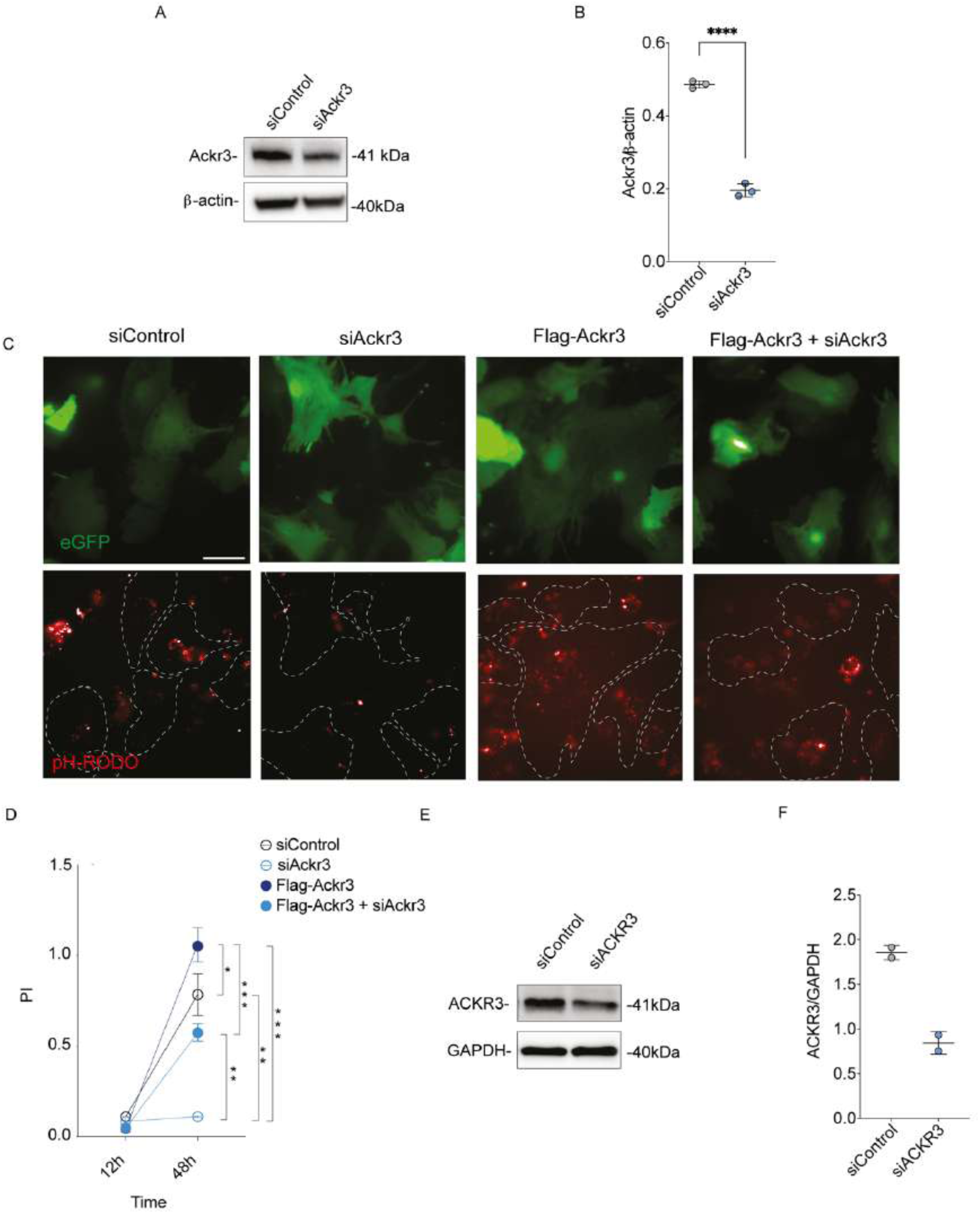
Endogenous Ackr3 modulates synaptosomes engulfment in primary astrocytes. **A.** Western blot analysis of primary astrocytes transfected with siControl and siAckr3. **B.** Quantification of Ackr3 band intensity was performed using ImageJ and normalized to β-actin (Control vs. Ackr3 p=0.0011); N=4 biological replicates. Statistical analysis was performed using unpaired t tests. **C.** Representative fluorescence images of primary astrocytes transfected with siControl, siAckr3, Flag-Ackr3 plasmid or Flag-Ackr3+ siAckr3 together with eGFP-encoding plasmid followed by pH-RODO synaptosome treatment (48h). Scale bar 50 µm. **D.** PI of primary astrocytes transfected with siControl, siAckr3, Flag-Ackr3 plasmid or Flag-Ackr3+siAckr3 together with eGFP-encoding plasmid and followed by pH-RODO synaptosome treatment (12h and 48h) (48h: Control vs. Flag-Ackr3 p=0.0019; Control vs. siAckr3 p=0.0017; siAckr3 vs. Flag-Ackr3 p=0.0011; siAckr3 vs. Flag-Ackr3+siAckr3 p=0.0019; Flag-Ackr3 vs. Flag-Ackr3+siAckr3 p=0.0004). N=3 biological replicates. Statistical analysis was performed using Two-Way ANOVA with Tukey’s multiple comparison test. **E.** Western blot analysis of iHA lysates transfected with siControl and siAckr3. **F.** Quantification of Ackr3 band intensity was performed using ImageJ and normalized to GAPDH; N=2 biological replicates.

**Supplementary 5.**
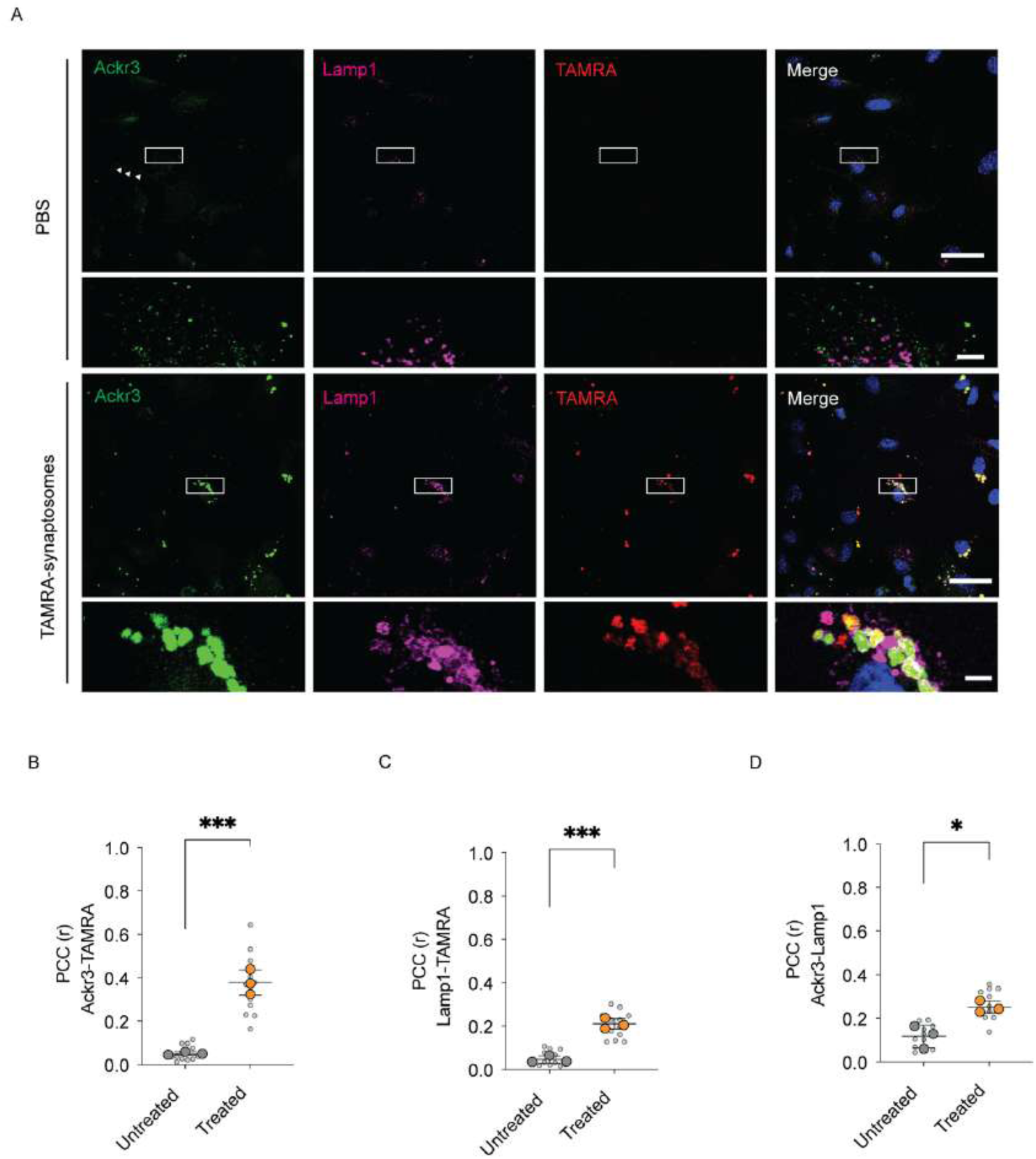
Ackr3-synaptosome complex relocates to the endo-lysosomal pathway. **A.** Representative z-stack confocal images of primary astrocytes under basal conditions or upon the treatment with TAMRA-synaptosomes (red). Cells were stained with anti-Ackr3 (green) and anti- LAMP1 (purple) antibodies. Scale bar 20 μm; insets 5 μm. **B.** Pearson’s correlation coefficient of Ackr3/TAMRA co-localization (untreated vs. treated p=0.0002; N=12 images per condition were analysed; N=3 biological replicates. **C.** Pearson’s correlation coefficient of TAMRA/Lamp1-positive compartment co-localization (untreated vs. treated p=0.0002) N=12 images per condition were analysed; N=3 biological replicates. **D.** Pearson’s correlation coefficient of Ackr3/Lamp1-positive compartment co-localization (untreated vs. treated p=0.0019) N=12 images per condition were analysed; N=3 biological replicates. Statistical analysis was performed using unpaired t tests. Statistical analysis in B-C-D was performed using unpaired t tests.

**Supplementary 6.**
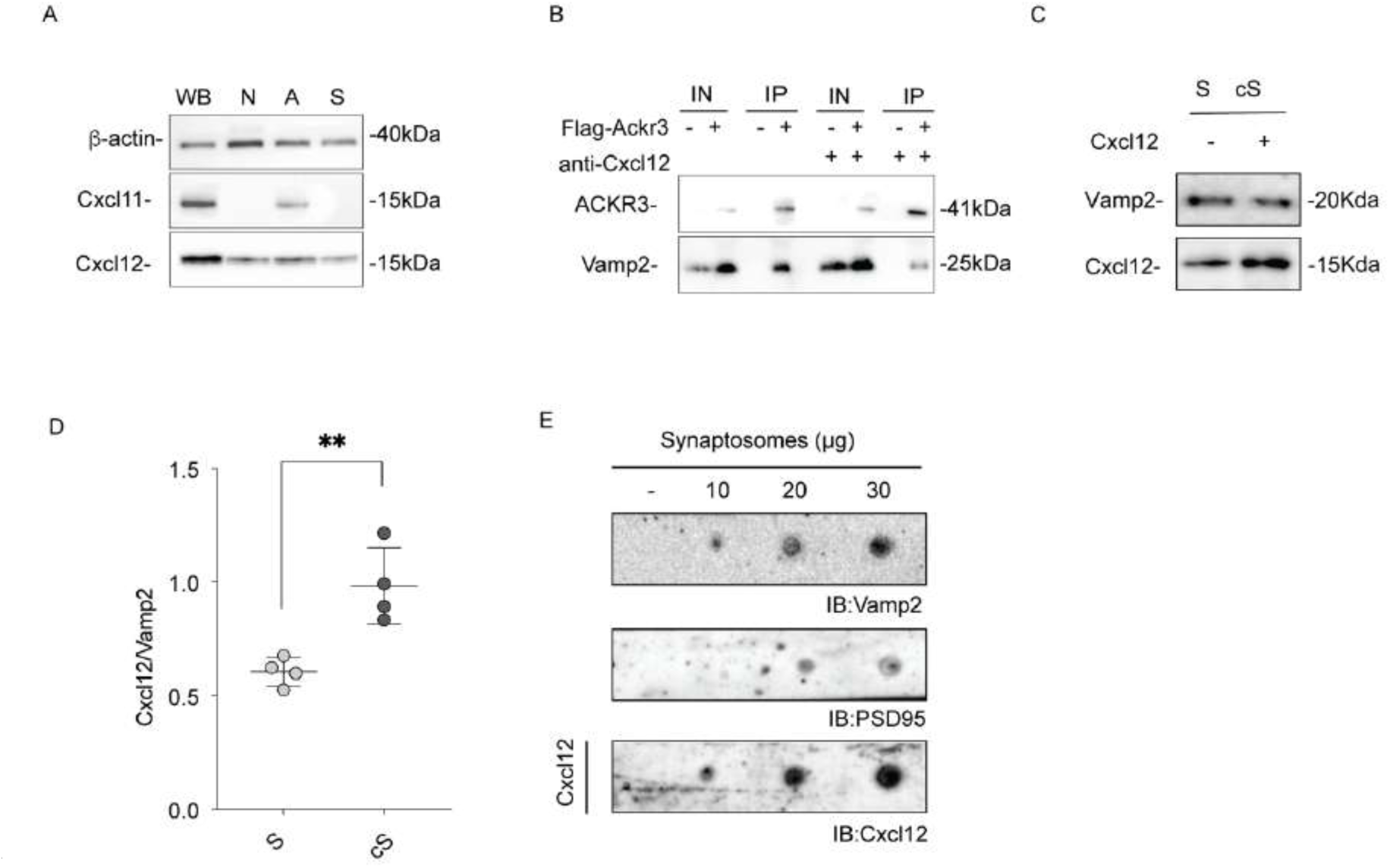
Biochemical evaluation of Cxcl12-synaptosomes binding. **A.** Western blot analysis of whole brain (WB), primary neurons (N), primary astrocytes (A) and synaptosomes (S) lysates. **B.** Western blot analysis of pull-down assay using purified Flag-Ackr3 as bait and intact synaptosomes as prey. Synaptosomes were incubated or not with anti-Cxcl12 antibody. Bound synaptosomes were detected using Vamp2. **C.** Western blot analysis of S and cS synaptosomes. **D.** Quantification of Cxcl12 band intensity normalized to Vamp2 using ImageJ software (untreated vs. treated p=0.0012), N=4 biological replicates. Statistical analysis was performed using unpaired t tests. **E.** Immuno-dot blot analysis on increasing concentration of synaptosomes spotted onto a nitrocellulose membrane and incubated with recombinant Cxcl12. Synaptosomes were detected using anti-VAMP2 and anti-PSD95 and the chemokine using anti- Cxcl12.

**Supplementary 7.**
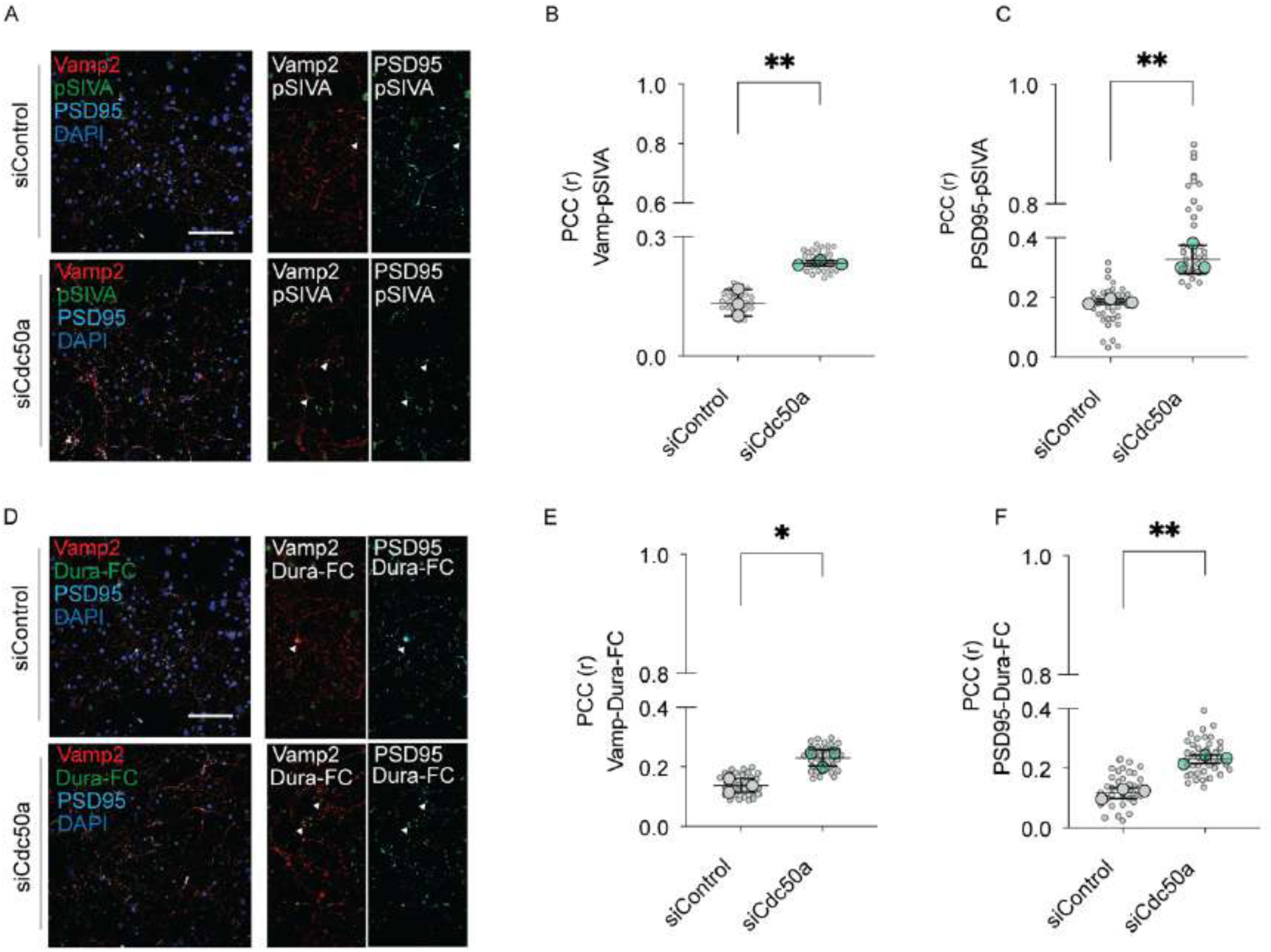
Cdc50a downregulation causes an increased exposure of PE at the synapse. **A.** Representative z-stack confocal images of primary cortical neurons transfected with siRNA Control and siRNA against Cdc50a. Cells were stained using pSIVA (green), anti-Vamp2 (red) and anti-PSD95 (cyano) antibodies. Scale bar 20 μm; insets 5 μm. **B.** PCC of Vamp2/pSIVA co- localization (siControl vs. siCdc50a p=0.0073) N=18 images per condition were analysed; N=3 biological replicates. Statistical analysis was performed using unpaired t tests. **C.** PCC of PSD95/pSIVA co-localization (siControl vs. siCdc50a p=0.0070) N=18 images per condition were analysed; N=3 biological replicates. Statistical analysis was performed using unpaired t tests. **D.** Representative z-stack confocal images of primary cortical neurons transfected with siRNA Control and siRNA against Cdc50a. Cells were stained with DuramycinLC (green), anti-Vamp2 (red) and anti-PSD95 (cyano) antibodies. Scale bar 20 μm; insets 5 μm; **E.** PCC of Vamp2/DuramycinLC co- localization (siControl vs. siCdc50a p=0.0114; N=18 images per condition were analysed; N=3 biological replicates. Statistical analysis was performed using unpaired t tests. **F.** PCC of PSD95/DuramycinLC co-localization (siControl vs. siCdc50a p=0.0010; N=18 images per condition were analysed; N=3 biological replicates. Statistical analysis was performed using unpaired t tests.

**Supplementary 8.**
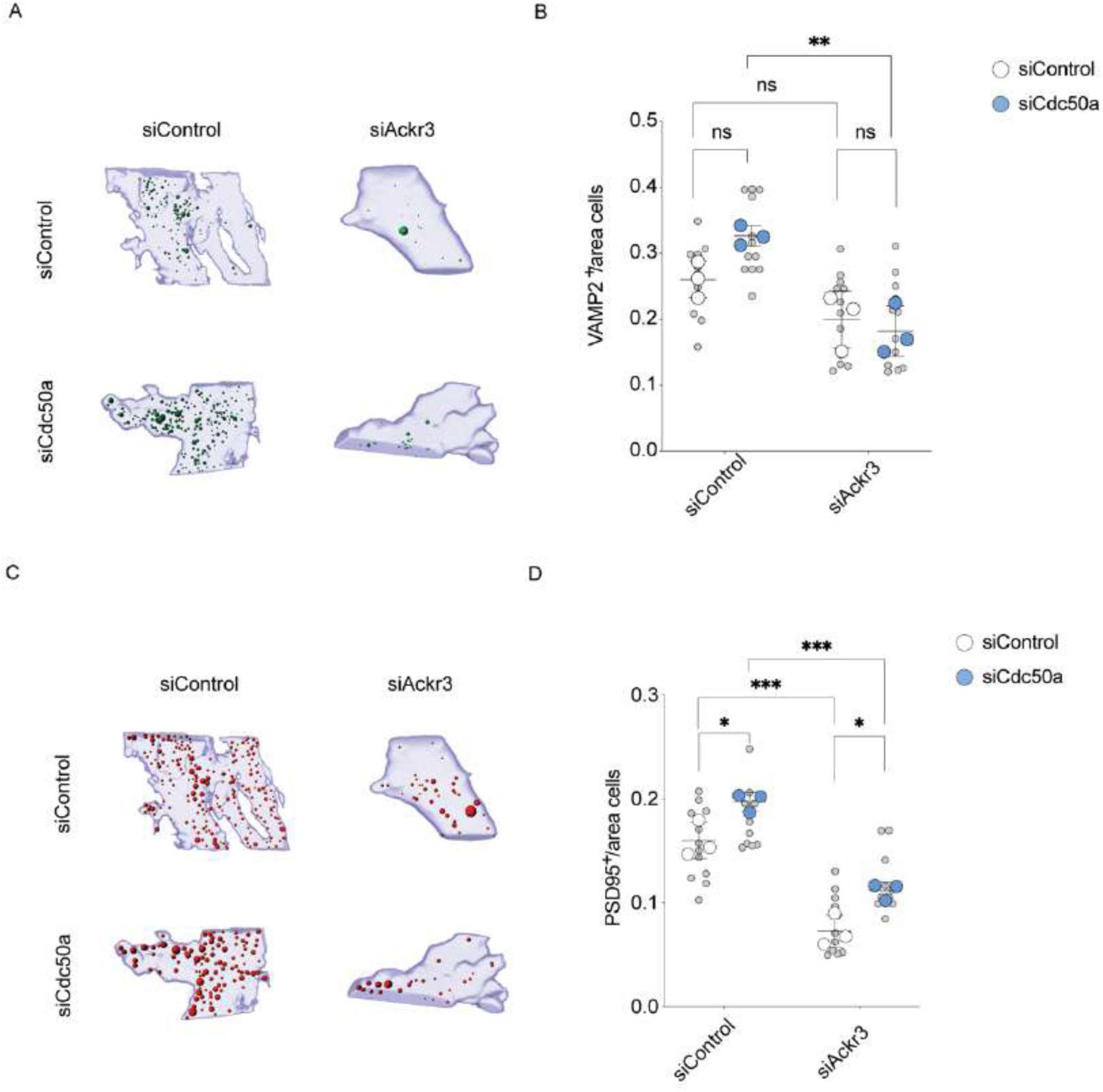
3D reconstruction of Ackr3-mediated synaptic removal. **A.** Representative 3d reconstruction of Vamp2+ synaptic terminals (green) internalized by primary astrocytes (blue). Neurons were transfected with siControl or siCdc50a. Astrocytes were transfected with siControl or siAckr3. Scale bar 20μm, inset scale bar 5 μm. **B.** Number of Vamp2+ synaptic terminals within the astrocyte area (siControl astrocytes/siControl neurons vs. siControl astrocytes/siCdc50a neurons p=0.1413; siControl astrocytes/siControl neurons vs. siAckr3 astrocytes/siControl neurons p=0.1850; siControl astrocytes/siCdc50a neurons vs. siAckr3 astrocytes/siCdc50a neurons p=0.0029; siAckr3 astrocytes/siCdc50a neurons vs. siAckr3 astrocytes/siCdc50a neurons p=0.6782). N=3 biological replicates. Statistical analysis was performed using Two-Way ANOVA with Tukey’s multiple comparison test. **C.** Representative 3d reconstruction of PSD95+ synaptic terminals (red)internalized by primary astrocytes (blue). Neurons were transfected with siControl or siCdc50a. Astrocytes were transfected with siControl or siAckr3. Scale bar 20 μm, inset scale bar 5 μm. **D.** Number of synaptic terminals (PSD95-positive) within the astrocyte area (siControl astrocytes/siControl neurons vs. siControl astrocytes/siCdc50a neurons p=0.0329; siControl astrocytes/siControl neurons vs. siAckr3 astrocytes/siControl neurons p=0.0002; siControl astrocytes/siCdc50a neurons vs. siAckr3 astrocytes/siCdc50a neurons p=0.0002; siAckr3 astrocytes/siCdc50a neurons vs. siAckr3 astrocytes/siCdc50a neurons p=0.0274). N=3 biological replicates. Statistical analysis was performed using Two-Way ANOVA with Tukey’s multiple comparison test.

**Supplementary 9.**
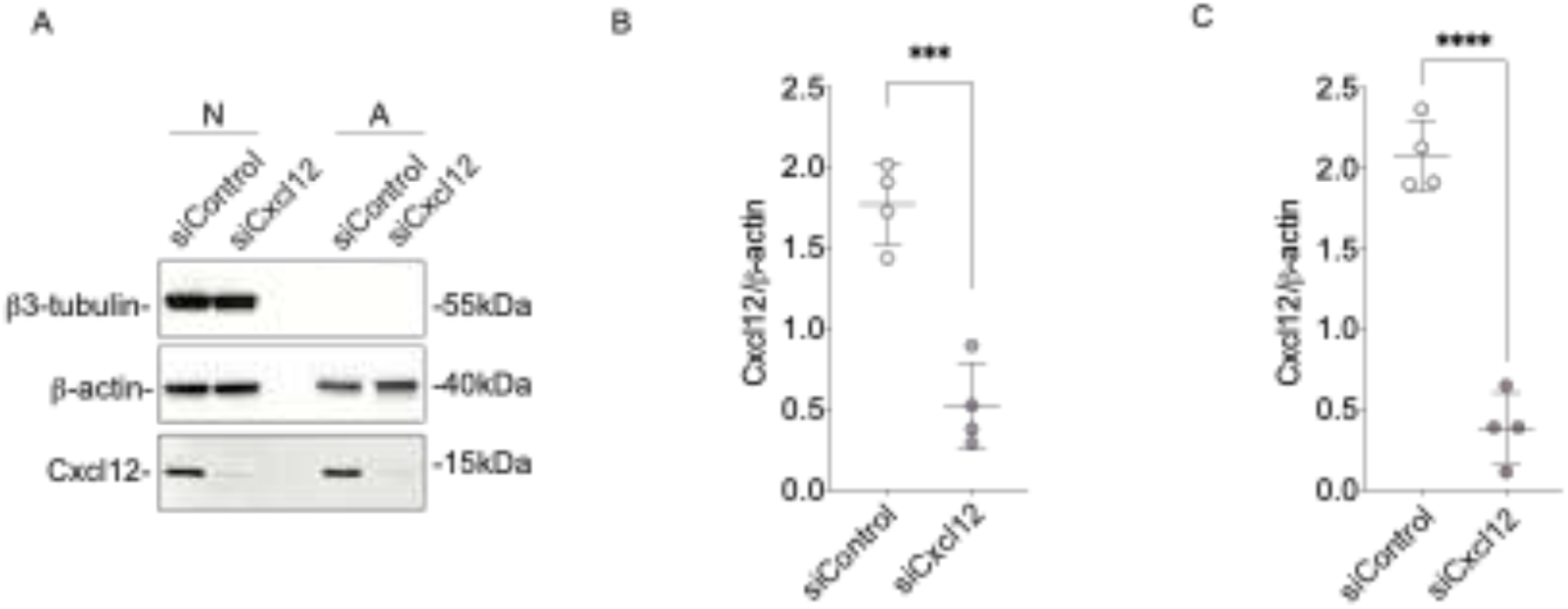
Biochemical evaluation of Cxcl12 downregulation in primary cultures. **A.** Western blot analysis of primary neurons (N) and primary astrocytes (A) transfected with siControl and siCxcl12. **B.** Quantification of siControl and siCxcl12 neurons. Band intensity analysis was performed using ImageJ and normalized to β-actin (siControl vs. siCxcl12 p=0.0011); N=4 biological replicates. **C.** Quantification of siControl and siCxcl12 astrocytes. Band intensity analysis was performed using ImageJ and normalized to β-actin (siControl vs. siCxcl12 p=0.0005); N=4 biological replicates. Statistical analysis in B and C was performed using unpaired t tests.

**Supplementary 10.**
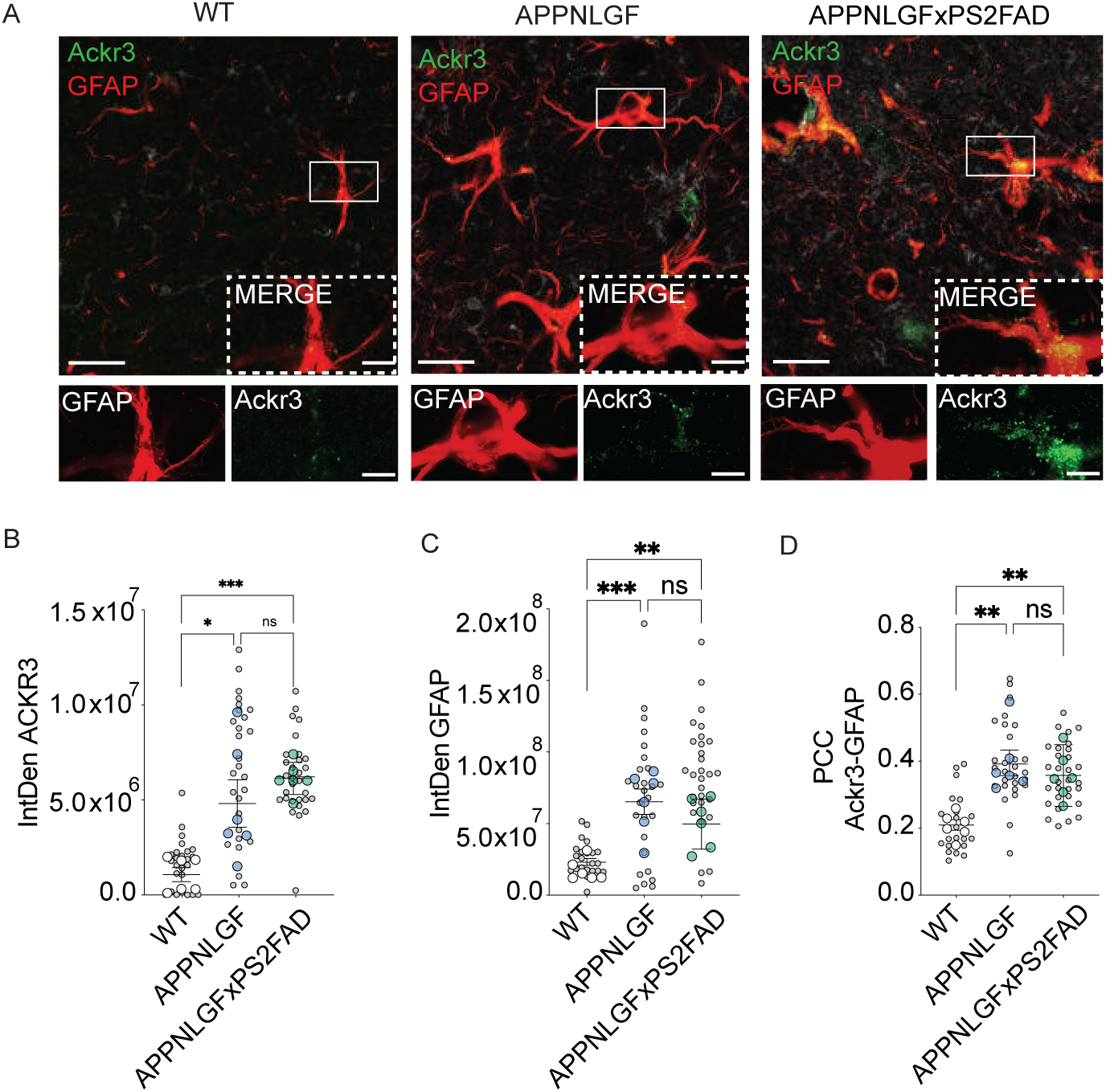
Ackr3 expression in AD mice model. **A.** Representative confocal images for ACKR3 (green), GFAP (red) in CA1 hippocampus of 9- month-old APPNLGF, APPNLGFxPS2FAD and WT mice. Scale bar 50 μm, inset 20 μm. **B.** Quantitative analysis of Ackr3 expression in the hippocampus of APPNLGF, APPNLGFxPS2FAD and WT mice; N=6 samples for WT, APPNLGF and APPNLGFxPS2FAD 9-month-old mice. Statistical analysis was performed using one-way ANOVA. **C.** Quantitative analysis of Ackr3 expression in the hippocampus of APPNLGF, APPNLGFxPS2FAD and WT mice; N=6 samples for WT, APPNLGF and APPNLGFxPS2FAD 9-month-old mice. Statistical analysis was performed using one-way ANOVA. **D.** Quantitative analysis of ACKR3 co-localizing with GFAP-positive astrocytes in the hippocampus of APPNLGF, APPNLGFxPS2FAD and WT mice; N=9 samples for WT, APPNLGF and APPNLGFxPS2FAD 9-month-old mice. Statistical analysis was performed using one-way ANOVA.

**Supplementary 11.**
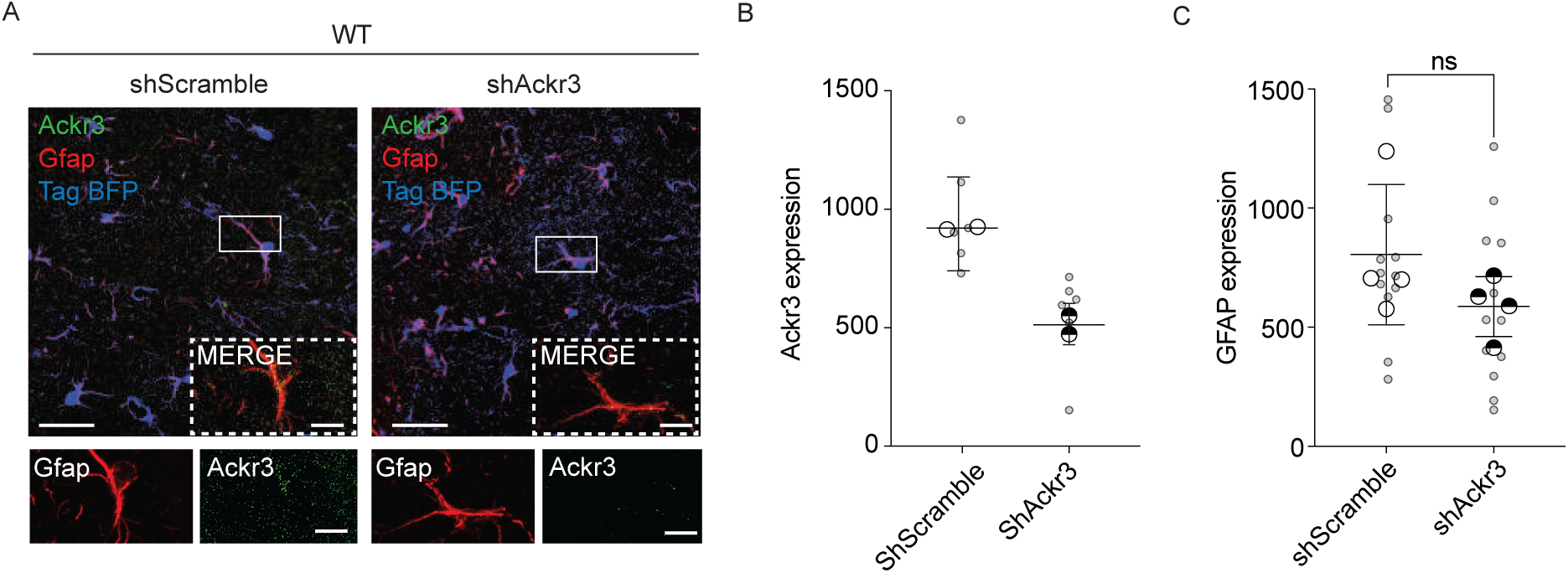
AAV expression in WT and 5xFAD mice. **A.** Representative confocal z-stack images for ACKR3 (green), GFAP (red) in CA1 hippocampus of WT mice injected with AAV9-GfaABC1D-tagBFP-shScramble and AVV9-GfaABC1D-tagBFP- ShAckr3. Scale bar 50 μm, inset 20 μm. **B.** Quantitative analysis of astrocytic Ackr3 expression in the hippocampus of WT mice injected with AAV9-GfaABC1D-tagBFP-shScramble and AAV9- GfaABC1D-tagBFP-ShAckr3; N=2 samples for both shScramble and ShAckr3 4-month-old injected mice. **C.** Quantitative analysis of astrocytic GFAP expression in the hippocampus of WT mice injected with AAV9-GfaABC1D-tagBFP-shScramble and AAV9-GfaABC1D-tagBFP-ShAckr3.; N=4 samples for both shScramble and ShAckr3 4-month-old injected mice. (ShScramble vs. ShAckr3 p=0.170). Statistical analysis was performed using unpaired t test

**Supplementary 12.**
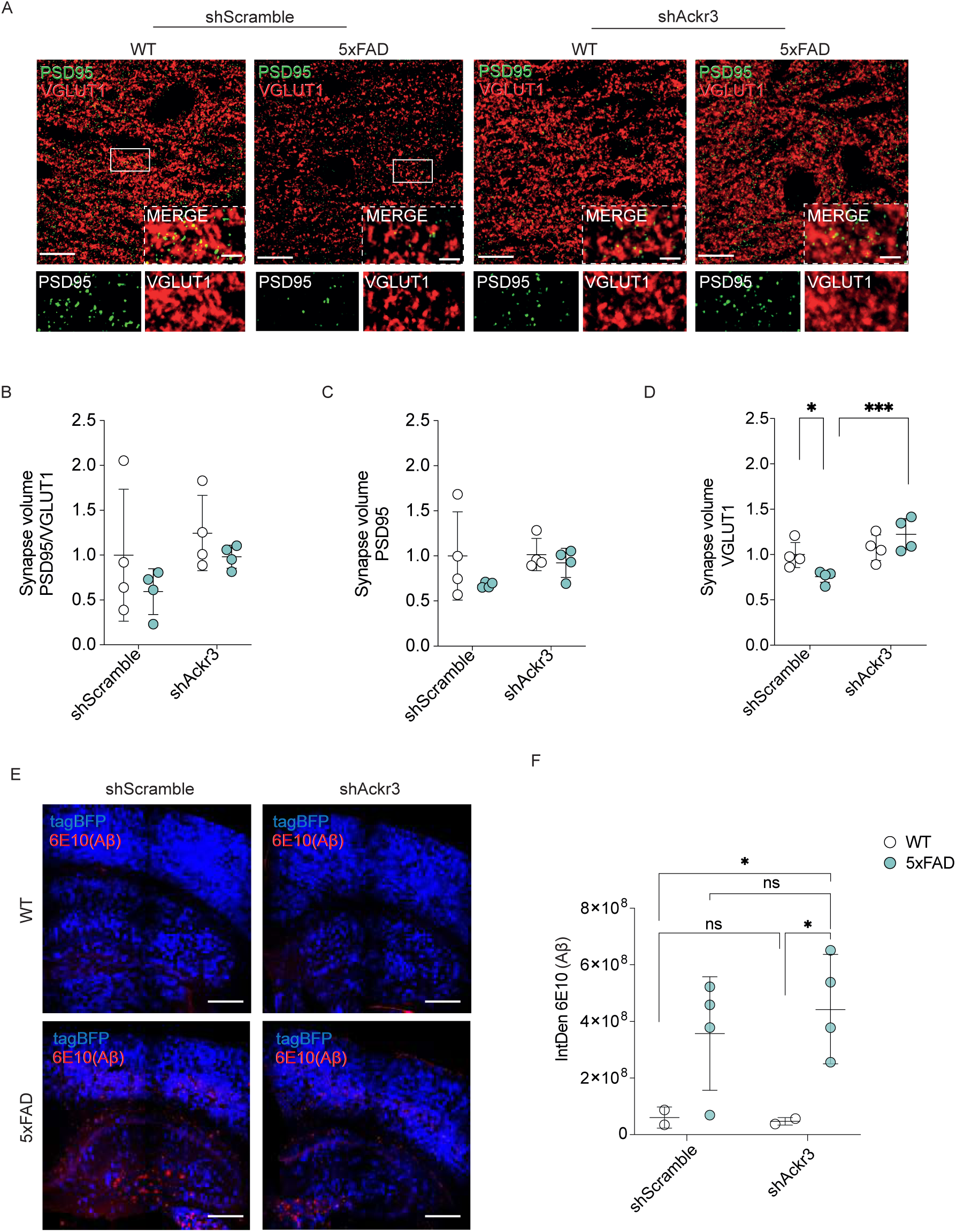
Synapses Functional evaluation and AAV expression in WT and 5xFAD mice. **A.** Representative confocal z-stack images of PSD95 (green)-and VGLUT (red)-positive synapses in the hippocampus of 5XFAD mice compared to WT mice injected with AAV9-GfaABC1D-tagBFP- shScramble and AAV9-GfaABC1D-tagBFP-ShAckr3. Scale bar 50μm, inset 20μm. **B.** Quantitative analysis of PSD95-positive synapses co-localizing with VGLUT1-positive synapses in the hippocampus of 5XFAD and WT mice injected with AAV9-GfaABC1D-tagBFP-shScramble and AAV9-GfaABC1D-tagBFP-ShAckr3; Scale bar 20 μm; N=4 samples for both WT and 5XFAD 4- month-old mice. Statistical analysis was performed using Two-Way ANOVA with Tukey’s multiple comparison test. **C.** Quantitative analysis of PSD95-positive synapses in the hippocampus of 5XFAD and WT mice; N=4 samples for both WT and 5XFAD 4-month-old mice. (WT vs. 5XFAD p= 0.182). Statistical analysis was performed using Two-Way ANOVA with Tukey’s multiple comparison test. **D.** Quantitative analysis of VGLUT1-positive synapses in the hippocampus of 5XFAD and WT mice; N=4 samples for both WT and 5XFAD 4-month-old mice. Statistical analysis was performed using Two-Way ANOVA with Tukey’s multiple comparison test. **E.** Representative confocal z-stack images for tagBFP (blue) and 6E10(Aβ) (red) in CA1 hippocampus of WT mice injected with AAV9- GfaABC1D-tagBFP-shScramble and AVV9-GfaABC1D-tagBFP-ShAckr3. Scale bar 50 μm, **F.** Quantitative analysis of 6E10(Aβ) IntDen in the hippocampus of WT and 5XFAD mice injected with AAV9-GfaABC1D-tagBFP-shScramble and AAV9-GfaABC1D-tagBFP-ShAckr3. N=2 samples for WT and N=4 samples for 5XFAD mice. Statistical analysis was performed using Two-Way ANOVA with Tukey’s multiple comparison test.

**Supplementary 13.**
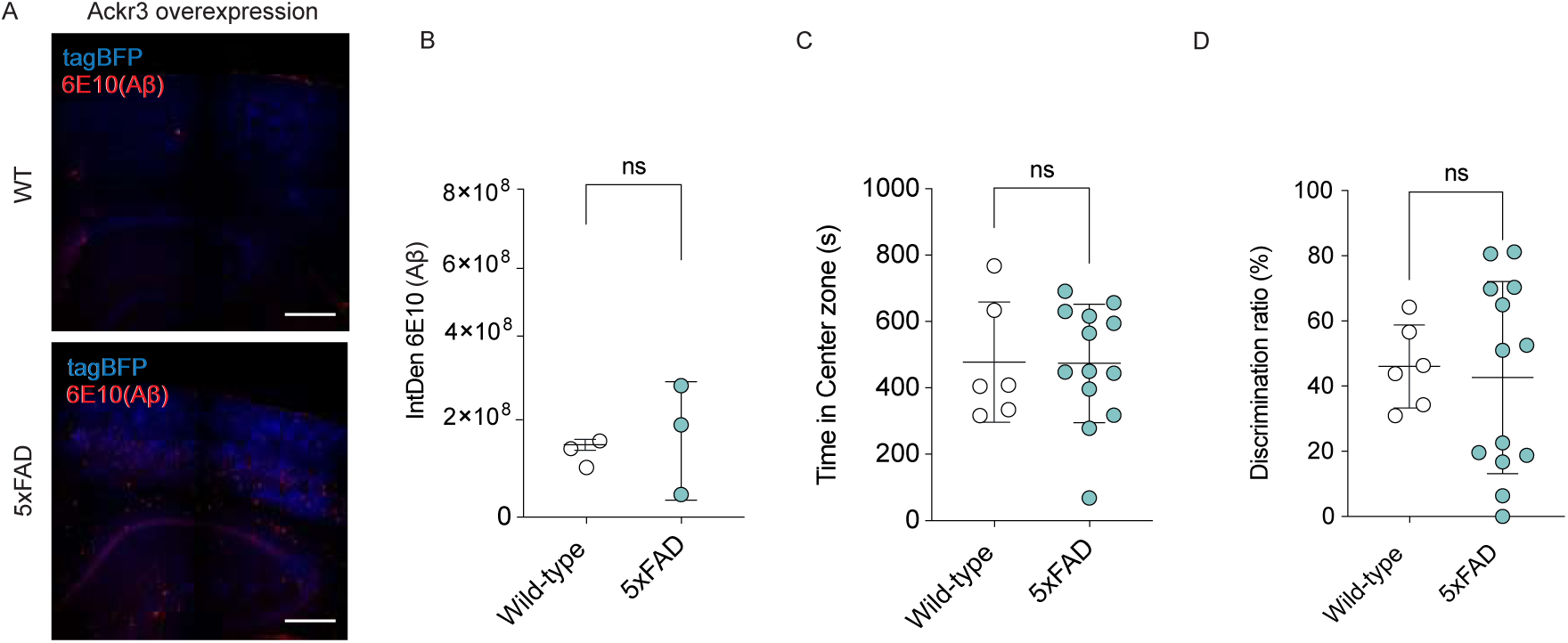
Functional evaluation of Ackr3-overexpression in WT and 5xFAD mice. **A.** Representative confocal z-stack images for tagBFP (blue) and 6E10(Aβ) (red) in CA1 hippocampus of WT and 5XFAD mice injected with PHP.eB-AVV9-GfaABC1D-tagBFP-Control and PHP.eB AVV9-GfaABC1D-tagBFP-Ackr3. Scale bar 50 μm, **B.** Quantitative analysis of 6E10(Aβ) IntDen in the hippocampus of WT and 5XFAD mice injected with5XFAD and WT mice injected with PHP.eB-tagBFP-Ackr3; N=3 samples for WT and N=3 samples for 5XFAD mice; Statistical analysis was performed using unpaired t test. **C.** Open field behavioural analysis showing the 4-month-old 5XFAD and WT mice injected with PHP.eB-tagBFP-Ackr3; N=5 samples for WT and N=13 samples for 5XFAD mice. Statistical analysis was performed using unpaired t test. **D.** Y-maze behavioural analysis showing the 4-month-old 5XFAD and WT mice injected with PHP.eB- tagBFP-Ackr3; N=5 samples for WT and N=13 samples for 5XFAD mice. Statistical analysis was performed using unpaired t test.

## Notes

### Competing Interest Statement

The authors have declared no competing interest.

### Summary of Updates

Manuscript updated, Figure revised and new author added

## REFERENCES

1. Chung, W.S., Allen, N.J., and Eroglu, C. (2015). Astrocytes Control Synapse Formation, Function, and Elimination. Cold Spring Harb Perspect Biol 7, a020370. 10.1101/cshperspect.a020370.

2. Konishi, H., Koizumi, S., and Kiyama, H. (2022). Phagocytic astrocytes: Emerging from the shadows of microglia. Glia 70, 1009–1026. 10.1002/glia.24145.

3. Wilton, D.K., Dissing-Olesen, L., and Stevens, B. (2019). Neuron-Glia Signaling in Synapse Elimination. Annu Rev Neurosci 42, 107–127. 10.1146/annurev-neuro-070918-050306.

4. Chung, W.S., Clarke, L.E., Wang, G.X., Stafford, B.K., Sher, A., Chakraborty, C., Joung, J., Foo, L.C., Thompson, A., Chen, C., et al. (2013). Astrocytes mediate synapse elimination through MEGF10 and MERTK pathways. Nature 504, 394–400. 10.1038/nature12776.

5. Yang, J., Yang, H., Liu, Y., Li, X., Qin, L., Lou, H., Duan, S., and Wang, H. (2016). Astrocytes contribute to synapse elimination via type 2 inositol 1,4,5-trisphosphate receptor-dependent release of ATP. Elife 5, e15043. 10.7554/eLife.15043.

6. Bellesi, M., de Vivo, L., Chini, M., Gilli, F., Tononi, G., and Cirelli, C. (2017). Sleep Loss Promotes Astrocytic Phagocytosis and Microglial Activation in Mouse Cerebral Cortex. J Neurosci 37, 5263– 5273. 10.1523/JNEUROSCI.3981-16.2017.

7. Morizawa, Y.M., Hirayama, Y., Ohno, N., Shibata, S., Shigetomi, E., Sui, Y., Nabekura, J., Sato, K., Okajima, F., Takebayashi, H., et al. (2017). Reactive astrocytes function as phagocytes after brain ischemia via ABCA1-mediated pathway. Nat Commun 8, 28. 10.1038/s41467-017-00037-1.

8. Damisah, E.C., Hill, R.A., Rai, A., Chen, F., Rothlin, C.V., Ghosh, S., and Grutzendler, J. (2020). Astrocytes and microglia play orchestrated roles and respect phagocytic territories during neuronal corpse removal in vivo. Sci Adv 6, eaba3239. 10.1126/sciadv.aba3239.

9. Zhou, T., Li, Y., Li, X., Zeng, F., Rao, Y., He, Y., Wang, Y., Liu, M., Li, D., Xu, Z., et al. (2022). Microglial debris is cleared by astrocytes via C4b-facilitated phagocytosis and degraded via RUBICON-dependent noncanonical autophagy in mice. Nat Commun 13, 6233. 10.1038/s41467-022-33932-3.

10. Konishi, H., Okamoto, T., Hara, Y., Komine, O., Tamada, H., Maeda, M., Osako, F., Kobayashi, M., Nishiyama, A., Kataoka, Y., et al. (2020). Astrocytic phagocytosis is a compensatory mechanism for microglial dysfunction. EMBO J 39, e104464. 10.15252/embj.2020104464.

11. Lee, J.H., Kim, J.Y., Noh, S., Lee, H., Lee, S.Y., Mun, J.Y., Park, H., and Chung, W.S. (2021). Astrocytes phagocytose adult hippocampal synapses for circuit homeostasis. Nature 590, 612–617. 10.1038/s41586-020-03060-3.

12. Dejanovic, B., Wu, T., Tsai, M.C., Graykowski, D., Gandham, V.D., Rose, C.M., Bakalarski, C.E., Ngu, H., Wang, Y., Pandey, S., et al. (2022). Complement C1q-dependent excitatory and inhibitory synapse elimination by astrocytes and microglia in Alzheimer’s disease mouse models. Nat Aging 2, 837–850. 10.1038/s43587-022-00281-1.

13. Morizawa, Y.M., Hirayama, Y., Ohno, N., Shibata, S., Shigetomi, E., Sui, Y., Nabekura, J., Sato, K., Okajima, F., Takebayashi, H., et al. (2017). Author Correction: Reactive astrocytes function as phagocytes after brain ischemia via ABCA1-mediated pathway. Nat Commun 8, 1598. 10.1038/s41467-017-01594-1.

14. Purice, M.D., Speese, S.D., and Logan, M.A. (2016). Delayed glial clearance of degenerating axons in aged Drosophila is due to reduced PI3K/Draper activity. Nat Commun 7, 12871. 10.1038/ncomms12871.

15. Chiu, H., Zou, Y., Suzuki, N., Hsieh, Y.W., Chuang, C.F., Wu, Y.C., and Chang, C. (2018). Engulfing cells promote neuronal regeneration and remove neuronal debris through distinct biochemical functions of CED-1. Nat Commun 9, 4842. 10.1038/s41467-018-07291-x.

16. Iram, T., Ramirez-Ortiz, Z., Byrne, M.H., Coleman, U.A., Kingery, N.D., Means, T.K., Frenkel, D., and El Khoury, J. (2016). Megf10 Is a Receptor for C1Q That Mediates Clearance of Apoptotic Cells by Astrocytes. J Neurosci 36, 5185–5192. 10.1523/JNEUROSCI.3850-15.2016.

17. Tzioras, M., Daniels, M.J.D., Davies, C., Baxter, P., King, D., McKay, S., Varga, B., Popovic, K., Hernandez, M., Stevenson, A.J., et al. (2023). Human astrocytes and microglia show augmented ingestion of synapses in Alzheimer’s disease via MFG-E8. Cell Rep Med 4, 101175. 10.1016/j.xcrm.2023.101175.

18. Vainchtein, I.D., Chin, G., Cho, F.S., Kelley, K.W., Miller, J.G., Chien, E.C., Liddelow, S.A., Nguyen, P.T., Nakao-Inoue, H., Dorman, L.C., et al. (2018). Astrocyte-derived interleukin-33 promotes microglial synapse engulfment and neural circuit development. Science 359, 1269–1273. 10.1126/science.aal3589.

19. Liddelow, S.A., Guttenplan, K.A., Clarke, L.E., Bennett, F.C., Bohlen, C.J., Schirmer, L., Bennett, M.L., Munch, A.E., Chung, W.S., Peterson, T.C., et al. (2017). Neurotoxic reactive astrocytes are induced by activated microglia. Nature 541, 481–487. 10.1038/nature21029.

20. Koch, C., and Engele, J. (2020). Functions of the CXCL12 Receptor ACKR3/CXCR7-What Has Been Perceived and What Has Been Overlooked. Mol Pharmacol 98, 577–585. 10.1124/molpharm.120.000056.

21. Shi, Y., Riese, D.J., 2nd, and Shen, J. (2020). The Role of the CXCL12/CXCR4/CXCR7 Chemokine Axis in Cancer. Front Pharmacol 11, 574667. 10.3389/fphar.2020.574667.

22. Huynh, C., Dingemanse, J., Meyer Zu Schwabedissen, H.E., and Sidharta, P.N. (2020). Relevance of the CXCR4/CXCR7-CXCL12 axis and its effect in pathophysiological conditions. Pharmacol Res 161, 105092. 10.1016/j.phrs.2020.105092.

23. Puchert, M., Pelkner, F., Stein, G., Angelov, D.N., Boltze, J., Wagner, D.C., Odoardi, F., Flugel, A., Streit, W.J., and Engele, J. (2017). Astrocytic expression of the CXCL12 receptor, CXCR7/ACKR3 is a hallmark of the diseased, but not developing CNS. Mol Cell Neurosci 85, 105–118. 10.1016/j.mcn.2017.09.001.

24. Smit, M.J., Schlecht-Louf, G., Neves, M., van den Bor, J., Penela, P., Siderius, M., Bachelerie, F., and Mayor, F., Jr. (2021). The CXCL12/CXCR4/ACKR3 Axis in the Tumor Microenvironment: Signaling, Crosstalk, and Therapeutic Targeting. Annu Rev Pharmacol Toxicol 61, 541–563. 10.1146/annurev-pharmtox-010919-023340.

25. Wurth, R., Tarn, K., Jernigan, D., Fernandez, S.V., Cristofanilli, M., Fatatis, A., and Meucci, O. (2015). A Preclinical Model of Inflammatory Breast Cancer to Study the Involvement of CXCR4 and ACKR3 in the Metastatic Process. Transl Oncol 8, 358–367. 10.1016/j.tranon.2015.07.002.

26. Whittaker, V.P., Michaelson, I.A., and Kirkland, R.J. (1964). The separation of synaptic vesicles from nerve-ending particles (’synaptosomes’). Biochem J 90, 293–303. 10.1042/bj0900293.

27. Byun, Y.G., and Chung, W.S. (2018). A Novel In Vitro Live-imaging Assay of Astrocyte-mediated Phagocytosis Using pH Indicator-conjugated Synaptosomes. J Vis Exp. 10.3791/56647.

28. Lehrman, E.K., Wilton, D.K., Litvina, E.Y., Welsh, C.A., Chang, S.T., Frouin, A., Walker, A.J., Heller, M.D., Umemori, H., Chen, C., and Stevens, B. (2018). CD47 Protects Synapses from Excess Microglia-Mediated Pruning during Development. Neuron 100, 120–134 e126. 10.1016/j.neuron.2018.09.017.

29. Streubel-Gallasch, L., Giusti, V., Sandre, M., Tessari, I., Plotegher, N., Giusto, E., Masato, A., Iovino, L., Battisti, I., Arrigoni, G., et al. (2021). Parkinson’s Disease-Associated LRRK2 Interferes with Astrocyte-Mediated Alpha-Synuclein Clearance. Mol Neurobiol 58, 3119–3140. 10.1007/s12035-021-02327-8.

30. van Rooijen, N., and Hendrikx, E. (2010). Liposomes for specific depletion of macrophages from organs and tissues. Methods Mol Biol 605, 189–203. 10.1007/978-1-60327-360-2_13.

31. Chung, W.S., and Barres, B.A. (2012). The role of glial cells in synapse elimination. Curr Opin Neurobiol 22, 438–445. 10.1016/j.conb.2011.10.003.

32. Wang, J., Zheng, M., Yang, X., Zhou, X., and Zhang, S. (2023). The Role of Cathepsin B in Pathophysiologies of Non-tumor and Tumor tissues: A Systematic Review. J Cancer 14, 2344–2358. 10.7150/jca.86531.

33. Zhang, Y., Chen, K., Sloan, S.A., Bennett, M.L., Scholze, A.R., O’Keeffe, S., Phatnani, H.P., Guarnieri, P., Caneda, C., Ruderisch, N., et al. (2014). An RNA-sequencing transcriptome and splicing database of glia, neurons, and vascular cells of the cerebral cortex. J Neurosci 34, 11929–11947. 10.1523/JNEUROSCI.1860-14.2014.

34. Burns, J.M., Summers, B.C., Wang, Y., Melikian, A., Berahovich, R., Miao, Z., Penfold, M.E., Sunshine, M.J., Littman, D.R., Kuo, C.J., et al. (2006). A novel chemokine receptor for SDF-1 and I-TAC involved in cell survival, cell adhesion, and tumor development. J Exp Med 203, 2201–2213. 10.1084/jem.20052144.

35. Wang, C., Chen, W., and Shen, J. (2018). CXCR7 Targeting and Its Major Disease Relevance. Front Pharmacol 9, 641. 10.3389/fphar.2018.00641.

36. Odemis, V., Boosmann, K., Heinen, A., Kury, P., and Engele, J. (2010). CXCR7 is an active component of SDF-1 signalling in astrocytes and Schwann cells. J Cell Sci 123, 1081–1088. 10.1242/jcs.062810.

37. Singh, A.K., Arya, R.K., Trivedi, A.K., Sanyal, S., Baral, R., Dormond, O., Briscoe, D.M., and Datta, D. (2013). Chemokine receptor trio: CXCR3, CXCR4 and CXCR7 crosstalk via CXCL11 and CXCL12. Cytokine Growth Factor Rev 24, 41–49. 10.1016/j.cytogfr.2012.08.007.

38. Sanchez-Martin, L., Sanchez-Mateos, P., and Cabanas, C. (2013). CXCR7 impact on CXCL12 biology and disease. Trends Mol Med 19, 12–22. 10.1016/j.molmed.2012.10.004.

39. Naumann, U., Cameroni, E., Pruenster, M., Mahabaleshwar, H., Raz, E., Zerwes, H.G., Rot, A., and Thelen, M. (2010). CXCR7 functions as a scavenger for CXCL12 and CXCL11. PLoS One 5, e9175. 10.1371/journal.pone.0009175.

40. Berning, P., Schaefer, C., Clemens, D., Korsching, E., Dirksen, U., and Potratz, J. (2018). The CXCR4 antagonist plerixafor (AMD3100) promotes proliferation of Ewing sarcoma cell lines in vitro and activates receptor tyrosine kinase signaling. Cell Commun Signal 16, 21. 10.1186/s12964-018-0233-2.

41. Pontejo, S.M., and Murphy, P.M. (2021). Chemokines act as phosphatidylserine-bound "find-me" signals in apoptotic cell clearance. PLoS Biol 19, e3001259. 10.1371/journal.pbio.3001259.

42. Faris, H., Almasieh, M., and Levin, L.A. (2021). Axonal degeneration induces distinct patterns of phosphatidylserine and phosphatidylethanolamine externalization. Cell Death Discov 7, 247. 10.1038/s41420-021-00641-7.

43. Hankins, H.M., Baldridge, R.D., Xu, P., and Graham, T.R. (2015). Role of flippases, scramblases and transfer proteins in phosphatidylserine subcellular distribution. Traffic 16, 35–47. 10.1111/tra.12233.

44. Takeda, M., Yamagami, K., and Tanaka, K. (2014). Role of phosphatidylserine in phospholipid flippase-mediated vesicle transport in Saccharomyces cerevisiae. Eukaryot Cell 13, 363–375. 10.1128/EC.00279-13.

45. Segawa, K., Kurata, S., Yanagihashi, Y., Brummelkamp, T.R., Matsuda, F., and Nagata, S. (2014). Caspase-mediated cleavage of phospholipid flippase for apoptotic phosphatidylserine exposure. Science 344, 1164–1168. 10.1126/science.1252809.

46. Grifell-Junyent, M., Baum, J.F., Valimets, S., Herrmann, A., Paulusma, C.C., Lopez-Marques, R.L., and Gunther Pomorski, T. (2022). CDC50A is required for aminophospholipid transport and cell fusion in mouse C2C12 myoblasts. J Cell Sci 135. 10.1242/jcs.258649.

47. Kim, Y.E., Chen, J., Langen, R., and Chan, J.R. (2010). Monitoring apoptosis and neuronal degeneration by real-time detection of phosphatidylserine externalization using a polarity-sensitive indicator of viability and apoptosis. Nat Protoc 5, 1396–1405. 10.1038/nprot.2010.101.

48. Kim, Y.E., Chen, J., Chan, J.R., and Langen, R. (2010). Engineering a polarity-sensitive biosensor for time-lapse imaging of apoptotic processes and degeneration. Nat Methods 7, 67–73. 10.1038/nmeth.1405.

49. Abe, P., Mueller, W., Schutz, D., MacKay, F., Thelen, M., Zhang, P., and Stumm, R. (2014). CXCR7 prevents excessive CXCL12-mediated downregulation of CXCR4 in migrating cortical interneurons. Development 141, 1857–1863. 10.1242/dev.104224.

50. Iwamoto, K., Hayakawa, T., Murate, M., Makino, A., Ito, K., Fujisawa, T., and Kobayashi, T. (2007). Curvature-dependent recognition of ethanolamine phospholipids by duramycin and cinnamycin. Biophys J 93, 1608–1619. 10.1529/biophysj.106.101584.

51. Hayashi, F., Nagashima, K., Terui, Y., Kawamura, Y., Matsumoto, K., and Itazaki, H. (1990). The structure of PA48009: the revised structure of duramycin. J Antibiot (Tokyo) 43, 1421–1430. 10.7164/antibiotics.43.1421.

52. Scheff, S.W., Price, D.A., Schmitt, F.A., and Mufson, E.J. (2006). Hippocampal synaptic loss in early Alzheimer’s disease and mild cognitive impairment. Neurobiol Aging 27, 1372–1384. 10.1016/j.neurobiolaging.2005.09.012.

53. Hong, S., Beja-Glasser, V.F., Nfonoyim, B.M., Frouin, A., Li, S., Ramakrishnan, S., Merry, K.M., Shi, Q., Rosenthal, A., Barres, B.A., et al. (2016). Complement and microglia mediate early synapse loss in Alzheimer mouse models. Science 352, 712–716. 10.1126/science.aad8373.

54. Jung, H., Lee, S.Y., Lim, S., Choi, H.R., Choi, Y., Kim, M., Kim, S., Lee, Y., Han, K.H., Chung, W.S., and Kim, C.H. (2022). Anti-inflammatory clearance of amyloid-beta by a chimeric Gas6 fusion protein. Nat Med 28, 1802–1812. 10.1038/s41591-022-01926-9.

55. Shin, H.J., Kim, I.S., Choi, S.G., Lee, K., Park, H., Shin, J., Kim, D., Beom, J., Yi, Y.Y., Gupta, D.P., et al. (2024). Rejuvenating aged microglia by p16(ink4a)-siRNA-loaded nanoparticles increases amyloid-beta clearance in animal models of Alzheimer’s disease. Mol Neurodegener 19, 25. 10.1186/s13024-024-00715-x.

56. Saito, T., Matsuba, Y., Mihira, N., Takano, J., Nilsson, P., Itohara, S., Iwata, N., and Saido, T.C. (2014). Single App knock-in mouse models of Alzheimer’s disease. Nat Neurosci 17, 661–663. 10.1038/nn.3697.

57. Perdok, A., Van Acker, Z.P., Vrancx, C., Sannerud, R., Vorsters, I., Verrengia, A., Callaerts-Vegh, Z., Creemers, E., Gutierrez Fernandez, S., D’Hauw, B., et al. (2024). Altered expression of Presenilin2 impacts endolysosomal homeostasis and synapse function in Alzheimer’s disease-relevant brain circuits. Nat Commun 15, 10412. 10.1038/s41467-024-54777-y.

58. Shen, H., Shi, X.J., Qi, L., Wang, C., Mamtilahun, M., Zhang, Z.J., Chung, W.S., Yang, G.Y., and Tang, Y.H. (2023). Microglia and astrocytes mediate synapse engulfment in a MER tyrosine kinase-dependent manner after traumatic brain injury. Neural Regen Res 18, 1770–1776. 10.4103/1673-5374.363187.

59. Byun, Y.G., Kim, N.S., Kim, G., Jeon, Y.S., Choi, J.B., Park, C.W., Kim, K., Jang, H., Kim, J., Kim, E., et al. (2023). Stress induces behavioral abnormalities by increasing expression of phagocytic receptor MERTK in astrocytes to promote synapse phagocytosis. Immunity 56, 2105–2120 e2113. 10.1016/j.immuni.2023.07.005.

60. Shi, Q., Chowdhury, S., Ma, R., Le, K.X., Hong, S., Caldarone, B.J., Stevens, B., and Lemere, C.A. (2017). Complement C3 deficiency protects against neurodegeneration in aged plaque-rich APP/PS1 mice. Sci Transl Med 9. 10.1126/scitranslmed.aaf6295.

61. Rajendran, L., and Paolicelli, R.C. (2018). Microglia-Mediated Synapse Loss in Alzheimer’s Disease. J Neurosci 38, 2911–2919. 10.1523/JNEUROSCI.1136-17.2017.

62. Ganesh, A., Choudhury, W., and Coutellier, L. (2024). Early spatial recognition memory deficits in 5XFAD female mice are associated with disruption of prefrontal parvalbumin neurons. Brain Res 1841, 149122. 10.1016/j.brainres.2024.149122.

63. Padua, M.S., Guil-Guerrero, J.L., and Lopes, P.A. (2024). Behaviour Hallmarks in Alzheimer’s Disease 5xFAD Mouse Model. Int J Mol Sci 25. 10.3390/ijms25126766.

64. Choi, M., Jang, H.S., Son, T., Kim, D., Youn, Y.J., Hwang, G.B., Choi, Y.P., and Jeong, Y.H. (2023). Effect Sizes of Cognitive and Locomotive Behavior Tests in the 5XFAD-J Mouse Model of Alzheimer’s Disease. Int J Mol Sci 24. 10.3390/ijms242015064.

65. Zieger, K., Cao, C., and Engele, J. (2024). Evaluating CXCL12 for Effects on Reactive Gene Expression in Primary Astrocytes. J Mol Neurosci 74, 57. 10.1007/s12031-024-02231-5.

66. Li, T., Liu, T., Chen, X., Li, L., Feng, M., Zhang, Y., Wan, L., Zhang, C., and Yao, W. (2020). Microglia induce the transformation of A1/A2 reactive astrocytes via the CXCR7/PI3K/Akt pathway in chronic post-surgical pain. J Neuroinflammation 17, 211. 10.1186/s12974-020-01891-5.

67. Schonemeier, B., Kolodziej, A., Schulz, S., Jacobs, S., Hoellt, V., and Stumm, R. (2008). Regional and cellular localization of the CXCl12/SDF-1 chemokine receptor CXCR7 in the developing and adult rat brain. J Comp Neurol 510, 207–220. 10.1002/cne.21780.

68. Balabanian, K., Lagane, B., Infantino, S., Chow, K.Y., Harriague, J., Moepps, B., Arenzana-Seisdedos, F., Thelen, M., and Bachelerie, F. (2005). The chemokine SDF-1/CXCL12 binds to and signals through the orphan receptor RDC1 in T lymphocytes. J Biol Chem 280, 35760–35766. 10.1074/jbc.M508234200.

69. Sanchez-Alcaniz, J.A., Haege, S., Mueller, W., Pla, R., Mackay, F., Schulz, S., Lopez-Bendito, G., Stumm, R., and Marin, O. (2011). Cxcr7 controls neuronal migration by regulating chemokine responsiveness. Neuron 69, 77–90. 10.1016/j.neuron.2010.12.006.

70. Ehrlich, A.T., Semache, M., Couvineau, P., Wojcik, S., Kobayashi, H., Thelen, M., Gross, F., Hogue, M., Le Gouill, C., Darcq, E., et al. (2021). Ackr3-Venus knock-in mouse lights up brain vasculature. Mol Brain 14, 151. 10.1186/s13041-021-00862-y.

71. Dietz, A., Senf, K., and Neuhaus, E.M. (2024). ACKR3 in olfactory glia cells shapes the immune defense of the olfactory mucosa. Glia 72, 1183–1200. 10.1002/glia.24527.

72. Fumagalli, A., Heuninck, J., Pizzoccaro, A., Moutin, E., Koenen, J., Seveno, M., Durroux, T., Junier, M.P., Schlecht-Louf, G., Bachelerie, F., et al. (2020). The atypical chemokine receptor 3 interacts with Connexin 43 inhibiting astrocytic gap junctional intercellular communication. Nat Commun 11, 4855. 10.1038/s41467-020-18634-y.

73. Salazar, N., Carlson, J.C., Huang, K., Zheng, Y., Oderup, C., Gross, J., Jang, A.D., Burke, T.M., Lewen, S., Scholz, A., et al. (2018). A Chimeric Antibody against ACKR3/CXCR7 in Combination with TMZ Activates Immune Responses and Extends Survival in Mouse GBM Models. Mol Ther 26, 1354– 1365. 10.1016/j.ymthe.2018.02.030.

74. Xu, S., Tang, J., Wang, C., Liu, J., Fu, Y., and Luo, Y. (2019). CXCR7 promotes melanoma tumorigenesis via Src kinase signaling. Cell Death Dis 10, 191. 10.1038/s41419-019-1442-3.

75. Liu, Y., Carson-Walter, E., and Walter, K.A. (2014). Chemokine receptor CXCR7 is a functional receptor for CXCL12 in brain endothelial cells. PLoS One 9, e103938. 10.1371/journal.pone.0103938.

76. Ma, W., Liu, Y., Ellison, N., and Shen, J. (2013). Induction of C-X-C chemokine receptor type 7 (CXCR7) switches stromal cell-derived factor-1 (SDF-1) signaling and phagocytic activity in macrophages linked to atherosclerosis. J Biol Chem 288, 15481–15494. 10.1074/jbc.M112.445510.

77. Yan, Y., Su, J., and Zhang, Z. (2022). The CXCL12/CXCR4/ACKR3 Response Axis in Chronic Neurodegenerative Disorders of the Central Nervous System: Therapeutic Target and Biomarker. Cell Mol Neurobiol 42, 2147–2156. 10.1007/s10571-021-01115-1.

78. Segev, Y., Barrera, I., Ounallah-Saad, H., Wibrand, K., Sporild, I., Livne, A., Rosenberg, T., David, O., Mints, M., Bramham, C.R., and Rosenblum, K. (2015). PKR Inhibition Rescues Memory Deficit and ATF4 Overexpression in ApoE epsilon4 Human Replacement Mice. J Neurosci 35, 12986–12993. 10.1523/JNEUROSCI.5241-14.2015.

79. Wen, L., Bi, D., and Shen, Y. (2024). Complement-mediated synapse loss in Alzheimer’s disease: mechanisms and involvement of risk factors. Trends Neurosci 47, 135–149. 10.1016/j.tins.2023.11.010.

80. Gomez-Arboledas, A., Fonseca, M.I., Kramar, E., Chu, S.H., Schartz, N.D., Selvan, P., Wood, M.A., and Tenner, A.J. (2024). C5aR1 signaling promotes region- and age-dependent synaptic pruning in models of Alzheimer’s disease. Alzheimers Dement 20, 2173–2190. 10.1002/alz.13682.

81. Rostami, J., Mothes, T., Kolahdouzan, M., Eriksson, O., Moslem, M., Bergstrom, J., Ingelsson, M., O’Callaghan, P., Healy, L.M., Falk, A., and Erlandsson, A. (2021). Crosstalk between astrocytes and microglia results in increased degradation of alpha-synuclein and amyloid-beta aggregates. J Neuroinflammation 18, 124. 10.1186/s12974-021-02158-3.

82. Morrone Parfitt, G., Coccia, E., Goldman, C., Whitney, K., Reyes, R., Sarrafha, L., Nam, K.H., Sohail, S., Jones, D.R., Crary, J.F., et al. (2024). Disruption of lysosomal proteolysis in astrocytes facilitates midbrain organoid proteostasis failure in an early-onset Parkinson’s disease model. Nat Commun 15, 447. 10.1038/s41467-024-44732-2.

83. Loov, C., Mitchell, C.H., Simonsson, M., and Erlandsson, A. (2015). Slow degradation in phagocytic astrocytes can be enhanced by lysosomal acidification. Glia 63, 1997–2009. 10.1002/glia.22873.

84. Guo, P., Xiao, X., El-Gohary, Y., Paredes, J., Prasadan, K., Shiota, C., Wiersch, J., Welsh, C., and Gittes, G.K. (2013). A simplified purification method for AAV variant by polyethylene glycol aqueous two-phase partitioning. Bioengineered 4, 103–106. 10.4161/bioe.22293.

85. Challis, R.C., Ravindra Kumar, S., Chan, K.Y., Challis, C., Beadle, K., Jang, M.J., Kim, H.M., Rajendran, P.S., Tompkins, J.D., Shivkumar, K., et al. (2019). Systemic AAV vectors for widespread and targeted gene delivery in rodents. Nat Protoc 14, 379–414. 10.1038/s41596-018-0097-3.

86. Iovino, L., Giusti, V., Pischedda, F., Giusto, E., Plotegher, N., Marte, A., Battisti, I., Di Iacovo, A., Marku, A., Piccoli, G., et al. (2022). Trafficking of the glutamate transporter is impaired in LRRK2- related Parkinson’s disease. Acta Neuropathol 144, 81–106. 10.1007/s00401-022-02437-0.

87. Ramos-Gonzalez, P., Mato, S., Chara, J.C., Verkhratsky, A., Matute, C., and Cavaliere, F. (2021). Astrocytic atrophy as a pathological feature of Parkinson’s disease with LRRK2 mutation. NPJ Parkinsons Dis 7, 31. 10.1038/s41531-021-00175-w.

88. Civiero, L., Cirnaru, M.D., Beilina, A., Rodella, U., Russo, I., Belluzzi, E., Lobbestael, E., Reyniers, L., Hondhamuni, G., Lewis, P.A., et al. (2015). Leucine-rich repeat kinase 2 interacts with p21-activated kinase 6 to control neurite complexity in mammalian brain. J Neurochem 135, 1242–1256. 10.1111/jnc.13369.

89. Westmark, P.R., Westmark, C.J., Jeevananthan, A., and Malter, J.S. (2011). Preparation of synaptoneurosomes from mouse cortex using a discontinuous percoll-sucrose density gradient. J Vis Exp. 10.3791/3196.

90. Hughes, C.S., Moggridge, S., Muller, T., Sorensen, P.H., Morin, G.B., and Krijgsveld, J. (2019). Single-pot, solid-phase-enhanced sample preparation for proteomics experiments. Nat Protoc 14, 68–85. 10.1038/s41596-018-0082-x.

91. Walch, P., Selkrig, J., Knodler, L.A., Rettel, M., Stein, F., Fernandez, K., Vieitez, C., Potel, C.M., Scholzen, K., Geyer, M., et al. (2021). Global mapping of Salmonella enterica-host protein-protein interactions during infection. Cell Host Microbe 29, 1316–1332 e1312. 10.1016/j.chom.2021.06.004.

92. Selkrig, J., Stanifer, M., Mateus, A., Mitosch, K., Barrio-Hernandez, I., Rettel, M., Kim, H., Voogdt, C.G.P., Walch, P., Kee, C., et al. (2021). SARS-CoV-2 infection remodels the host protein thermal stability landscape. Mol Syst Biol 17, e10188. 10.15252/msb.202010188.

93. Maatta, T.A., Rettel, M., Sridharan, S., Helm, D., Kurzawa, N., Stein, F., and Savitski, M.M. (2020). Aggregation and disaggregation features of the human proteome. Mol Syst Biol 16, e9500. 10.15252/msb.20209500.

94. Ashburner, J., Brudfors, M., Bronik, K., and Balbastre, Y. (2019). An algorithm for learning shape and appearance models without annotations. Med Image Anal 55, 197–215. 10.1016/j.media.2019.04.008.

95. Wu, T., Hu, E., Xu, S., Chen, M., Guo, P., Dai, Z., Feng, T., Zhou, L., Tang, W., Zhan, L., et al. (2021). clusterProfiler 4.0: A universal enrichment tool for interpreting omics data. Innovation (Camb) 2, 100141. 10.1016/j.xinn.2021.100141.

96. Yu, G., Wang, L.G., Han, Y., and He, Q.Y. (2012). clusterProfiler: an R package for comparing biological themes among gene clusters. OMICS 16, 284–287. 10.1089/omi.2011.0118.

97. Huber, W., Carey, V.J., Gentleman, R., Anders, S., Carlson, M., Carvalho, B.S., Bravo, H.C., Davis, S., Gatto, L., Girke, T., et al. (2015). Orchestrating high-throughput genomic analysis with Bioconductor. Nat Methods 12, 115–121. 10.1038/nmeth.3252.

98. Civiero, L., Vancraenenbroeck, R., Belluzzi, E., Beilina, A., Lobbestael, E., Reyniers, L., Gao, F., Micetic, I., De Maeyer, M., Bubacco, L., et al. (2012). Biochemical characterization of highly purified leucine-rich repeat kinases 1 and 2 demonstrates formation of homodimers. PLoS One 7, e43472. 10.1371/journal.pone.0043472.

